# First paleoproteomics evidence of *Panicum miliaceum* in human dental calculus revealed through expanded protein database approaches

**DOI:** 10.64898/2026.04.03.716286

**Authors:** Marine Morvan, Giedrė Motuzaitė Matuzevičiūtė

## Abstract

Ancient proteins provide a direct window into past diets by enabling the identification of consumed foods through the analysis of dental calculus. While previous studies have reliably detected animal-derived proteins such as milk, plant-derived proteins remain markedly underrepresented, leaving a significant gap in our understanding of the role of plants in past human diets.

Here, we re-analyze open-access paleoproteomics datasets using an expanded protein database approach. This approach incorporates both reviewed and unreviewed entries, enabling the detection of species-specific protein sequences not validated in commonly used databases. We focus on the proteome of *Panicum miliaceum* and revisit two archaeological dental calculus datasets spanning the Eneolithic to Iron Age from the Pontic–Caspian region and the Levantine coast (*n* = 63 individuals).

We identify 60 unique peptides derived from 60 previously overlooked proteins of *Panicum miliaceum* in 39 individuals. All peptides are taxonomically unique to *Panicum miliaceum* and were confidently assigned using a stringent multi-tier validation strategy. These results provide the first paleoproteomic evidence of its consumption in human dental calculus, thereby revising its chronology and dispersal pathways across Eurasia.

More broadly, this study demonstrates that expanding protein databases beyond current annotations enables the detection of underrepresented plant taxa and provides a generalizable framework for improving plant identification in paleoproteomics.

## 1. Introduction

### 1.1. Dental calculus

Ancient proteins represent a direct window into ancient diets by enabling the identification of consumed foods through the analysis of dental calculus. While previous studies on human dental calculus have reliably detected animal-derived milk proteins, offering clear evidence of dairy consumption [1–3], plant-derived proteins remain significantly underrepresented [4–6]. This limitation largely stems from the protein databases used for taxonomic assignment, in which plant proteomes remain underrepresented and poorly annotated. Compared with animal proteomes, plant proteomes remain substantially less curated, containing relatively few manually reviewed (Swiss-Prot) entries and a predominance of unreviewed (TrEMBL) records. This imbalance reduces the availability of taxonomically informative peptide sequences and limits the identification of plant taxa in human dental calculus.

Yet reconstructing plant consumption is essential for understanding past lifeways. Beyond their dietary contribution, plants provide insights into agricultural practices, cultural interactions, trade networks, and the processes through which societies adapted to changing environments and incorporated new resources. By expanding the detection of plant taxa, paleoproteomics offers the potential to reveal previously unrecognized aspects of crop dispersal, dietary change, and broader cultural transformations associated with the emergence of increasingly interconnected food systems.

Traditionally, the contribution of plants to human diets has been investigated through archaeobotanical approaches. However, the preservation of plant remains is strongly affected by processing and taphonomic factors, including exposure to fire, oxygen availability during charring, and depositional conditions [7]. Plant remains are often preserved only through accidental burning, cooking-related events, or abrupt waterlogging, whereas many economically important plants may disappear from the archaeological record without leaving identifiable traces [8]. Moreover, archaeobotanical evidence generally documents plant use at the site level and cannot determine whether particular plants were consumed by all individuals or only specific segments of a population, while minor or occasional consumption may remain archaeologically invisible.

Stable isotope analysis has also been widely used to investigate past diets and can provide information on broad dietary patterns, such as the relative contributions of C3 and C4 plants [9]. However, isotopic approaches cannot identify plants at the species level and are generally unable to detect seasonal, occasional, or low-level consumption, resulting in an incomplete picture of plant exploitation in past economies.

Previous paleoproteomics studies of dental calculus have largely relied on reviewed plant protein databases (Swiss-Prot) and have reported the identification of a limited number of domesticated species, including *Solanum tuberosum* [4], *Sesamum indicum* [5], *Pisum sativum* [4], *Glycine max* [4], *Arachis hypogaea* [4], *Triticum aestivum* [6], *Hordeum vulgare* [6], and *Avena sativa* [4]. Because Swiss-Prot contains only a fraction of the protein diversity represented in plant proteomes, additional taxa may have remained undetected in archaeological samples.

In this study, we explored the utility of incorporating both reviewed (Swiss-Prot) and unreviewed (TrEMBL) plant protein entries for the analysis of dental calculus, focusing on the proteome of *Panicum miliaceum* (Broomcorn millet). By integrating validated and unreviewed protein entries, this approach extends beyond the limitations of conventional databases by substantially increasing proteome coverage and the availability of taxonomically informative peptide sequences, thereby enhancing the detection of underrepresented plant taxa in archaeological samples.

Modern proteomics and paleoproteomics actively contribute to open science through the deposition of mass spectrometry (MS) raw data in public repositories such as PRIDE [10] and MassIVE [11]. However, approximately 75% of spectra generated in MS experiments remain unidentified [12,13], highlighting a substantial potential for future re-analysis. In the present study, we re-analyzed two paleoproteomics datasets to assess the feasibility of detecting *Panicum miliaceum* proteins in human dental calculus as direct evidence of broomcorn millet consumption, beyond the limits of indirect archaeobotanical evidence. Using *Panicum miliaceum* as a case study, we demonstrate how the re-analysis of publicly available paleoproteomics datasets, combined with the integration of unreviewed plant protein entries, enables the identification of previously overlooked peptides. More broadly, this data-driven approach illustrates how public paleoproteomics repositories can be leveraged to uncover underrepresented plant taxa preserved in dental calculus and to refine reconstructions of past diets, crop dispersal pathways, and human–plant interactions.

### 1.2. Broomcorn millet as focus

Broomcorn millet was domesticated in northern China around 8000 cal. BCE [14,15], subsequently spreading across Eurasia [16–18] and reaching Europe by approximately 1550 cal. BCE as shown from directly dating broomcorn millet macroremains [19,20]. Yet, it has been extensively discussed about the pathways, timing of its arrival and how this crop was integrated into existing agricultural package once it reached Europe [17,19]. Previous research has suggested that the integration of broomcorn millet in the past food systems of present day European population was very sudden [21], while the identification of millet biomarker at the sites much earlier than the radiocarbon dated millet macroremains [22] raises the question about gradual millet integration process.

Analyzing *Panicum miliaceum* proteins directly from human dental calculus holds immense potential for understanding minute broomcorn millet consumption prior to its broader use, in turn overcoming the issues derived from taphonomy of macroremains.

## 2. Methods

### 2.1. Datasets selection

Two publicly available archaeological dental calculus paleoproteomics datasets were re-analyzed from the PRIDE archive [10]: PXD021498 [5] (*Homo sapiens*, Bronze Age – Iron Age, *n* = 14 individuals) and PXD02300 [2] (*Homo sapiens*, Eneolithic – Bronze Age, *n* = 49 individuals). The datasets were re-analyzed to evaluate the detectability of *Panicum miliaceum*-derived peptides in ancient dental calculus.

Two additional archaeological bone proteomics datasets: PXD006256 [23] (*Homo sapiens*, Historical, *n* = 4 individuals) and PXD058447 [24] (*Homo sapiens*, Pleistocene, *n* = 22 individuals) were included as a negative control to evaluate the specificity of *Panicum miliaceum* peptide identification and to assess potential false positive taxonomic assignments arising from minimal and extended reference database searches.

### 2.2. Databases selection

The extended FASTA database, comprising the complete *Panicum miliaceum* proteome (55,861 entries) downloaded from UniProt on 2025-12-28, was restricted to the target species to maximize peptide identification sensitivity. Protein entries were annotated with evidence at the protein level, transcript level, homology, or predicted level. Only a single entry was reviewed (Swiss-Prot), whereas all remaining entries were unreviewed (TrEMBL).

The minimal FASTA database comprised 22 *Panicum miliaceum* proteins annotated with evidence at the protein or transcript level and downloaded from UniProt on 2026-05-15. Of these, one proteins was reviewed (Swiss-Prot), whereas the remaining entries were unreviewed (TrEMBL).

### 2.3. Protein searches

Dental calculus paleoproteomics datasets (PXD021498 and PXD02300) were re-analyzed using Novor Cloud (v1.15.8) [25] with the extended FASTA database. Variable post-translational modifications included deamidation (N, Q), dioxidation (M), oxidation (F, H, M, W), and pyro-glutamate formation from glutamic acid (E) or glutamine (Q). Each dataset was re-analyzed twice, using either automatic enzyme selection or a non-specific digestion setting.

Bones paleoproteomics datasets (PXD006256 and PXD058447) were also re-analyzed using Novor Cloud (v1.15.8) with the extended and minimal FASTA databases. Variable post-translational modifications included deamidation (N, Q), dioxidation (M), oxidation (F, H, M, W), and pyro-glutamate formation from glutamic acid (E) or glutamine (Q). Each dataset was re-analyzed twice, using either automatic enzyme selection or a non-specific digestion setting. A double validation performed using MaxQuant [26] (v. 2.7.0.0) with the extended and minimal FASTA databases. Variable post-translational modifications included deamidation (N, Q), dioxidation (M), oxidation (M), and pyro-glutamate formation from glutamic acid (E) or glutamine (Q). Each dataset was re-analyzed twice, using either trypsin enzyme selection or a non-specific digestion setting.

### 2.4. Data analysis

Precursor and fragment mass tolerances were automatically determined by the software, with a fragment mass error tolerance set to 0.02 Da. Peptide-spectrum matches (PSMs) were filtered to a 1% false discovery rate using a target–decoy approach. PSMs were ranked by score, and a cutoff based on a scoring rank, retaining only preceding target identifications.

### 2.5. Data validation

These resulting peptides were considered *Panicum miliaceum*-specific only when confirmed as unique across the protein-protein BLAST UniProt, NCBI, and Unipept databases, ensuring multi-tier validation of taxonomic specificity with 100% sequence identity and 100% query coverage. Exceptions were made for peptides absent from one of these databases. This workflow provides proof-of-concept that *Panicum miliaceum*-specific peptides can be robustly detected in dental calculus samples.

## 3. Results

Re-analysis of two open-access paleoproteomics datasets from dental calculus (PXD021498, *n* = 14 individuals, and PXD02300, *n* = 49 individuals) using our *Panicum miliaceum*-focused proteomics workflow (Figure 1) resulted in the first detection of unique peptides (Table 1). Conversely, no *Panicum miliaceum* peptides were detected following re-analysis of two additional archaeological bone proteomics datasets (PXD006256, *n* = 4 individuals, and PXD058447, *n* = 22 individuals) included as negative controls.

**Figure 1:**
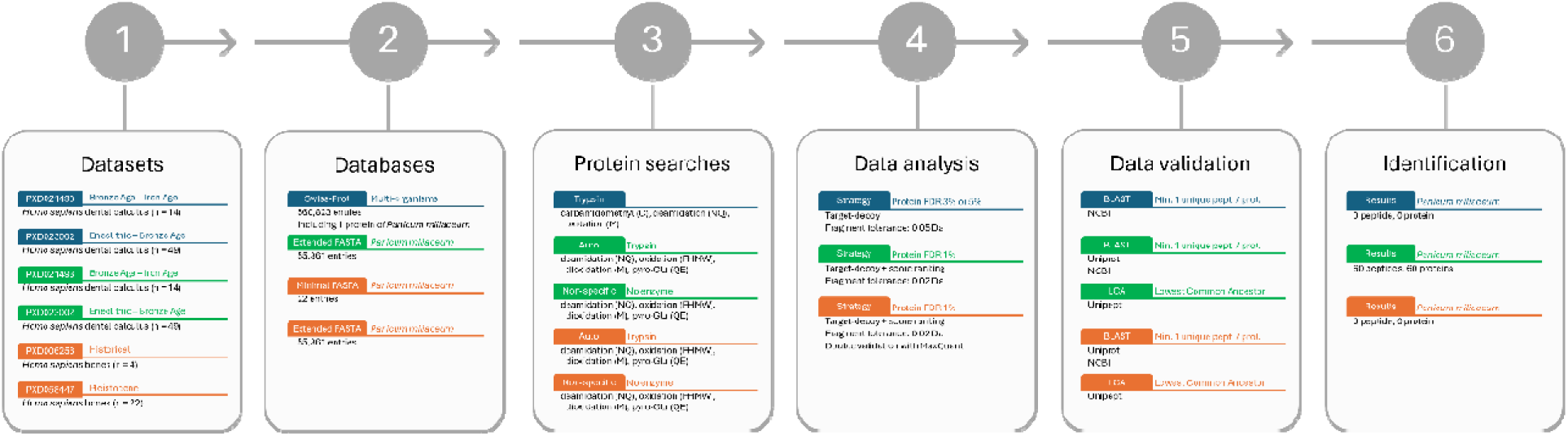
Comparison of proteomics workflow from original studies (blue), this study (green) and negatives controls (orange).

**Table 1:**
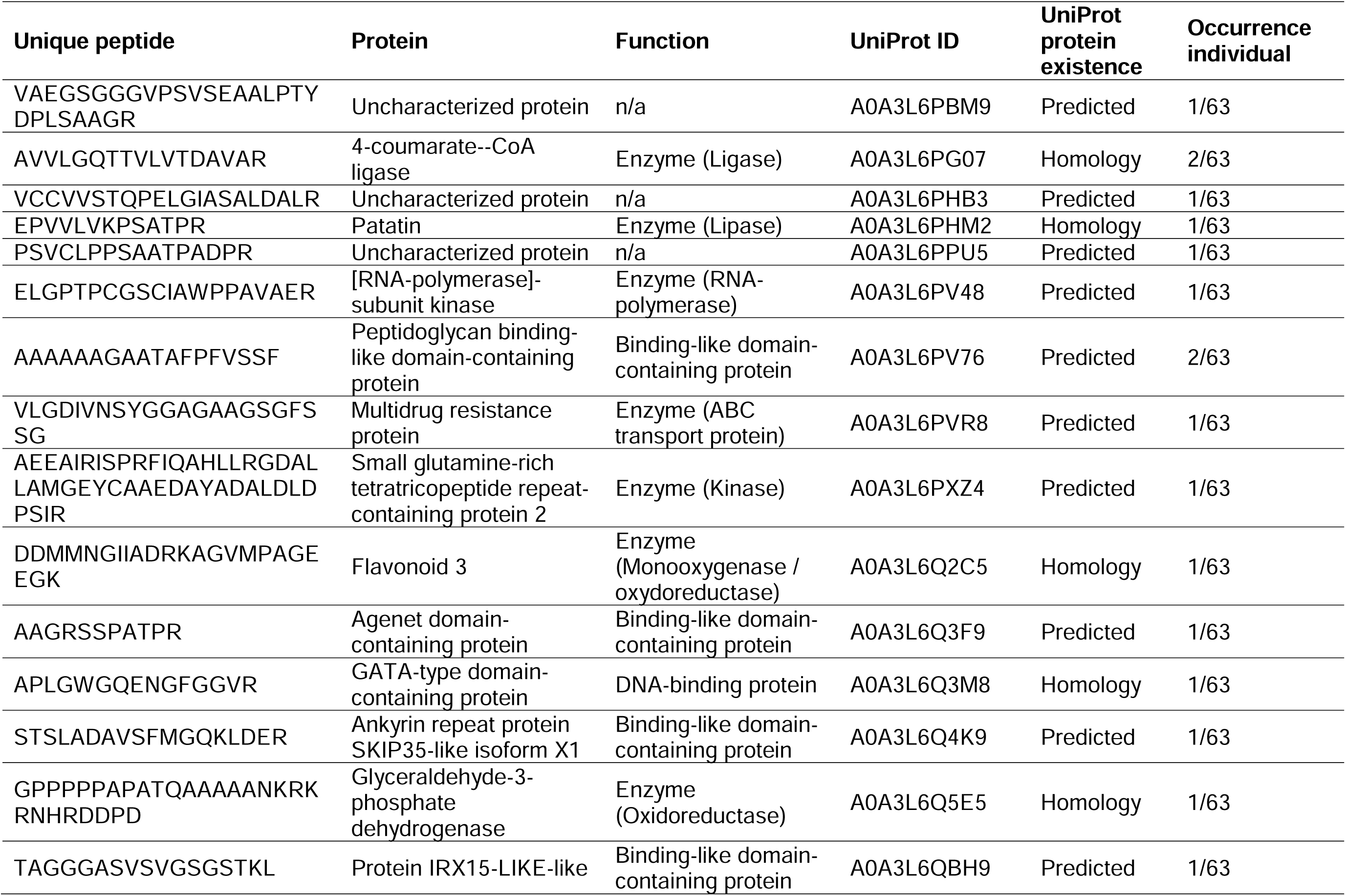
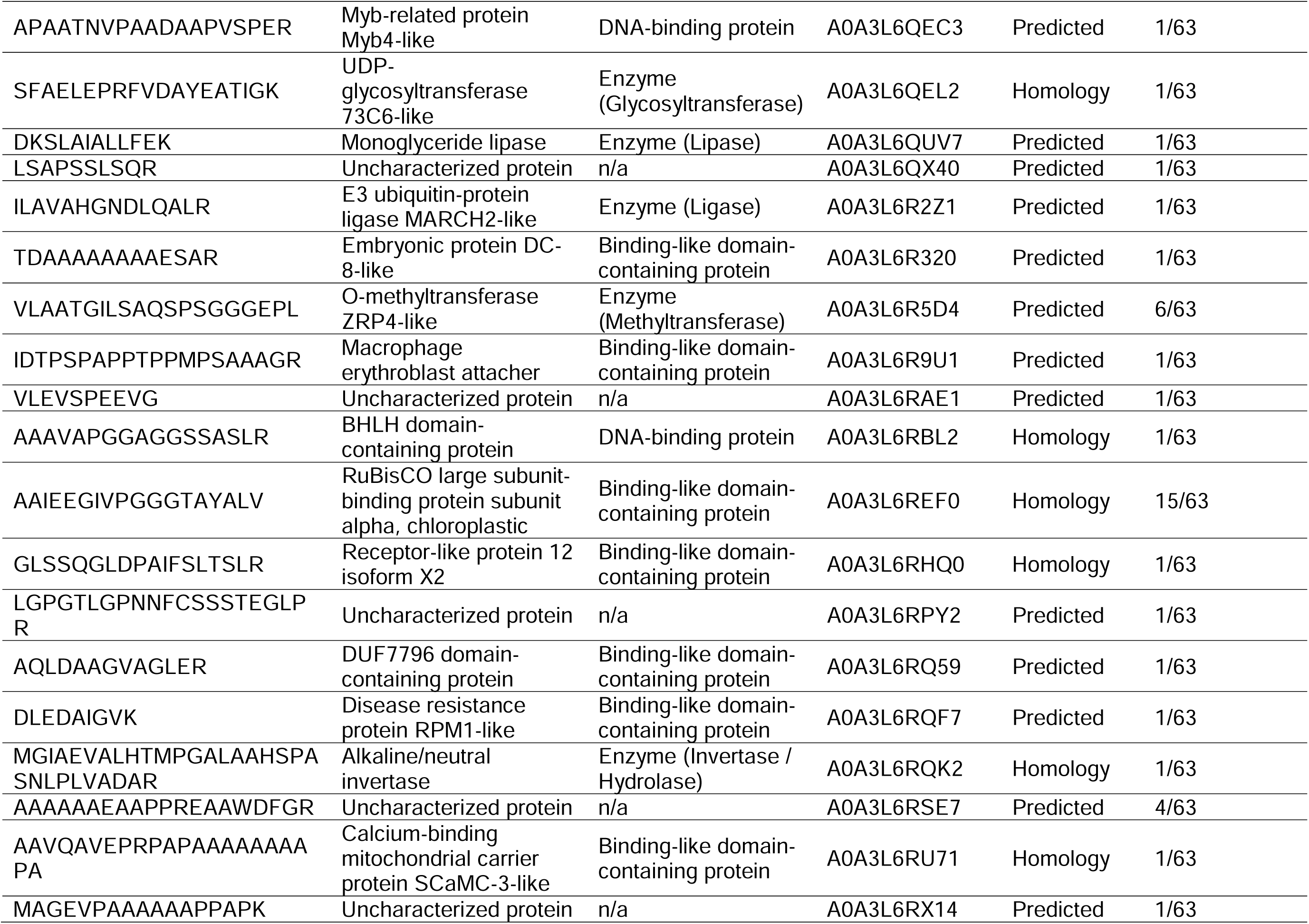
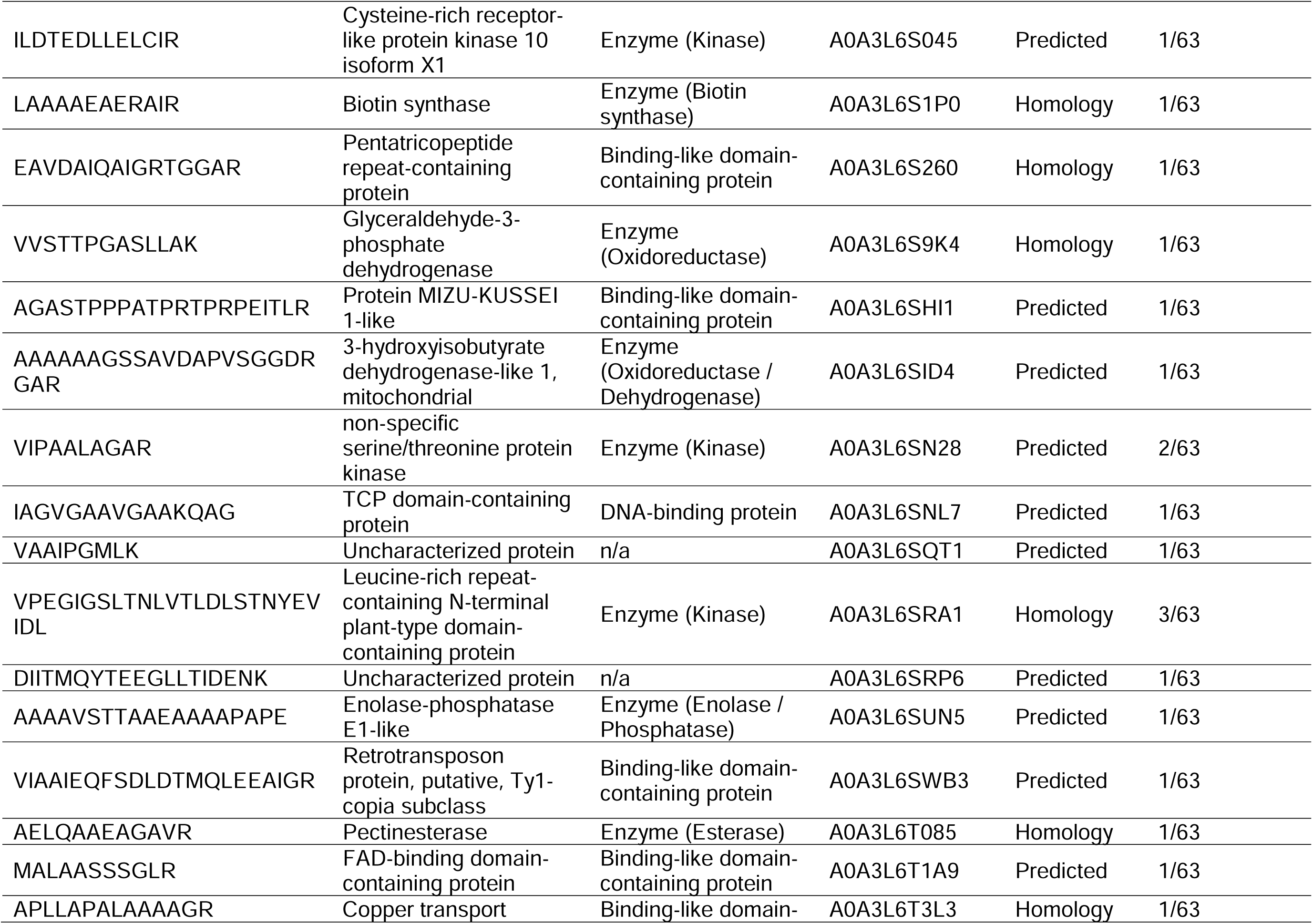
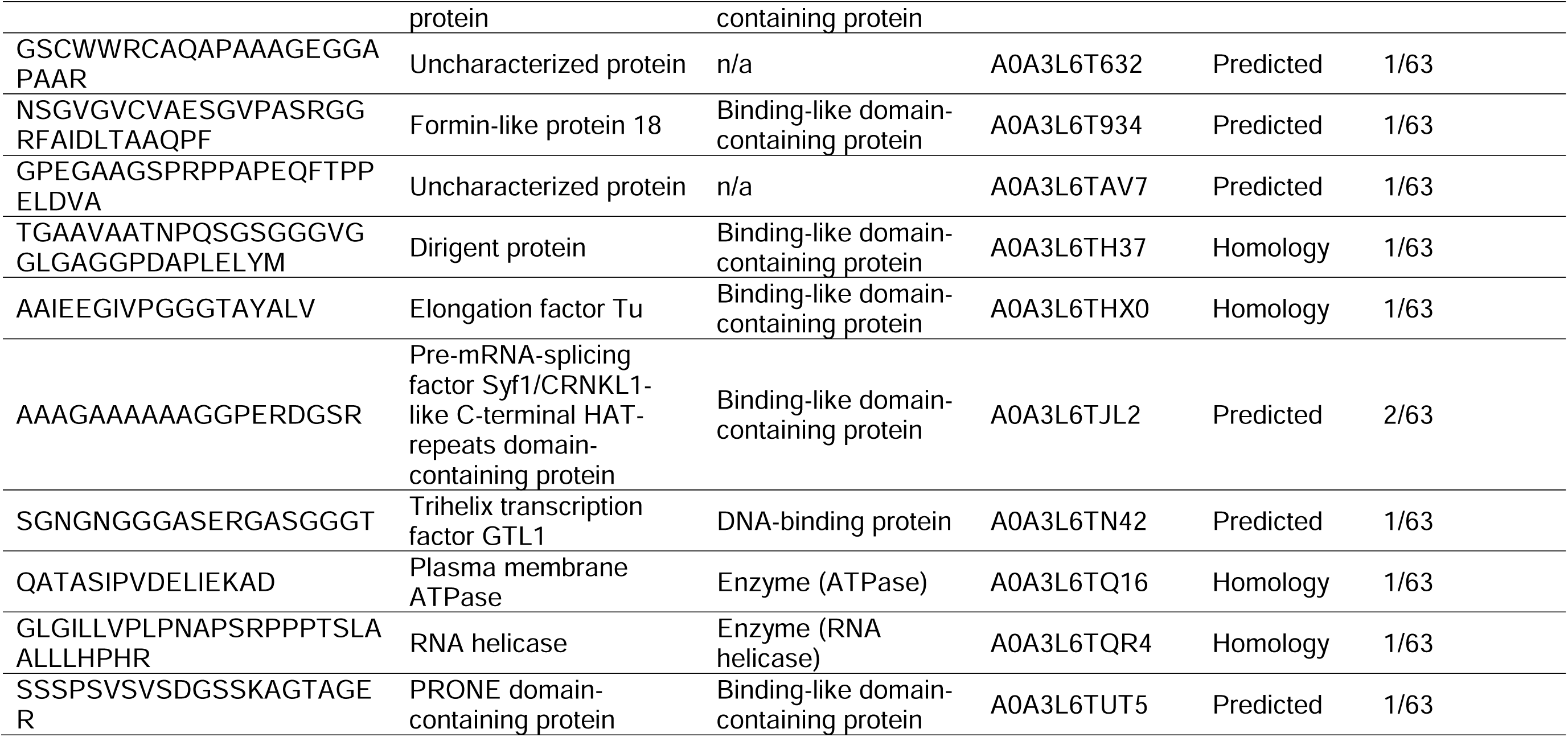
Unique *Panicum miliaceum* peptides and their associated proteins.

In the PXD021498 dataset, Scott *et al.* identified *Poaceae* microremains in the dental calculus of each individual through archaeobotanical analysis, including broomcorn millet in ERA017 individual [5]. However, no *Paicum miliaceum* proteins were reported for any individual using paleoproteomics. In the present study, re-analysis of this dataset showed that 9 of these 14 individuals, spanning the Middle Bronze Age to the Early Iron Age, have clear evidence of broomcorn millet consumption based on paleoproteomics analysis, including the ERA017 individual (Figure 2 on the southern Levantine coast).

**Figure 2:**
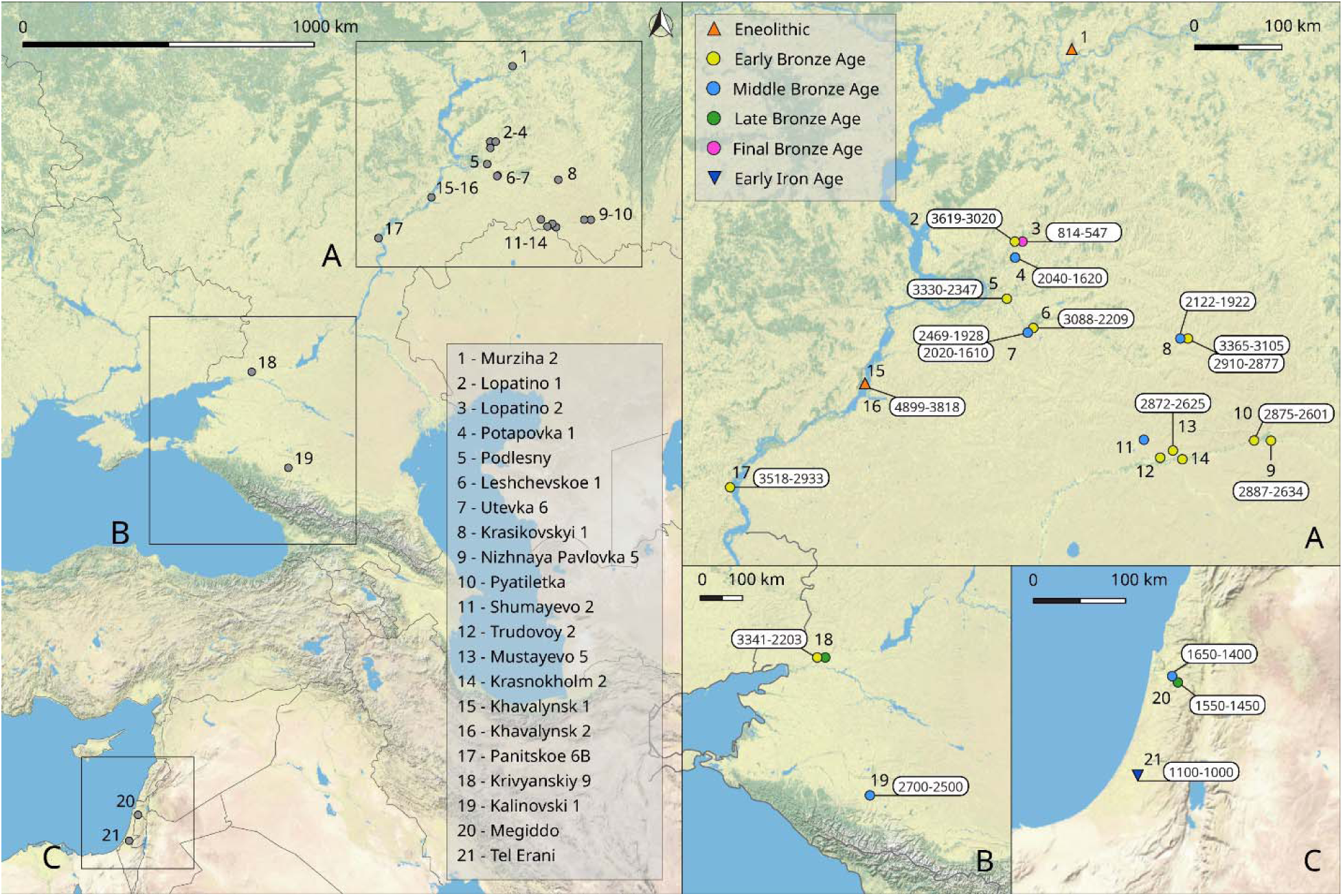
Representation of archeological sites with radiocarbon dates where *Panicum miliaceum* proteins were newly identified from human dental calculus (map by Dovidas Jurkenas).

In the PXD023000 dataset, Wilkin *et al.* focused on the discovery of milk-associated proteins by paleoproteomics [2]. Re-analysis in the present study reveals clear evidence of broomcorn millet consumption in 30 of 49 individuals, spanning the Eneolithic to the Final Bronze Age (Figure 2 in the Pontic–Caspian region), which provides additional information for the reconstruction of the diet of these populations beyond the milk consumption.

The coordinates of sites where proteins of *Panicum miliaceum* were identified in dental calculus, along with their radiocarbon dates as reported in the original publications, are presented in Figure 2. All radiocarbon date ranges are expressed in “cal. BCE”. Individuals from the same site and period with overlapping radiocarbon date ranges were grouped together.

## 4. Discussion

### 4.1. Identification of Panicum miliaceum proteins

In this exploratory study, the primary objective was not to achieve a comprehensive catalogue of peptides in each dental calculus sample, but rather to demonstrate the feasibility of confidently detecting peptides specific to *Panicum miliaceum*, even under highly stringent identification criteria. Accordingly, a highly conservative filtering strategy based on ranking score was employed, prioritizing specificity over sensitivity and minimizing the risk of false-positive identifications. *Panicum miliaceum* is particularly poorly represented in the UniProt proteins database compared to other plant species previously identified in archaeological dental calculus, with only 2 proteins annotated at the protein level, 20 at the transcript level, 22,899 based on homology, and 32,950 predicted entries (Table 2). In shotgun proteomics -as in the original publications-, FASTA files are typically derived from the Swiss-Prot database and therefore include only reviewed proteins. For *Panicum miliaceum*, this corresponds to a single reviewed protein, which would extremely limit the chances of detecting this organism. An additional challenge arises from the limited biological annotation of *Panicum miliaceum* proteins. The sole reviewed protein currently available in Swiss-Prot lacks detailed information regarding its tissue-specific expression and distribution within the plant. Consequently, even if this protein were used as the sole reference in a conventional Swiss-Prot-based search, its recovery from dental calculus would depend on whether it is expressed in plant tissues that were consumed by past populations. Because this information is currently unavailable, failure to detect the reviewed protein cannot be considered evidence for the absence of broomcorn millet consumption. To address these limitations, the present study employed a *Panicum miliaceum* FASTA database containing all 55,861 entries (Swiss-Prot and TrEMBL), maximizing the likelihood of identifying true species-specific peptides. More broadly, the approach presented here may be applicable to other plant species that are currently underrepresented in UniProt and other public protein databases. By leveraging comprehensive FASTA databases that include unreviewed protein entries annotated at the transcript, homology, and predicted levels, future studies may improve the detection of archaeologically relevant taxa that would otherwise remain undetected using conventional Swiss-Prot-based searches. Such developments could substantially expand the range of plants accessible through dental calculus paleoproteomics.

**Table 2:**
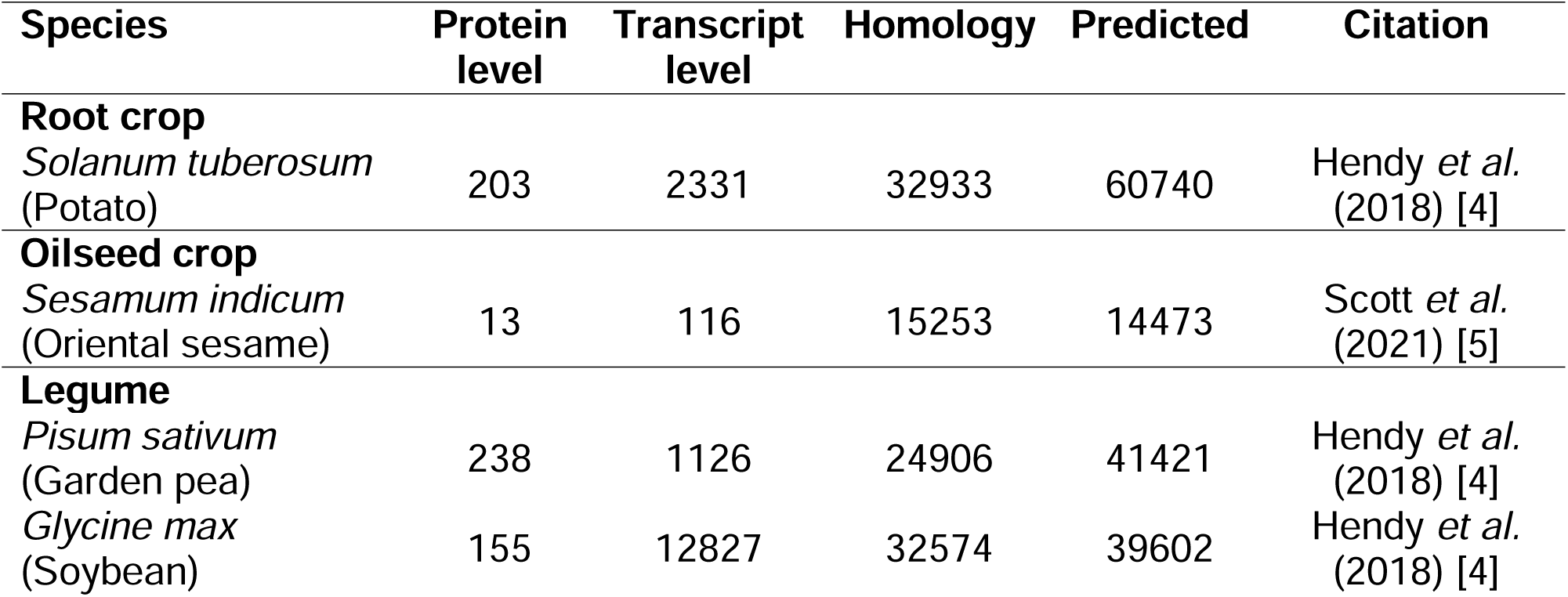
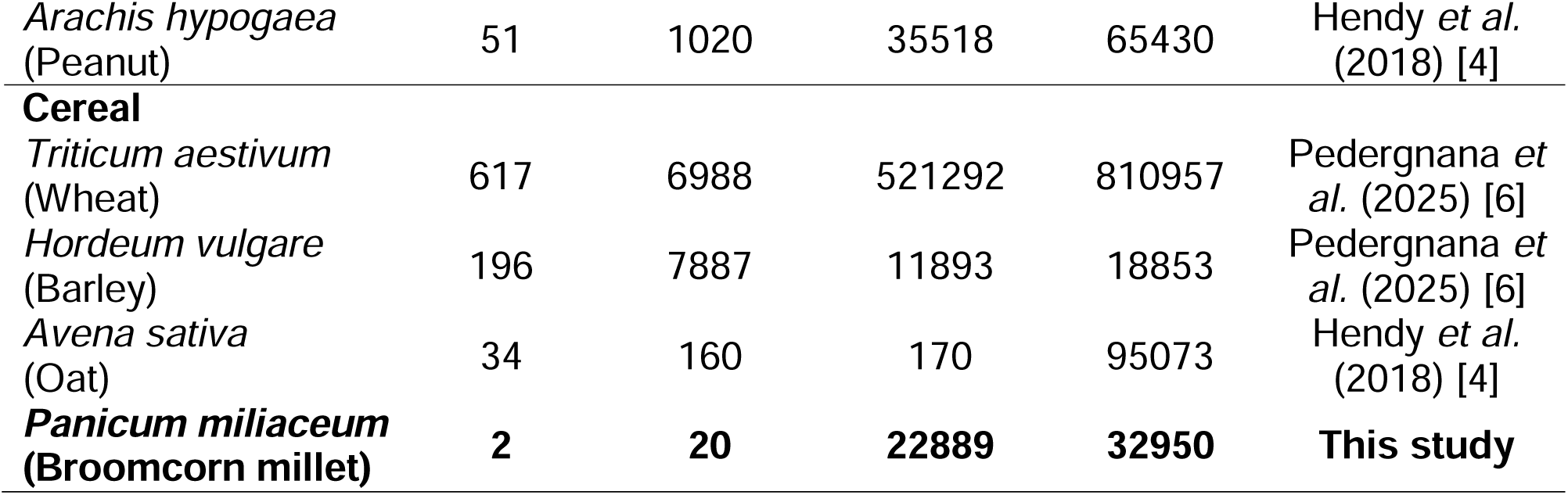
Identified plant species in dental calculus and their corresponding protein description on UniProt on 2026-03-16.

In total, 60 unique peptides corresponding to 60 proteins were confidently identified, with multi-tier BLAST and lowest common ancestor (LCA) validations, for *Panicum miliaceum*. According to annotations in the UniProt protein database, 21 of these proteins are supported at the homology level and 39 at the predicted level, with no entries currently annotated at the protein or transcript evidence level (Table 1). Among them, 12 proteins are further annotated as uncharacterized, lacking assigned protein names and biological functional annotation. The detection of peptides corresponding to these entries therefore provides experimental evidence supporting their expression and highlights the limited characterization currently available for this species proteome. Of the identified peptides, 43 were consistent with tryptic cleavage -authors performed trypsin digestion when preparing their samples-, whereas 17 were identified only under non-enzyme-specific conditions, underscoring the value of combining complementary enzyme-specific and non-specific search strategies to maximize peptide detection in complex and potentially degraded samples. Such complementary search strategies are particularly relevant in aging proteomics and paleoproteomics contexts, where protein degradation can generate peptides that do not conform to enzymatic digestion patterns [27]. Together, these findings highlight the value of proteomics-based approaches for providing experimental evidence that supports computationally predicted proteins and for improving the annotation of poorly characterized plant species. No peptides exhibiting sequence discrepancies relative to the predicted protein sequences contained in the *Panicum miliaceum* FASTA database were detected, thereby supporting the accuracy of the current protein predictions. Although no sequence corrections were required, the present study provides the first peptide-level experimental validation for several proteins that had previously been annotated solely through computational prediction. These results therefore strengthen confidence in the existing proteome annotation and contribute novel experimental evidence for the protein repertoire of this understudied species.

The identification of *Panicum miliaceum*-specific peptides from multiple individuals (Table 1, SI) further strengthens the evidence for the presence of this species in dental calculus samples and demonstrates the robustness and reproducibility of the employed data analysis workflow with a multi-tier validation strategy. The absence of protein overlap between the two re-analyzed datasets is not unexpected, as variations in dietary habits among individuals from different archaeological sites and periods, along with differences in sample preparations, LC-HRMS acquisition methods, and the application of cutoff according to the ranking score, can all influence which peptides are detectable in each sample. The full list of identified peptides, obtained using a 1% FDR without the restrictive cutoff, is provided in the supplementary materials and underscores the potential to identify other *Panicum miliaceum*-specific peptides.

### 4.2. Implications for understanding millet timing and pathways

Along the Levantine coast, isotopic analyses conducted in 2024 at Tell Tweini (northern) did not provide evidence for the consumption of C4 plants, such as broomcorn millet, during the Middle Bronze Age [28]. At the same period, at Megiddo (southern), Scott *et al.* reported no archaeobotanical evidence of broomcorn millet, although *Poaceae* microremains were identified in dental calculus [5]. In contrast, at Tel Erani (southern), *Poaceae* microremains were likewise documented in dental calculus from the Early Iron Age, with *Panicum miliaceum* identified in one individual (ERA017) [5]. The present paleoproteomics study for the first time provides direct molecular evidence for the consumption of broomcorn millet by human populations along the southern Levantine coast from the Middle Bronze Age (Megiddo) to the Early Iron Age (Tel Erani). These results both complement and extend beyond the archaeobotanical data previously reported, thereby refining our understanding of plant exploitation and dietary practices in the region. Future paleoproteomics analyses of dental calculus from northern Levantine coast contexts could be undertaken to further explore the spatial distribution of broomcorn millet consumption. Given that broomcorn millet was introduced from east Asia [29], its presence in northern regions appears plausible for targeted investigation. Such an approach would provide a valuable complementary line of evidence to isotopic studies, particularly in contexts characterized by mixed C3 and C4 plant consumption, where isotopic signals may be difficult to interpret.

More surprisingly, in the Pontic–Caspian region, the re-analysis of dental calculus samples from 28 archaeological sites spanning the Eneolithic to the Middle Bronze Age indicates the consumption of broomcorn millet much earlier than previously thought, pushing broomcorn millet consumption in the western steppe by a few thousand years. Stable isotope studies by Shishlina *et al.* as identified elevated stable carbon isotope values in Early Bronze Age population of this region, yet it was interpreted as wild C4 plant consumption [30]. In addition, specific to broomcorn millet miliacin lipid biomarker was also identifies at the horse herder site in northern Kazakhstan dated to the 4^th^ millennium BCE [31]. The discovery of broomcorn millet proteins in human dental calculus confirms the direct consumption of broomcorn millet by the steppe population and calling to refine the early pathways and chronology of broomcorn millet introduction into Europe. Currently not all individuals that contain *Panicum miliaceum* proteins in their dental calculus were directly dated (Table 3, Figure 2). The oldest *Panicum miliaceum* proteins were found in individuals that have a very broad radiocarbon range stretching from the 5^th^ to 4^th^ millennia that needs to be redated. Most directly dated individuals that contain *Panicum miliaceum* proteins in dental calculus from the northern Caspian steppe were dated to the early 3^rd^ millennium BCE which is by millennia older than the directed dated broomcorn millet grain from the southern Caspian region identified by Huang *et al.* [29].

**Table 3:**
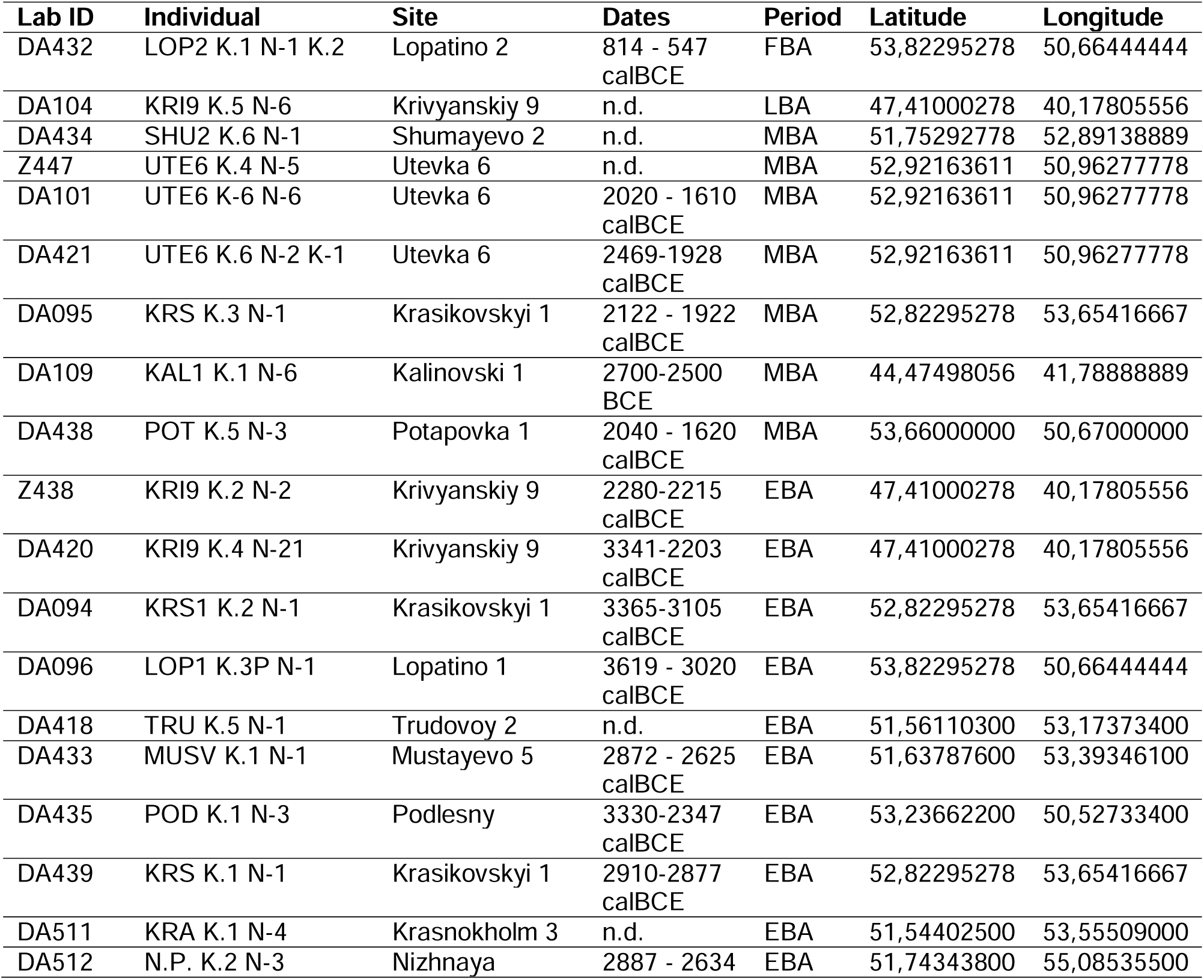
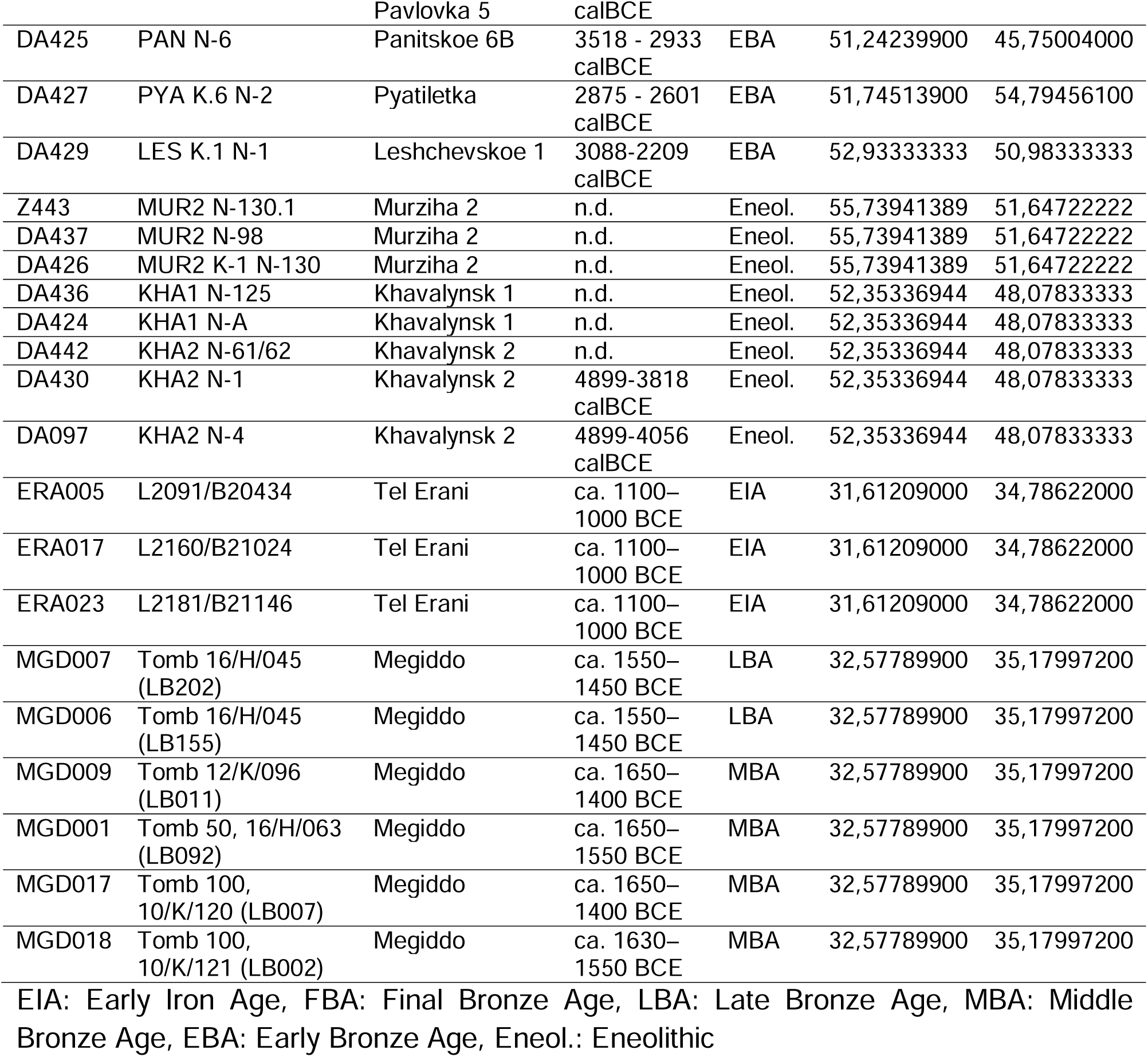
Identified individuals with *Panicum miliaceum* proteins in dental calculus. All data, including human radiocarbon dates for the Pontic–Caspian region and relative date estimates for Levantine sites (Tel Erani and Megiddo), were reported from the original publications.

## 5. Conclusion

This study demonstrates that the re-analysis of open-access paleoproteomics datasets, combined with the integration of expanded protein databases incorporating unreviewed entries, enables the robust detection of plant taxa that have remained underrepresented in reconstructions of past diets. The identification of 60 unique peptides attributable to *Panicum miliaceum* in the dental calculus of 39 individuals provides the first direct molecular evidence for the consumption of broomcorn millet in these archaeological contexts. Importantly, several of these peptides correspond to proteins previously supported only by computational predictions, thereby extending empirical evidence beyond current database annotations and contributing to the validation of poorly characterized proteomes.

The robustness and feasibility of this approach are supported by stringent analytical criteria, including the application of a 1% false discovery rate (FDR), a conservative ranking score threshold, multi-tier BLAST validation, and the requirement for multiple unique peptides derived from distinct proteins. Together, these parameters provide high confidence in the taxonomic assignment and demonstrate that reliable detection of *Panicum miliaceum* is achievable even within complex and degraded paleoproteomic datasets.

These findings challenge established chronologies based solely on archaeobotanical and isotopic evidence by indicating an earlier and potentially more widespread consumption of millet. In particular, the detection of *Panicum miliaceum* proteins in populations from the Pontic-Caspian steppe suggests its consumption as early as the Eneolithic and Early Bronze Age, substantially revising current models of its dispersal across Eurasia. Similarly, evidence from the Levantine coast confirms its consumption from the Middle Bronze Age onward. Collectively, these results refine our understanding of crop diffusion pathways and highlight previously unrecognized patterns of dietary adaptation and plant use.

Beyond the specific case of *Panicum miliaceum*, this work demonstrates that the proposed proteomics framework is broadly applicable to plant taxa whose proteomes remain poorly curated and are largely composed of unreviewed entries. By moving beyond conventional reliance on reviewed databases, this approach provides a generalizable and scalable strategy for detecting underrepresented plant species in archaeological contexts and significantly enhances the taxonomic resolution of paleoproteomic analyses.

More broadly, this study underscores the importance of revisiting existing datasets and fully exploiting the untapped potential of publicly available proteomics repositories. It highlights the central role of open science in enabling new discoveries and advancing the field. Taken together, these results open new avenues for reconstructing past human-plant interactions and offer a powerful framework for investigating ancient food systems, cultural exchanges, and the processes underlying the spread and integration of agricultural resources across Eurasia.

## CRediT authorship contribution statement

Conceptualization: MM & GMM, Data curation: MM, Formal analysis: MM, Funding acquisition: GMM, Investigation: MM, Methodology: MM, Project administration: MM, Resources: MM, Software: MM, Validation: MM, Visualization: MM, Writing – original draft: MM & GMM, Writing – review & editing: MM & GMM

## Declaration of competing interest

The authors declare that they have no known competing financial interests or personal relationships that could have influenced the work reported in this paper.

## Acknowledgements

This research was funded by the European Union with a Consolidator Grant awarded to Giedrė Motuzaitė Matuzevičiūtė (ERC-CoG, MILWAYS, 101087964). Views and opinions expressed are those of the authors only and do not necessarily reflect those of the European Union or the European Research Council Executive Agency. Neither the European Union nor the granting authority can be held responsible for them.

We warmly acknowledged authors for sharing paleoproteomics datasets.

## Notes

### Competing Interest Statement

The authors have declared no competing interest.

### Summary of Updates

We have significantly reworked the main text. The results remain the same.

