## Supplementary materials 1 for "First paleoproteomics evidence of *Panicum miliaceum* in human dental calculus revealed through expanded protein database approaches"

Supplementary material

**ERA005 – Tel Erani – Iron Age**
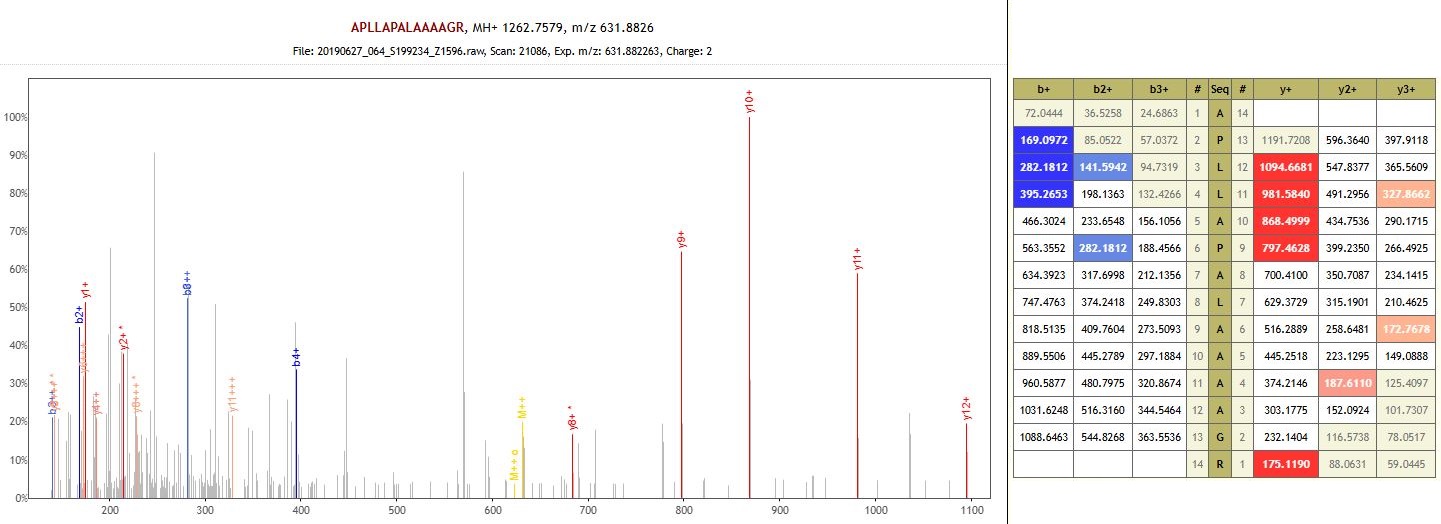

Protein: Copper transport protein (auto-enzyme)

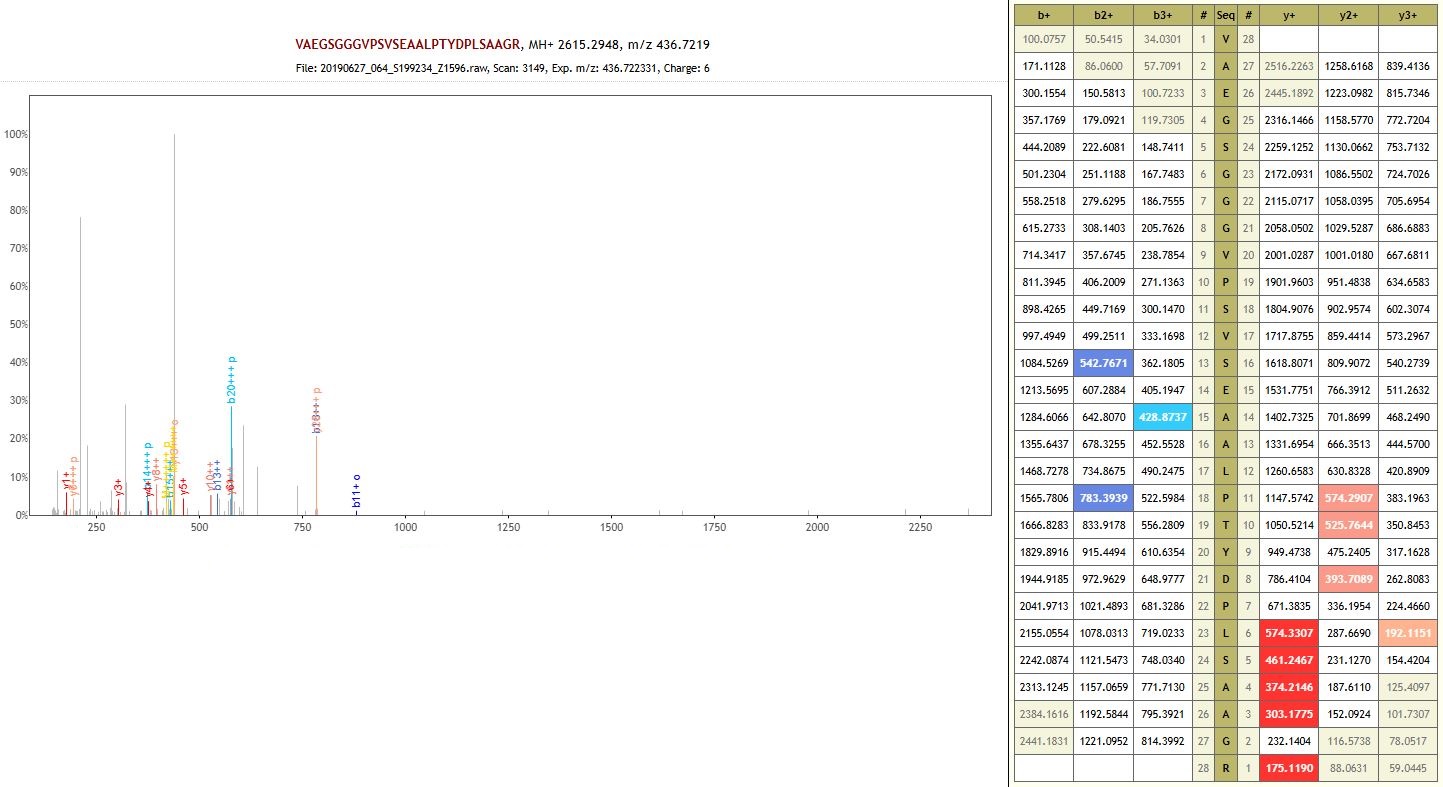
Protein: Uncharacterized protein (auto-enzyme)

**ERA017 – Tel Erani – Iron Age**

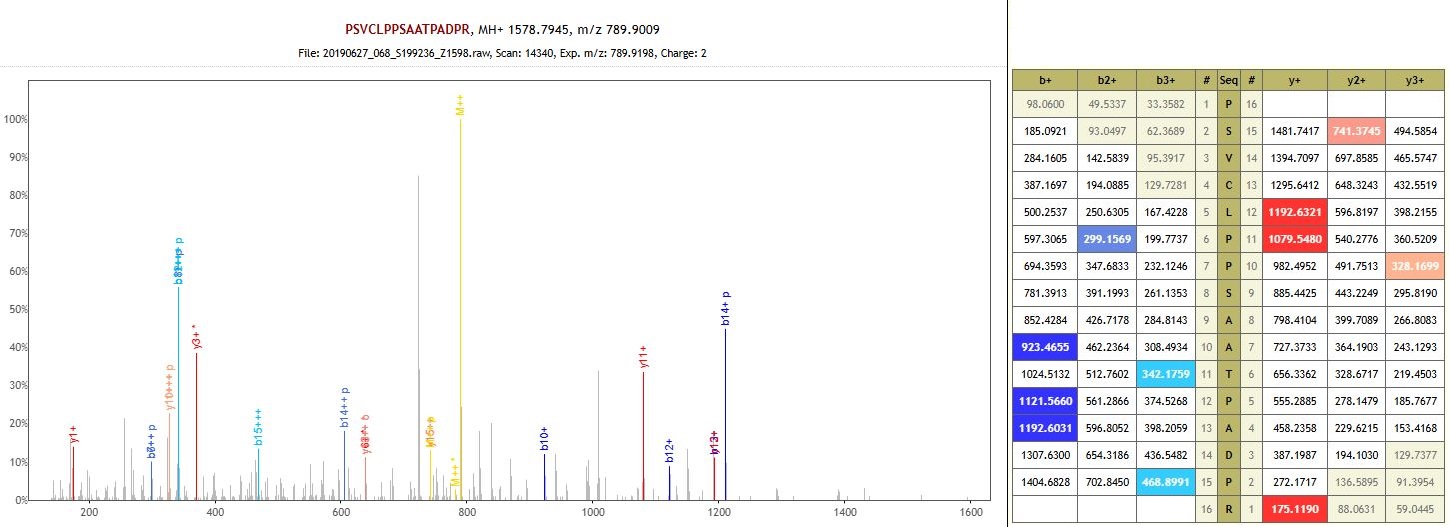

Protein: Uncharacterized protein (auto-enzyme)

**ERA023 – Tel Erani – Iron Age**

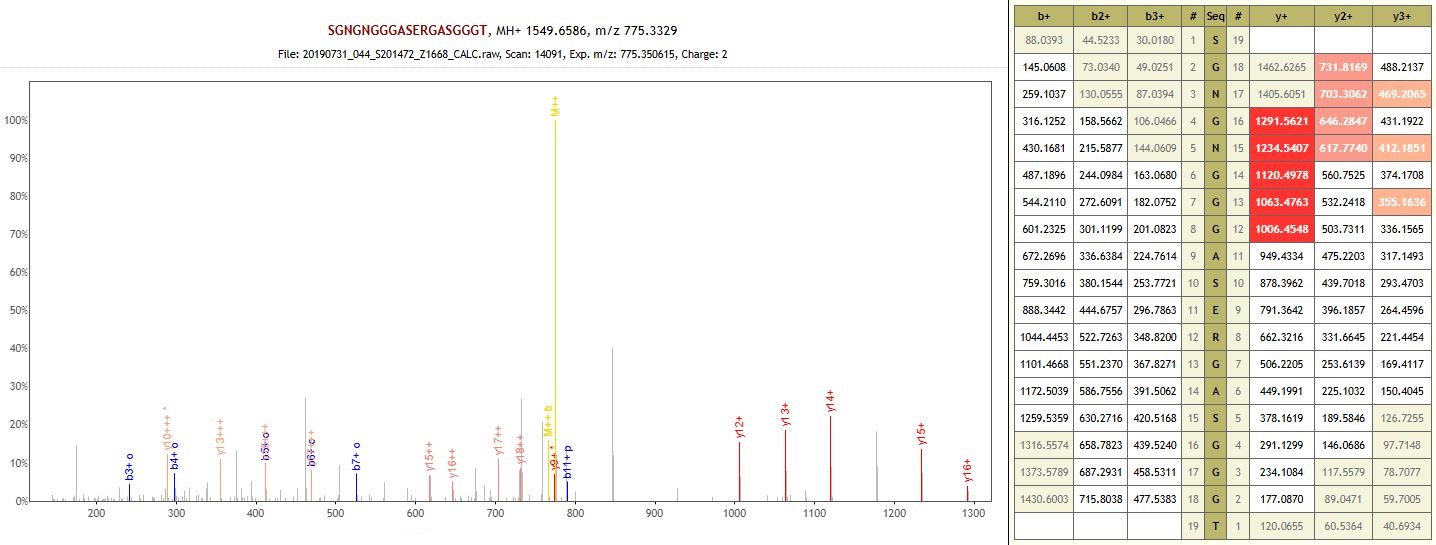

Protein: Trihelix transcription factor GTL1 (auto-enzyme)

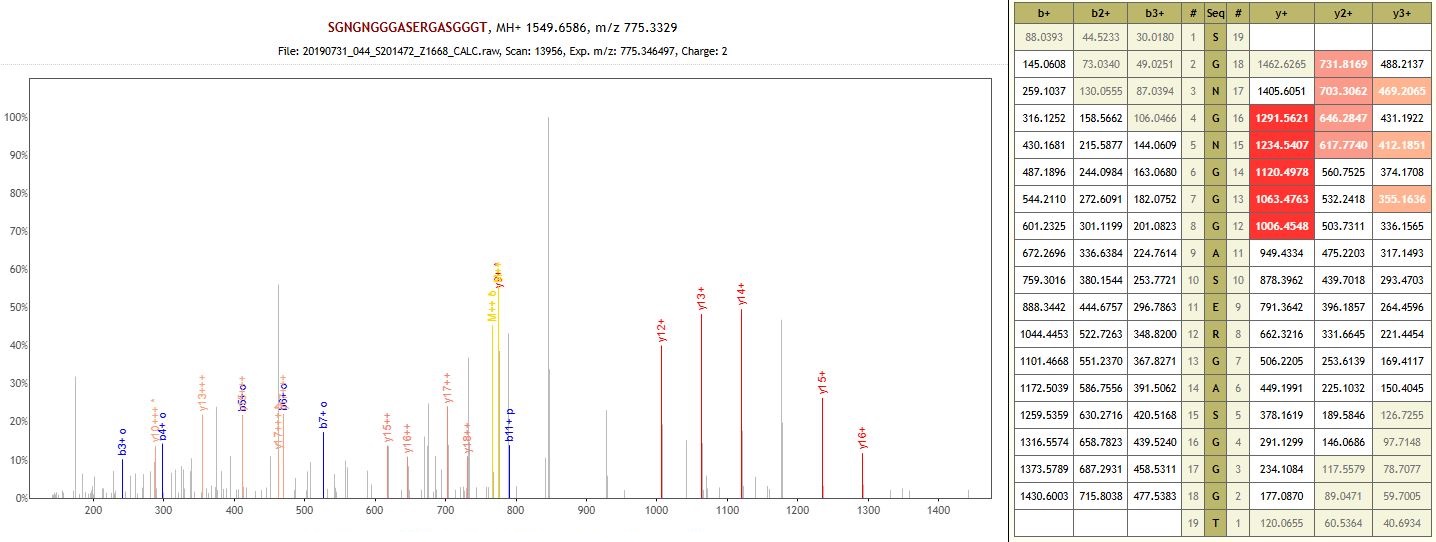
 Protein: Trihelix transcription factor GTL1 (auto-enzyme)

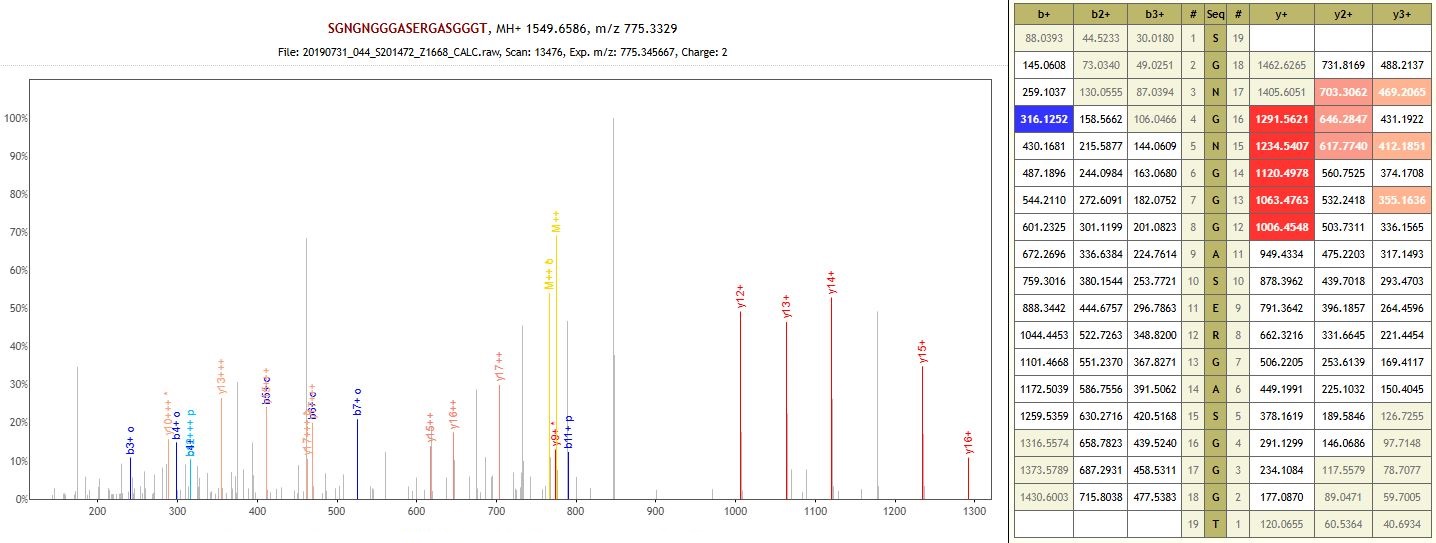
 Protein: Trihelix transcription factor GTL1 (auto-enzyme)

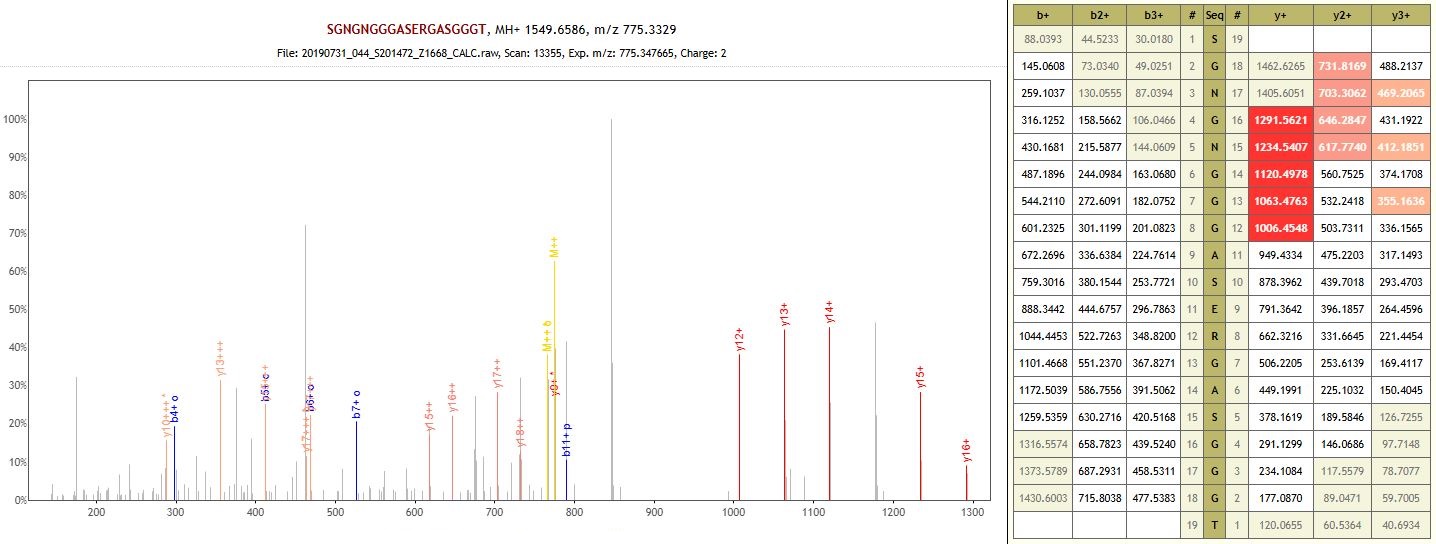
 Protein: Trihelix transcription factor GTL1 (auto-enzyme)

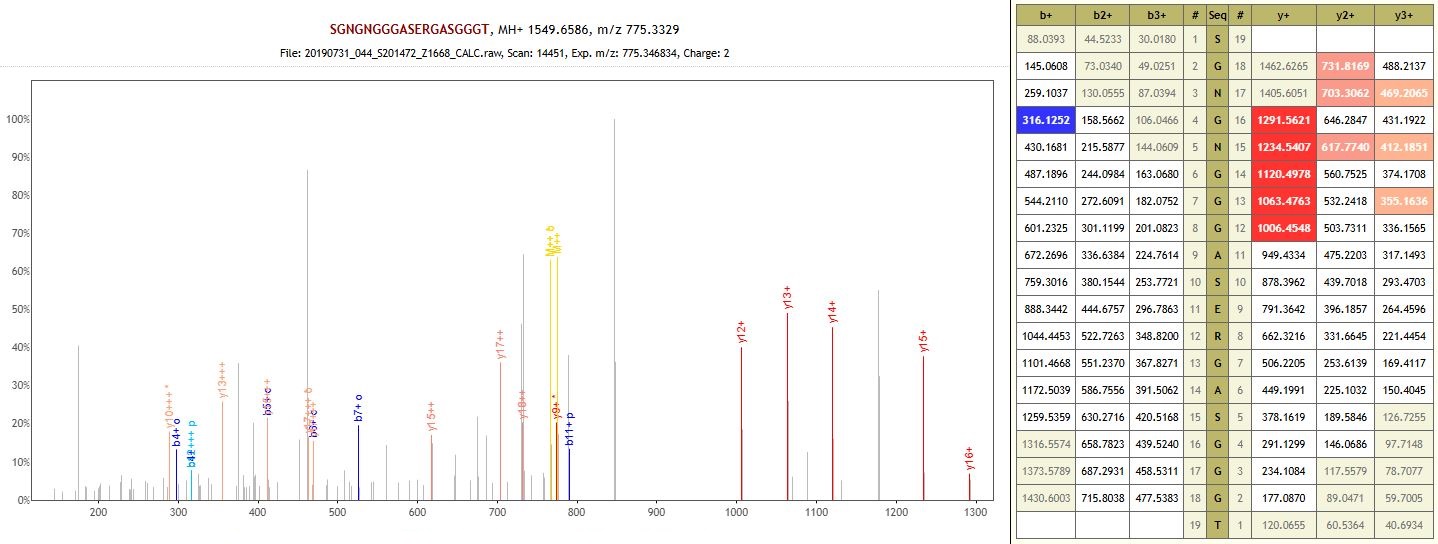
 Protein: Trihelix transcription factor GTL1 (auto-enzyme)

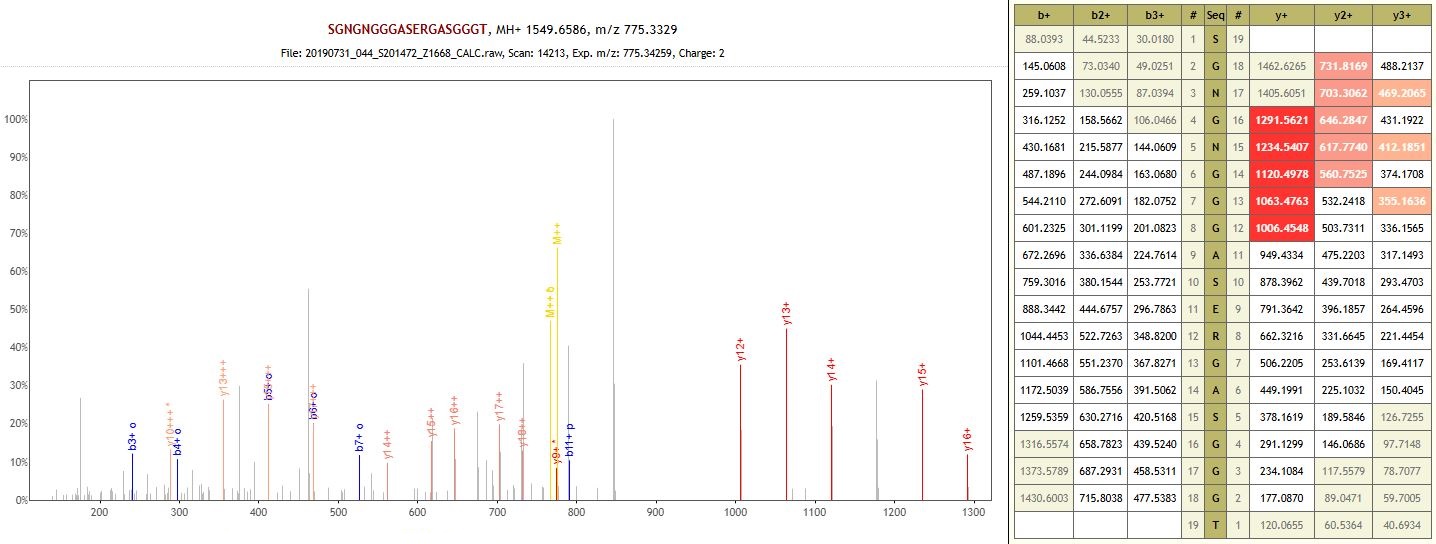
 Protein: Trihelix transcription factor GTL1 (auto-enzyme)

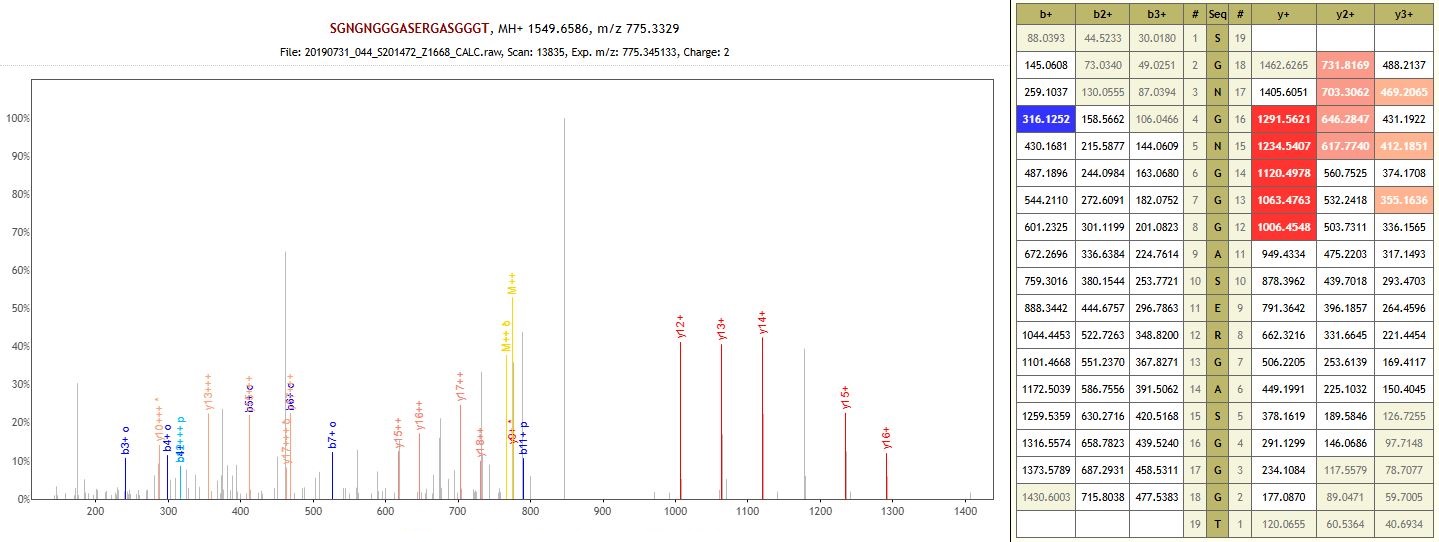
 Protein: Trihelix transcription factor GTL1 (auto-enzyme)

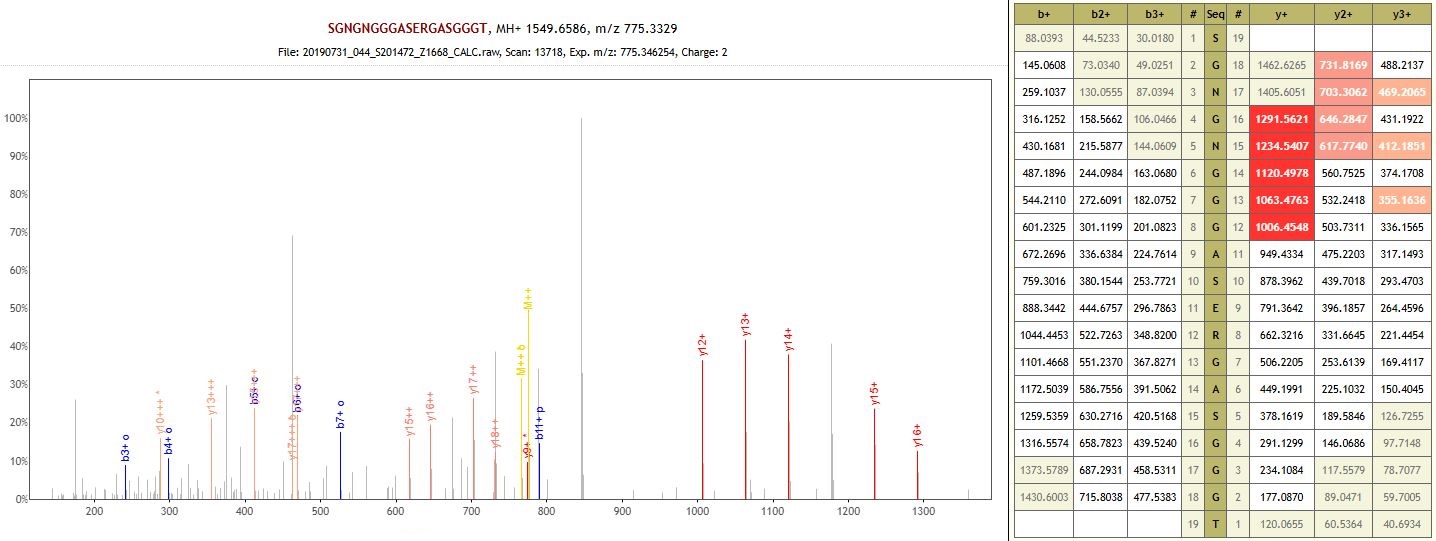
Protein: Trihelix transcription factor GTL1 (auto-enzyme)
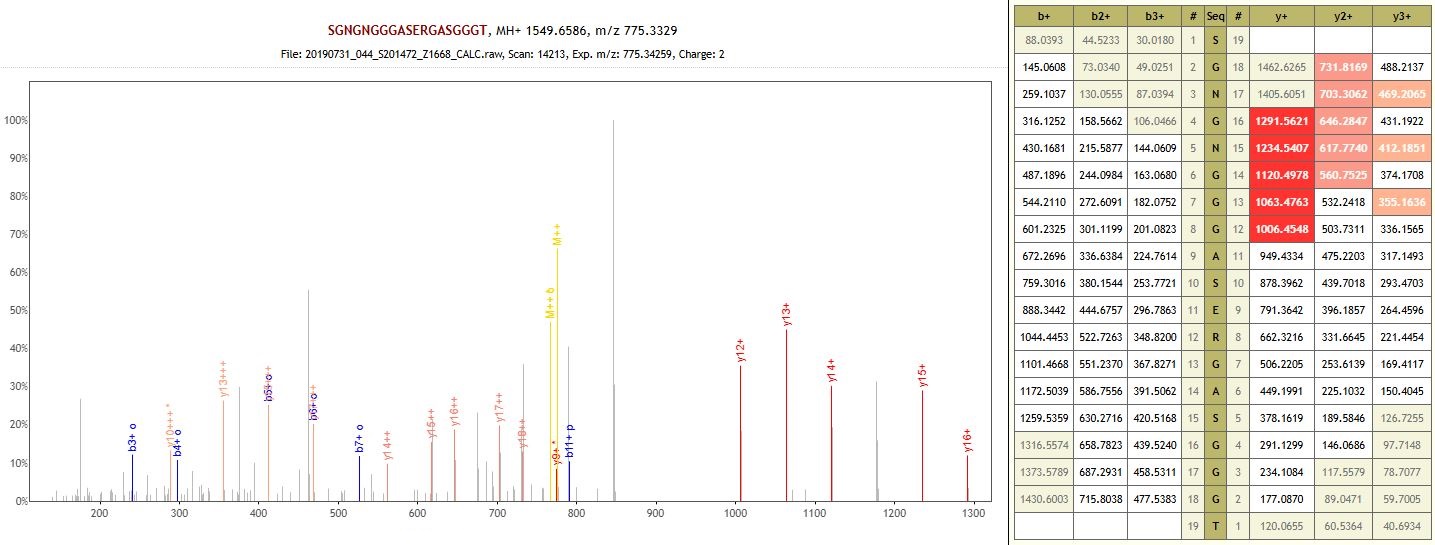

Protein: Trihelix transcription factor GTL1 (non-specific enzyme)

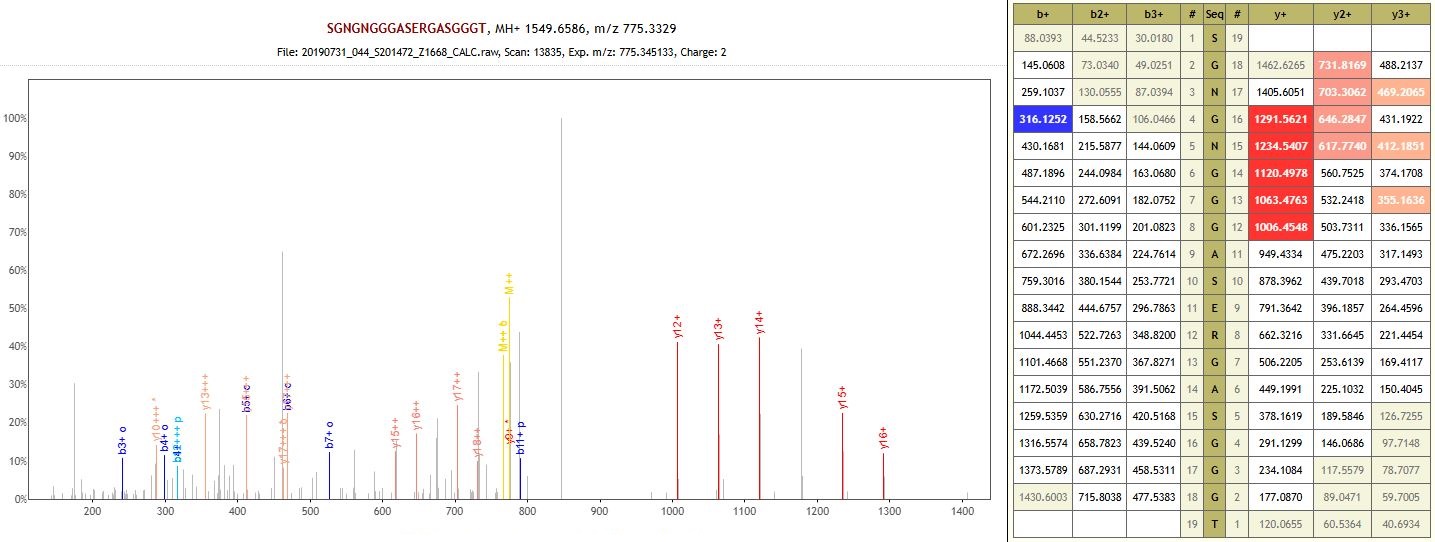

Protein: Trihelix transcription factor GTL1 (non-specific enzyme)

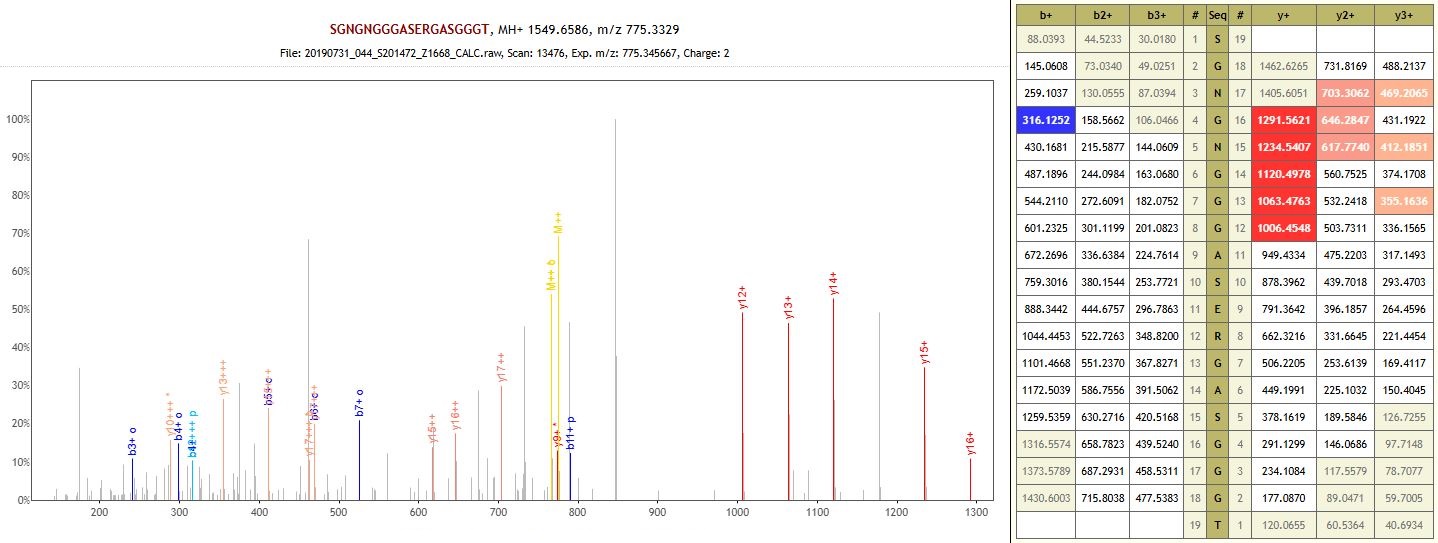

Protein: Trihelix transcription factor GTL1 (non-specific enzyme)

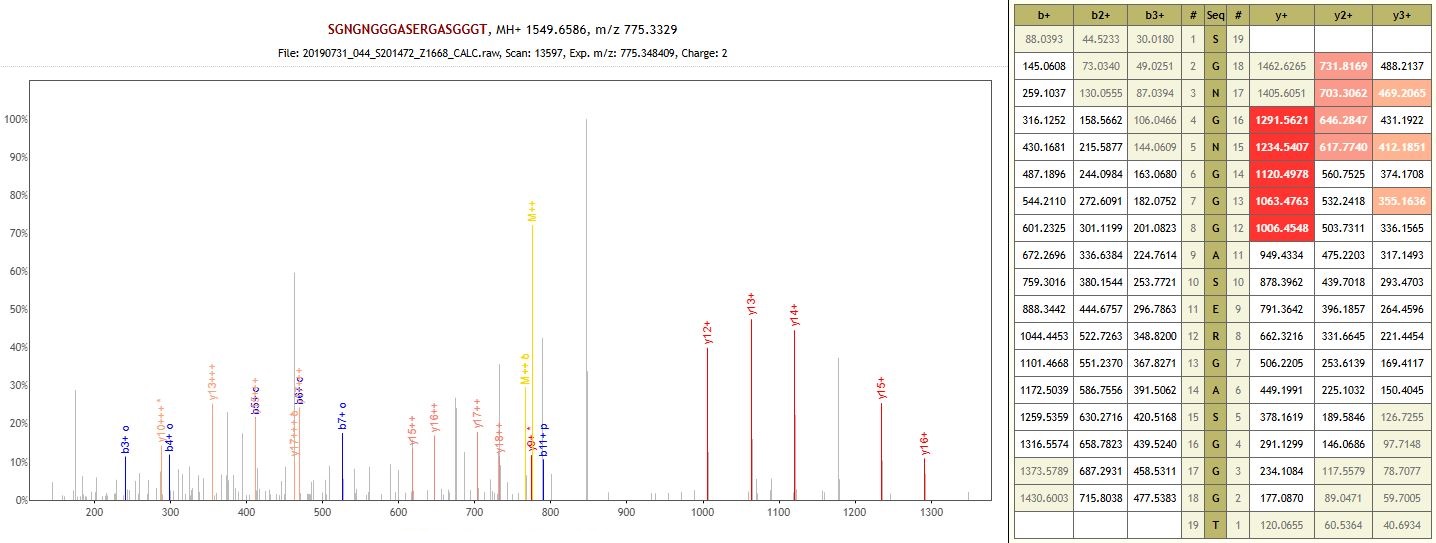

Protein: Trihelix transcription factor GTL1 (non-specific enzyme)

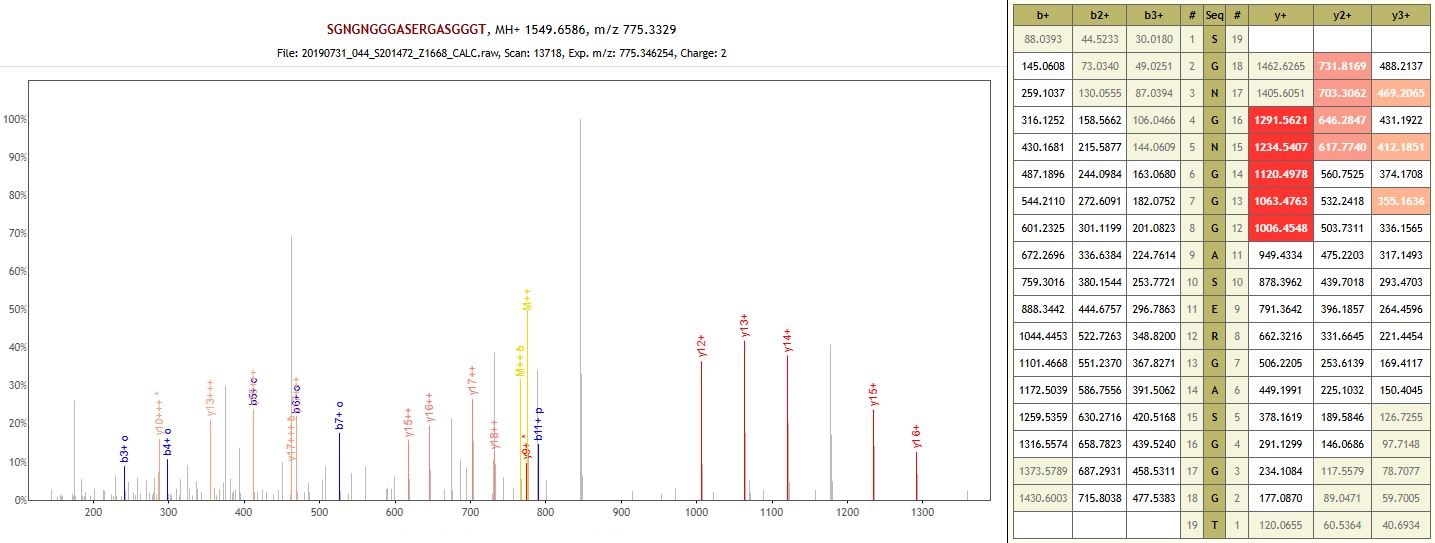

Protein: Trihelix transcription factor GTL1 (non-specific enzyme)

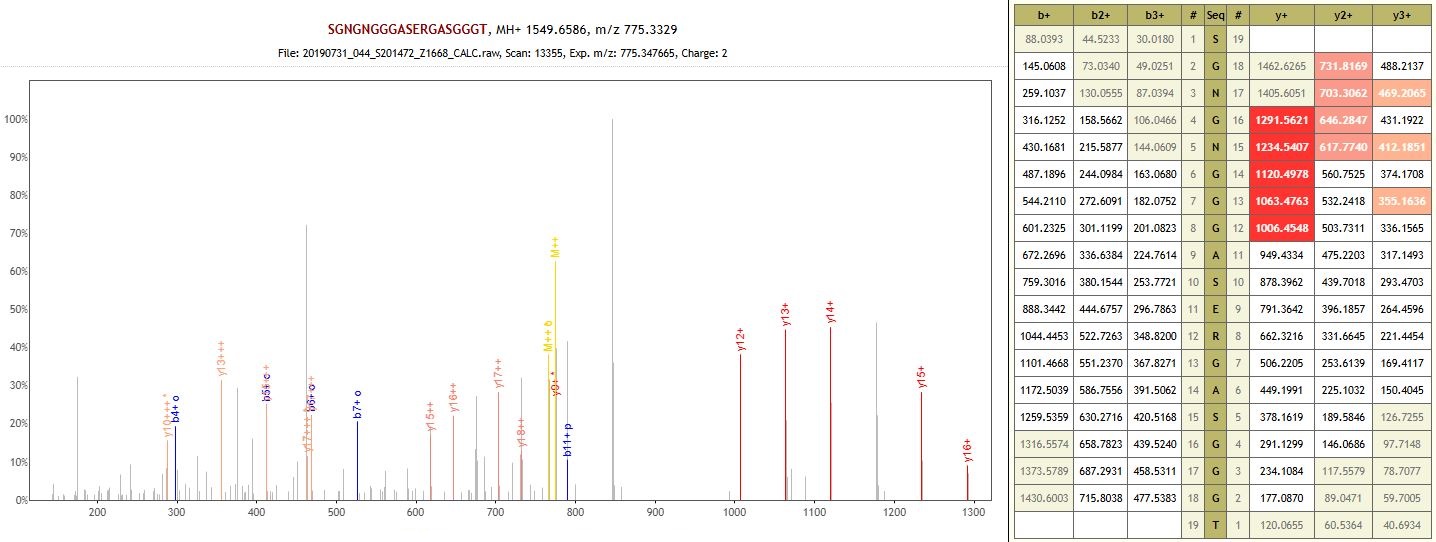

Protein: Trihelix transcription factor GTL1 (non-specific enzyme)

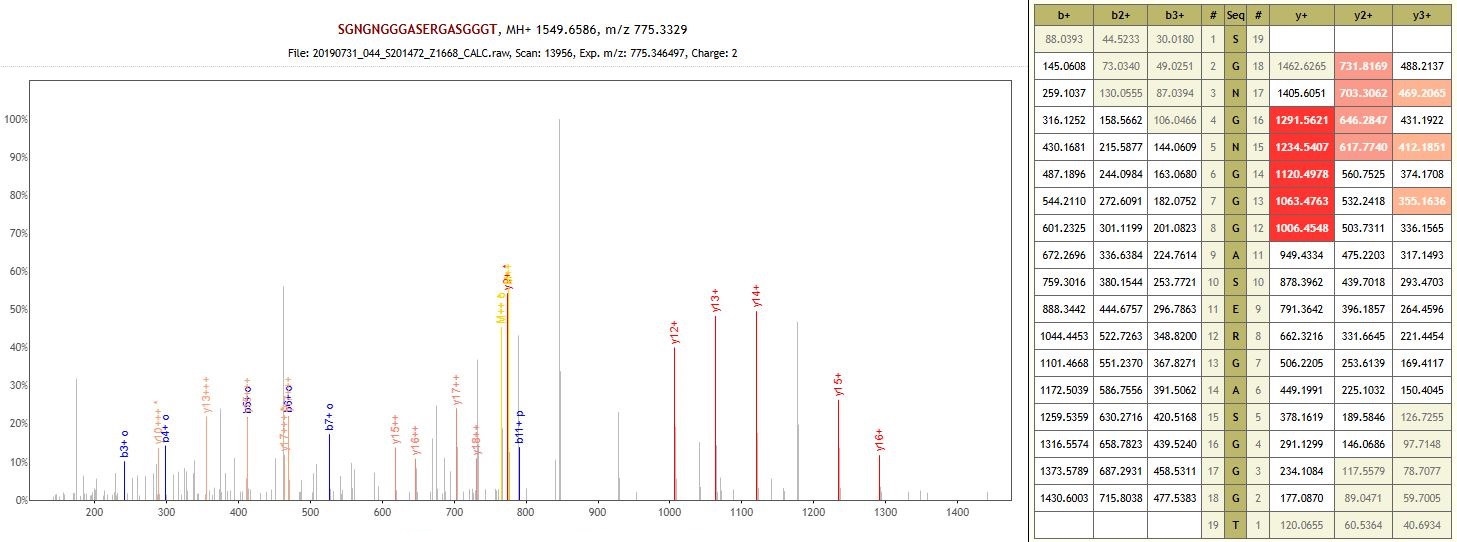

Protein: Trihelix transcription factor GTL1 (non-specific enzyme)

**MGD001 – Megiddo – Middle Bronze Age**

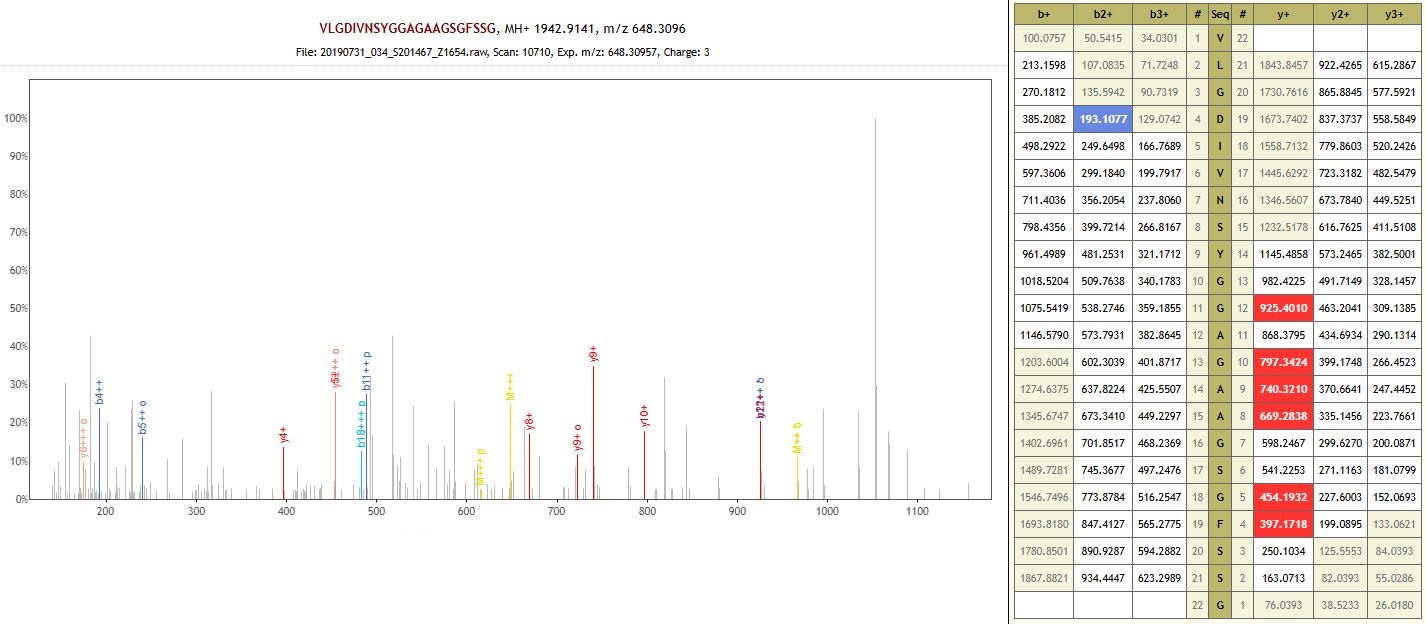

Protein: Multidrug resistance protein (non-specific enzyme)

**MGD006 – Megiddo – Late Bronze Age**
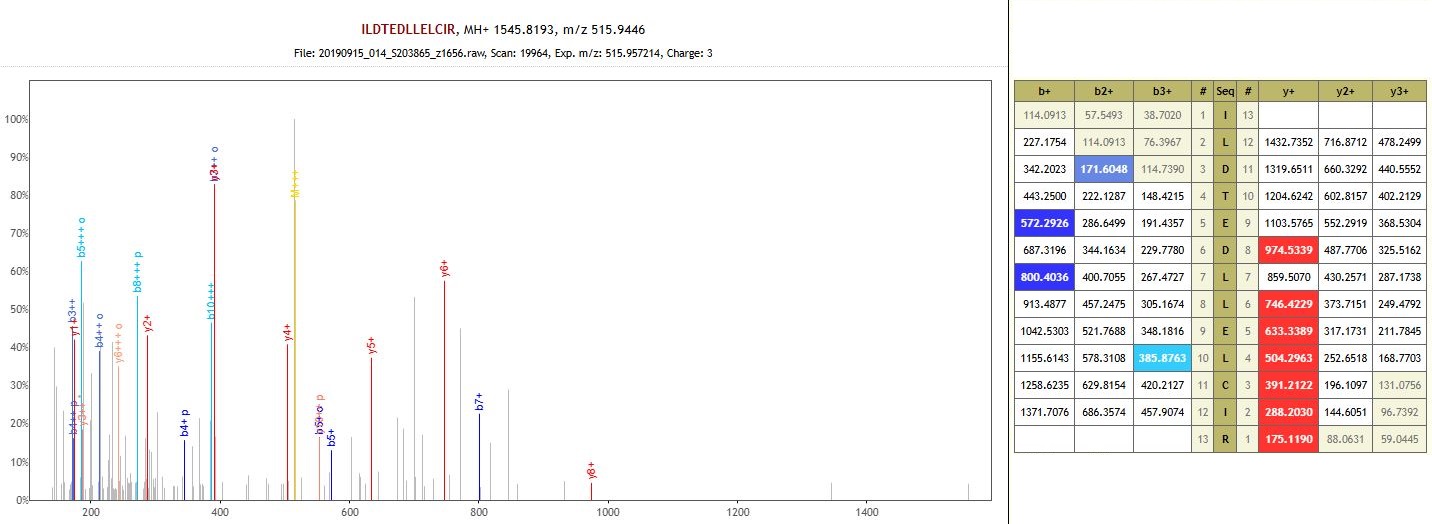

Protein: Cysteine-rich receptor-like protein kinase 10 isoform X1 (auto-enzyme)

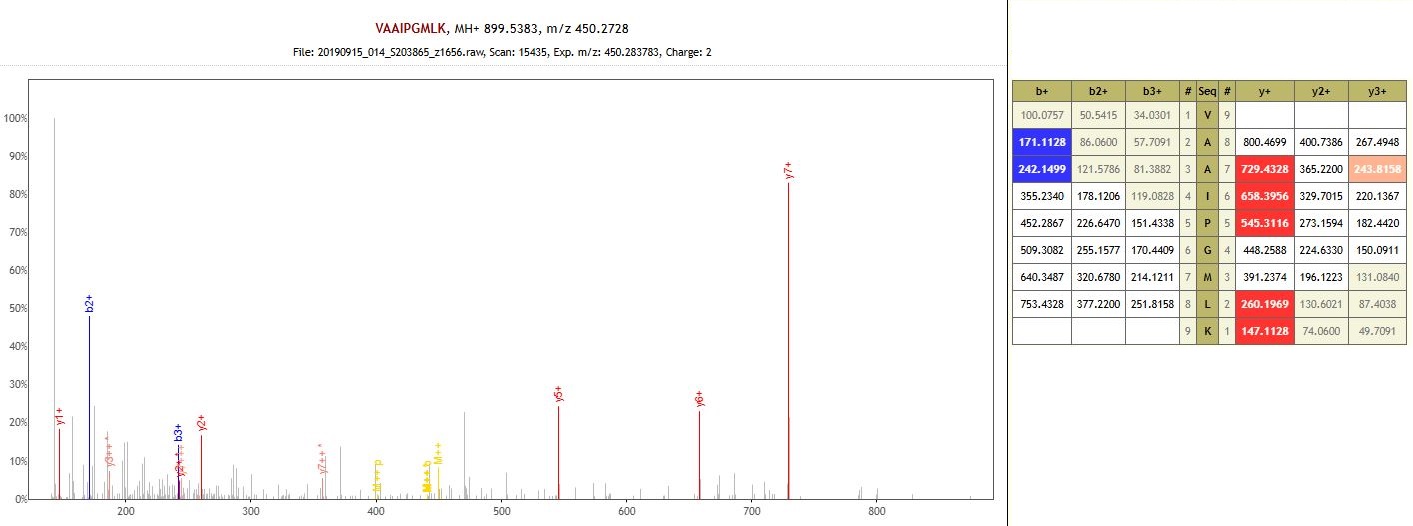
 Protein: Uncharacterized protein (auto-enzyme)

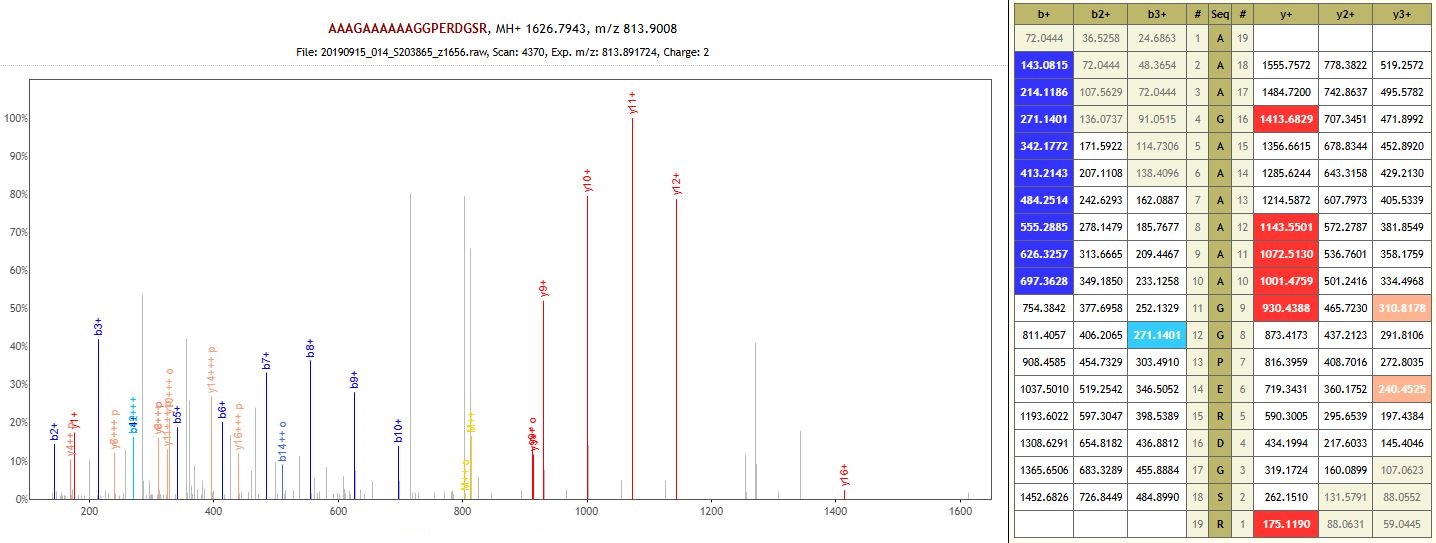

Protein: Pre-mRNA-splicing factor Syf1/CRNKL1-like C-terminal HAT-repeats domain-containing protein (non-specific enzyme)

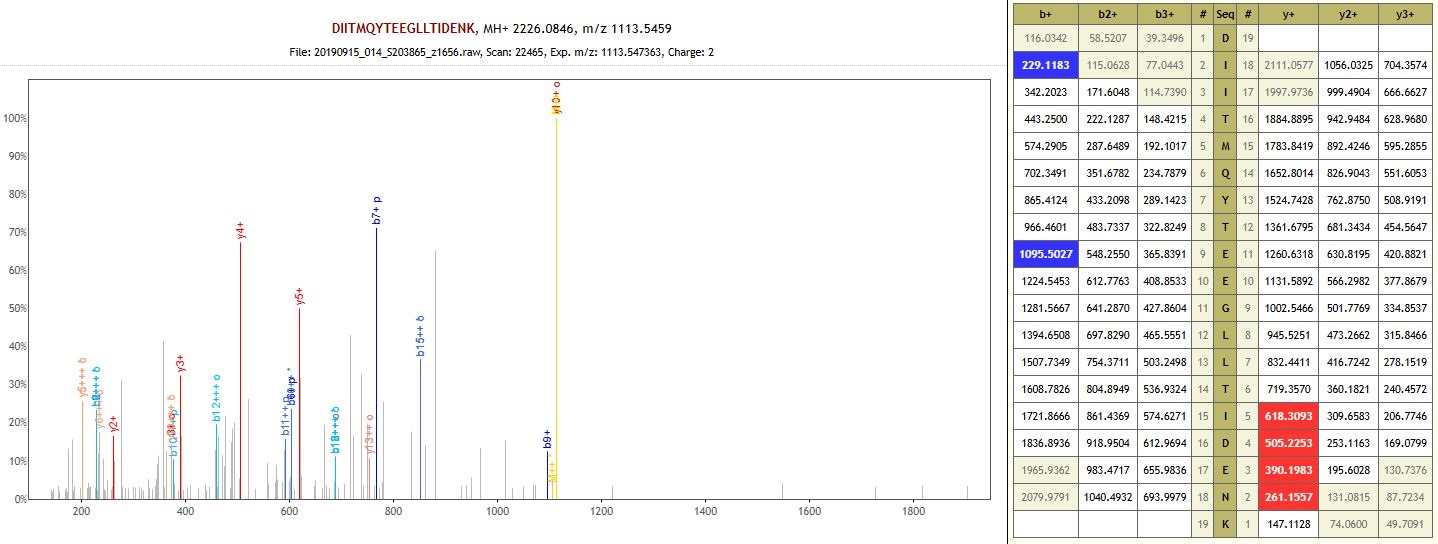

Protein: Uncharacterized protein (non-specific enzyme)

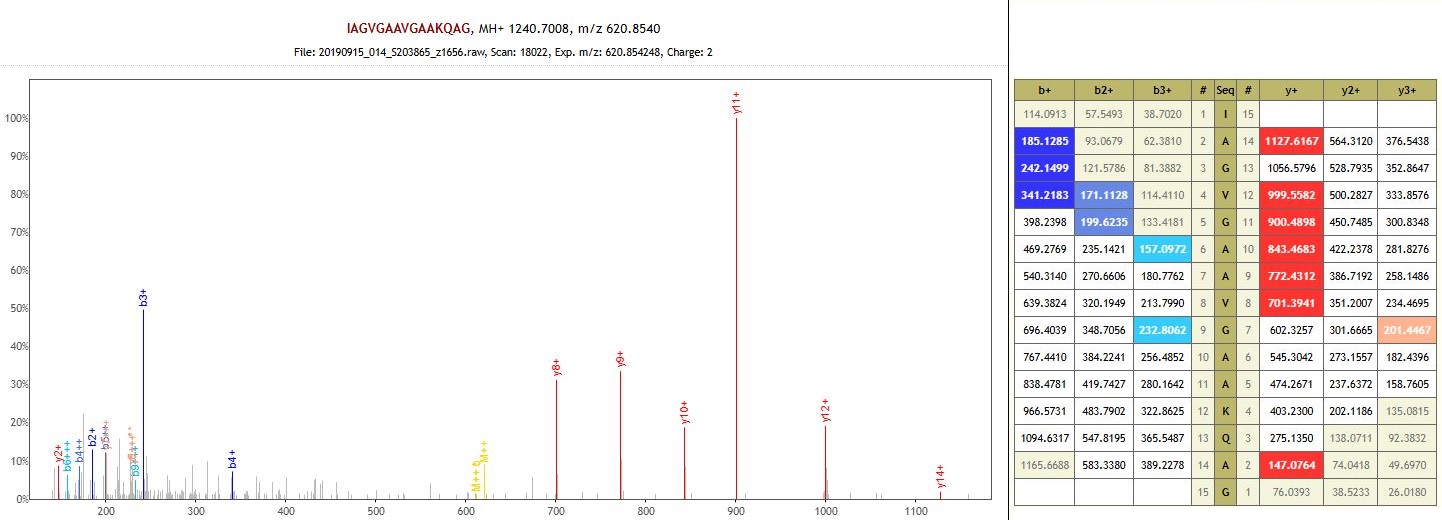

Protein: TCP domain-containing protein (non-specific enzyme)

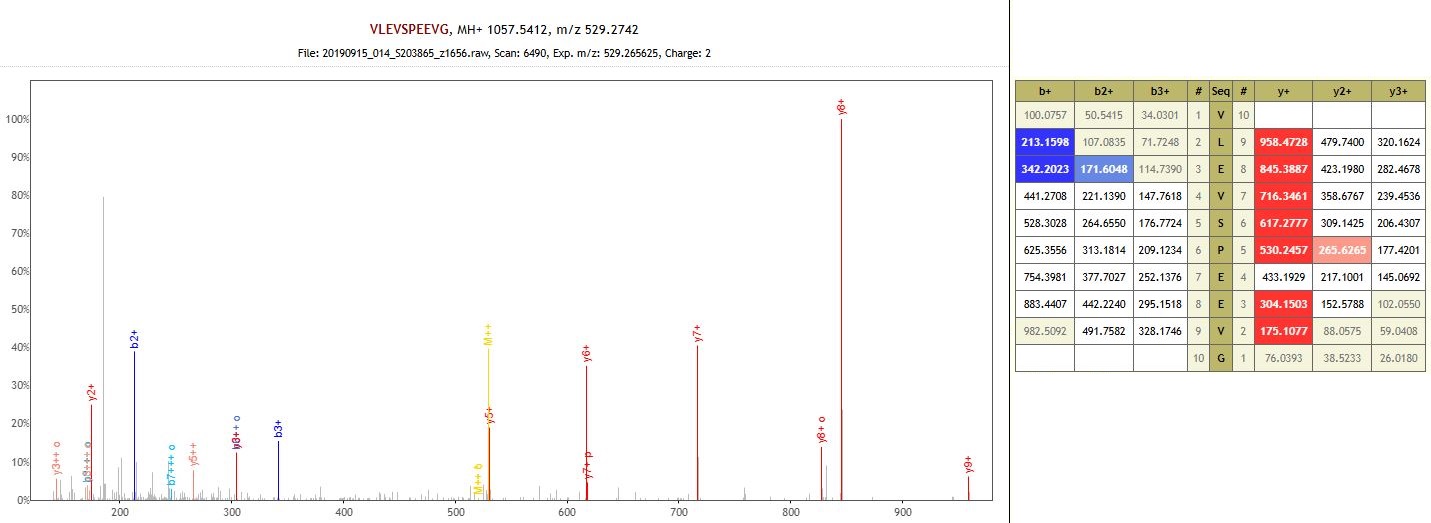

Protein: Uncharacterized protein (non-specific enzyme)

**MGD007 – Megiddo – Late Bronze Age**

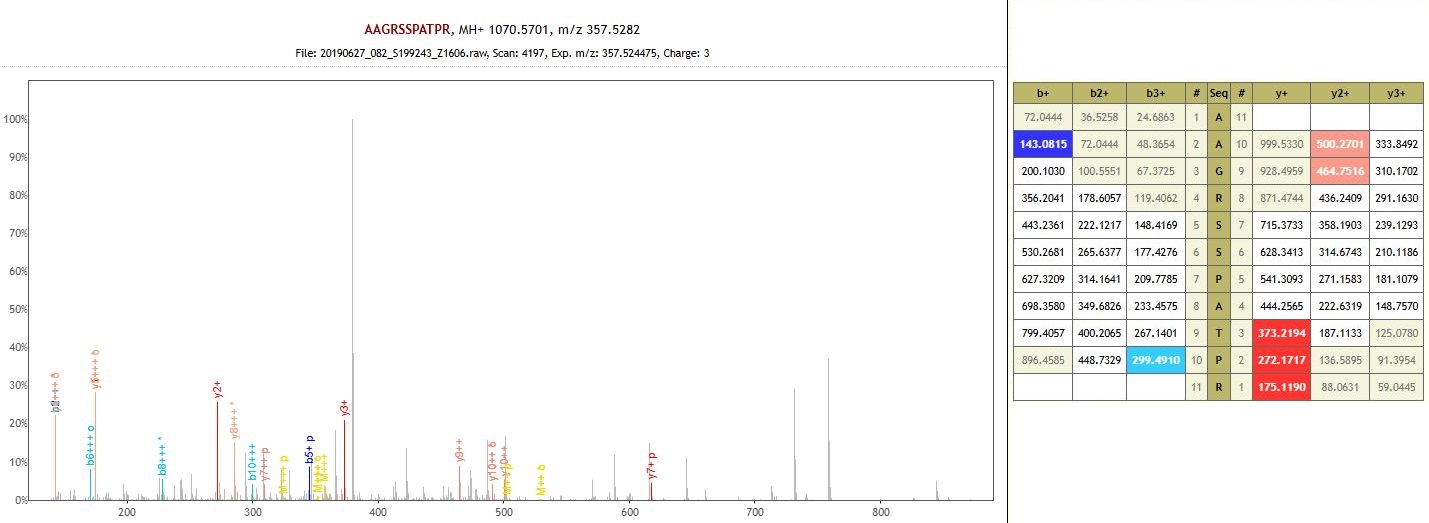

Protein: Agenet domain-containing protein (non-specific enzyme)

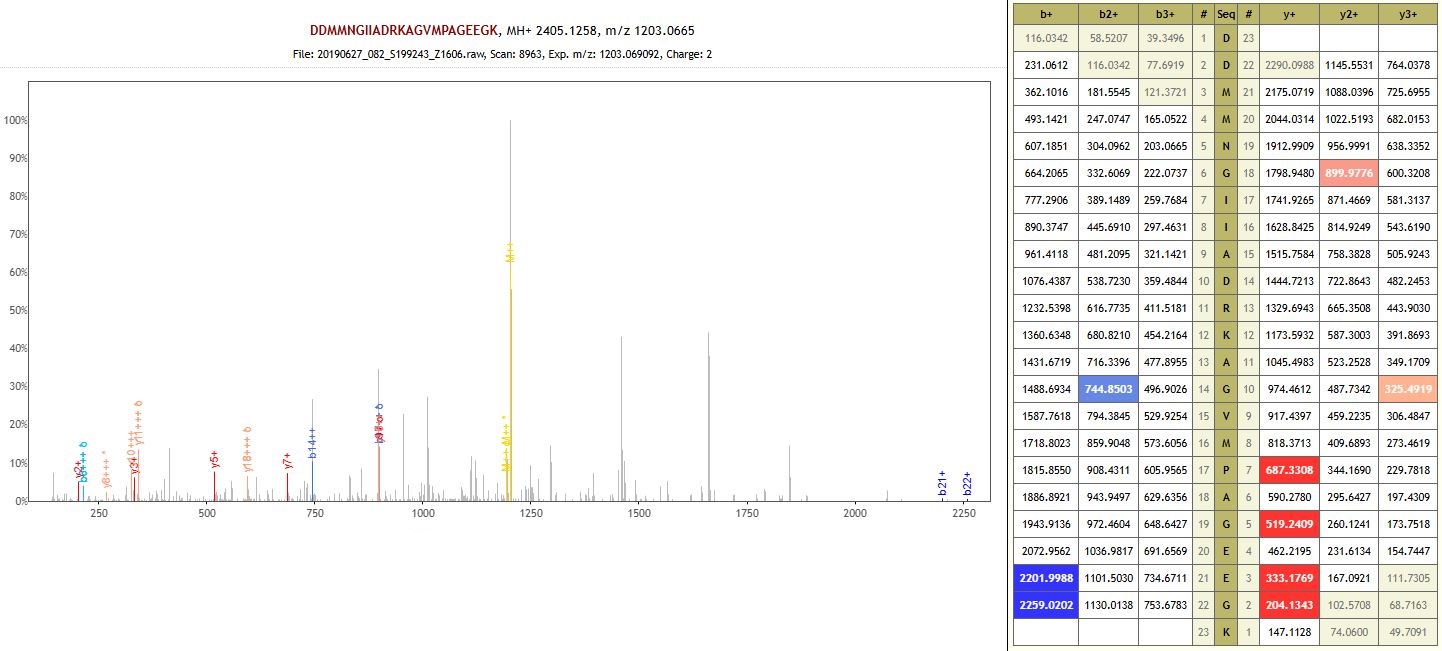
Protein: Flavonoid 3 (non-specific enzyme)

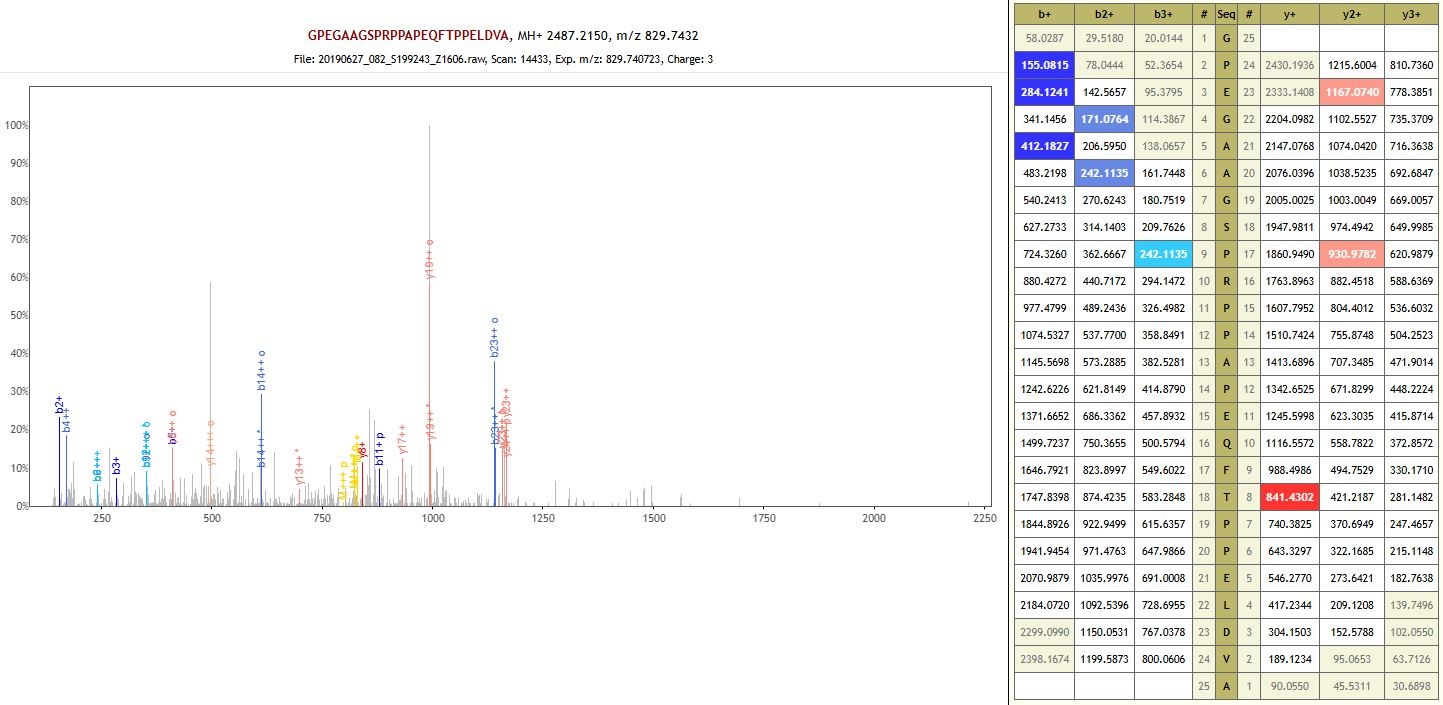

Protein: Uncharacterized protein (non-specific enzyme)

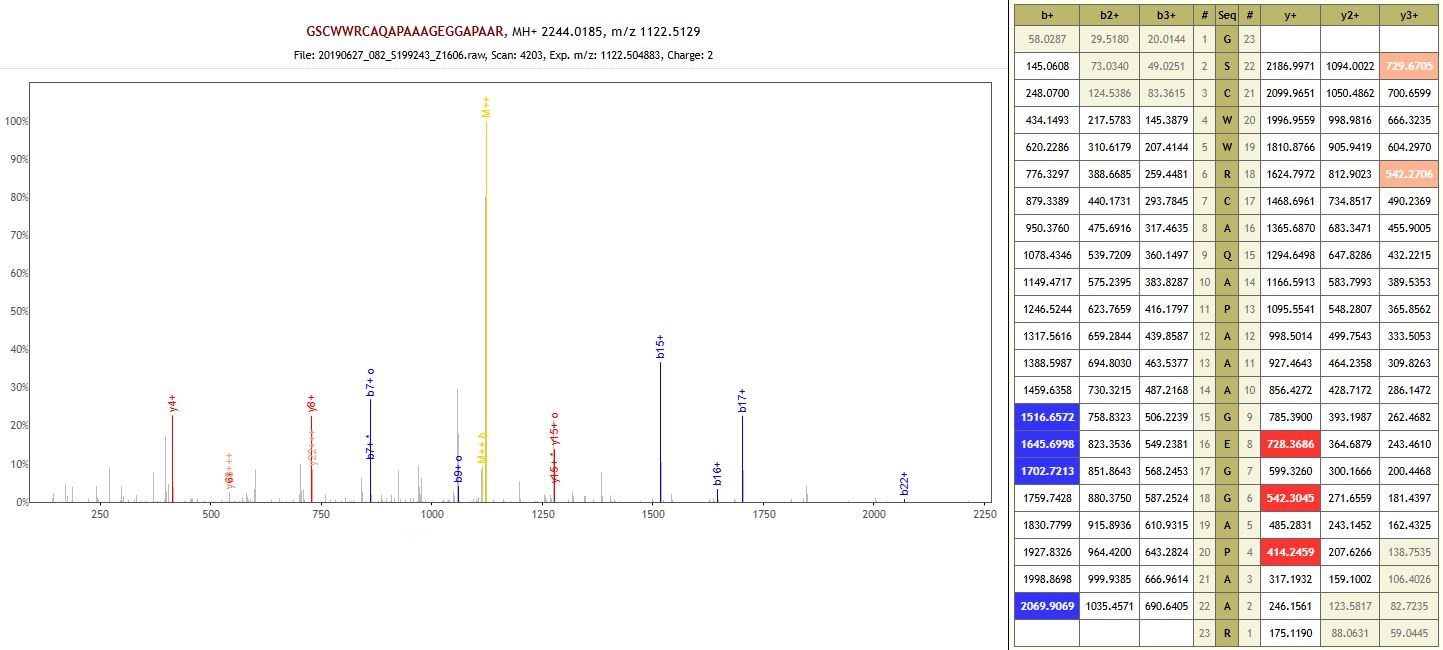

Protein: Uncharacterized protein (non-specific enzyme)

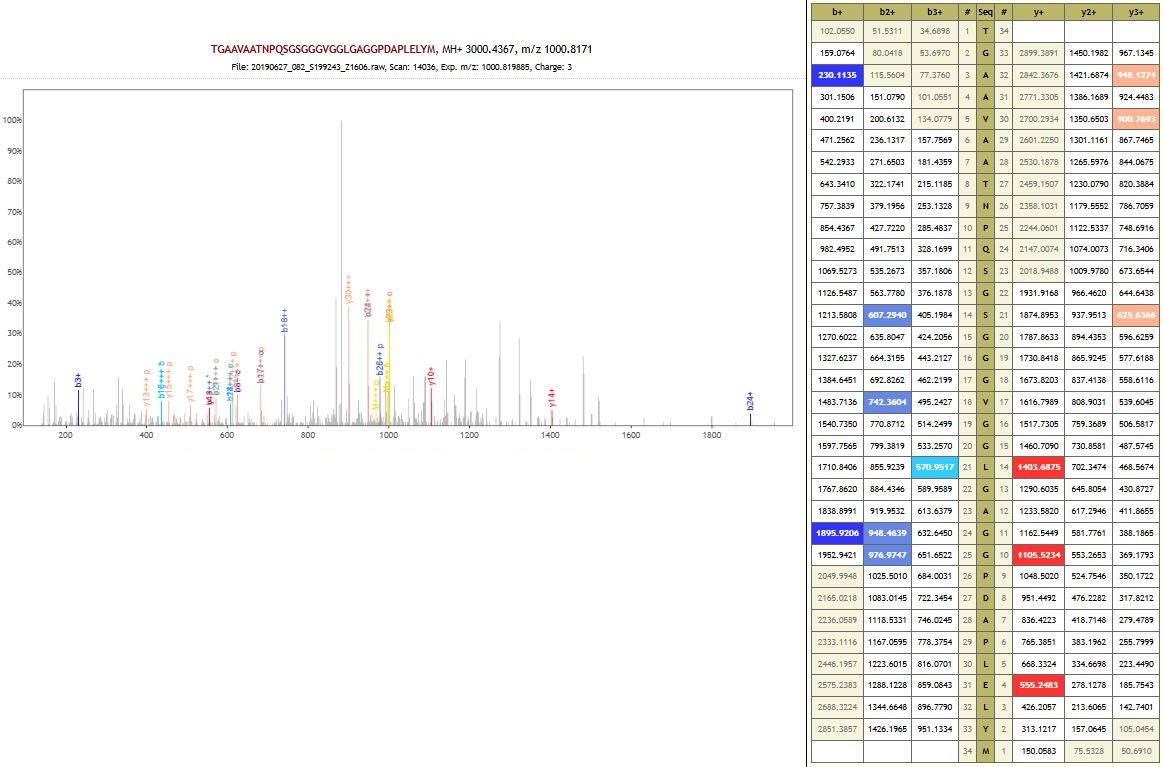

Protein: Dirigent protein (non-specific enzyme)

**MGD009 – Megiddo – Middle Bronze Age**

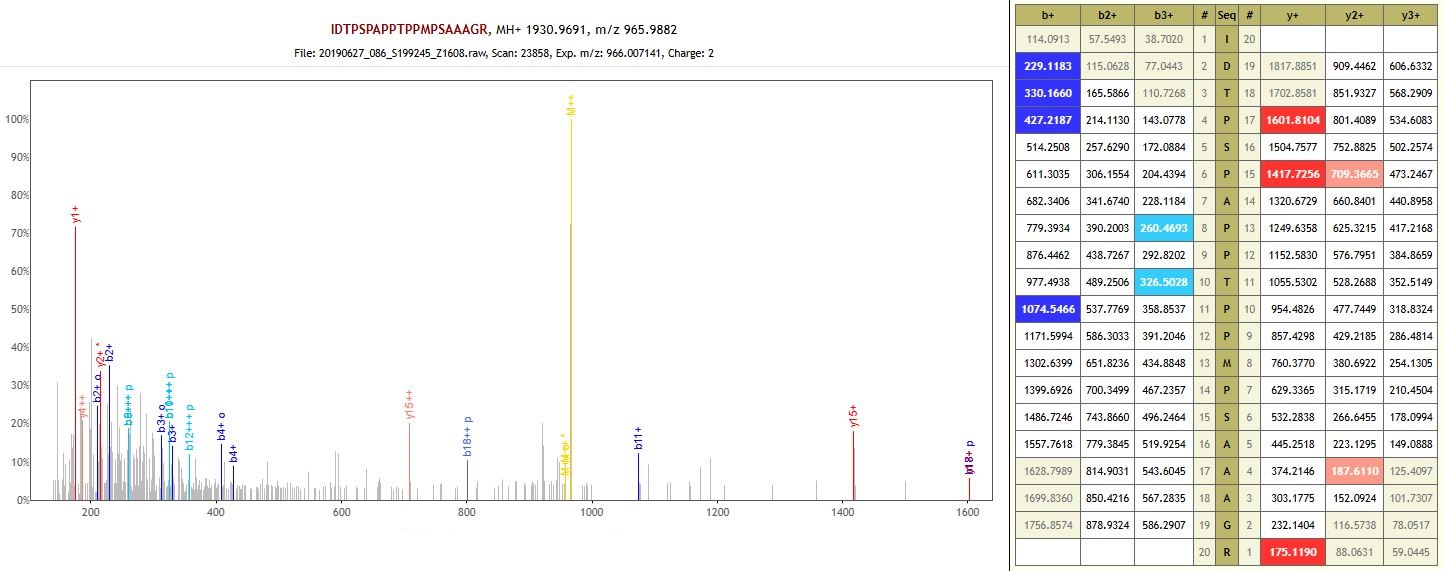
 Protein: Macrophage erythroblast attacher (auto-enzyme)

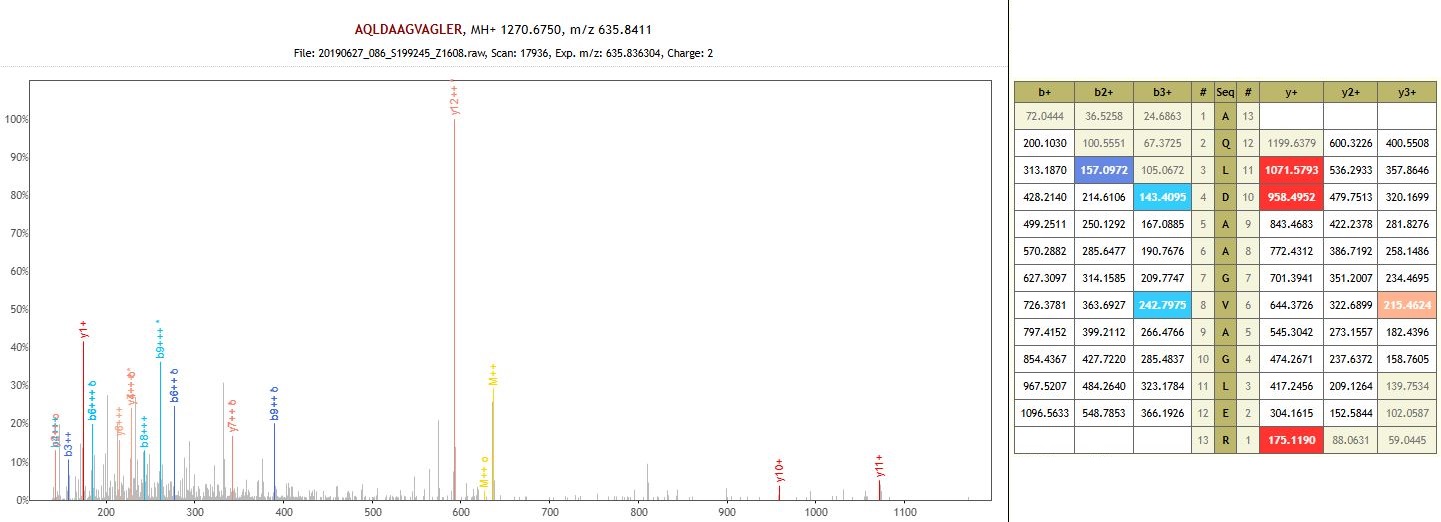
 Protein: DUF7796 domain-containing protein (auto-enzyme)

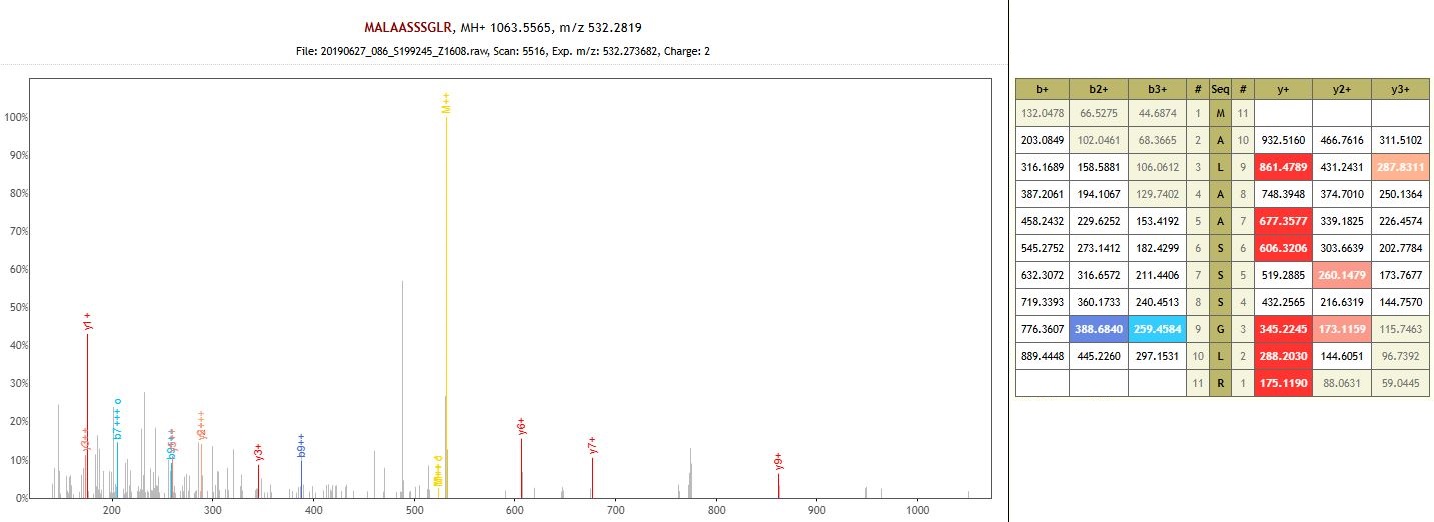
Protein: FAD-binding domain-containing protein (auto-enzyme)
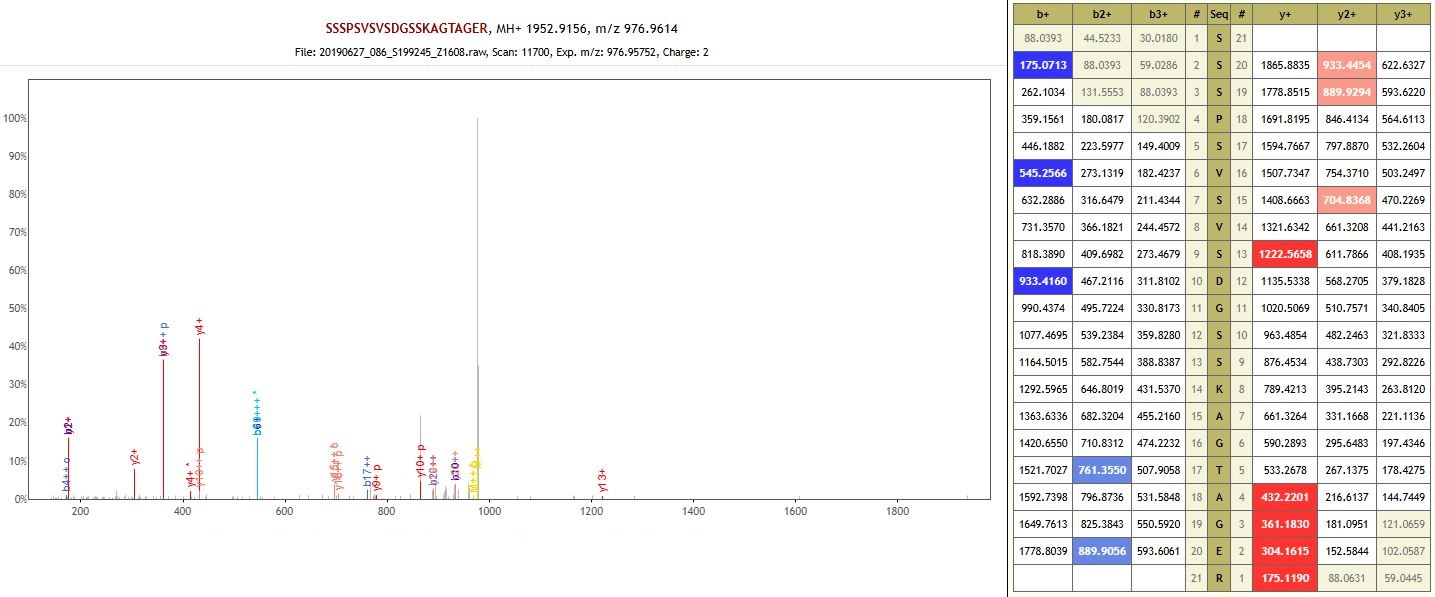

Protein: PRONE domain-containing protein (auto-enzyme)

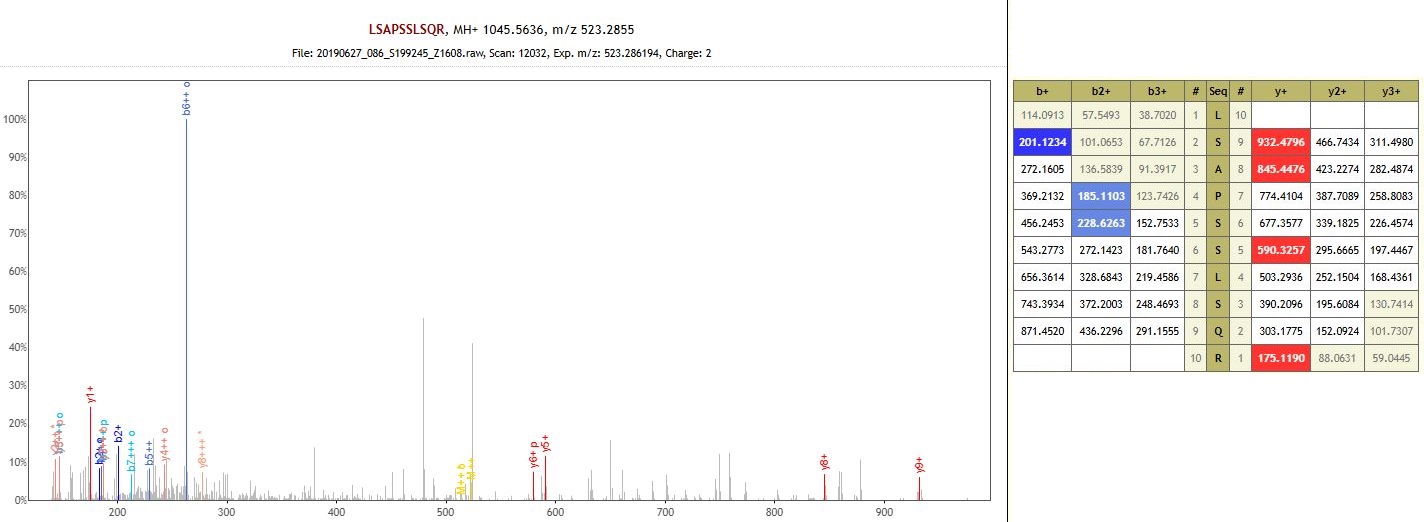

Protein: Uncharacterized protein (non-specific enzyme)

**MGD017 – Megiddo – Middle Bronze Age**

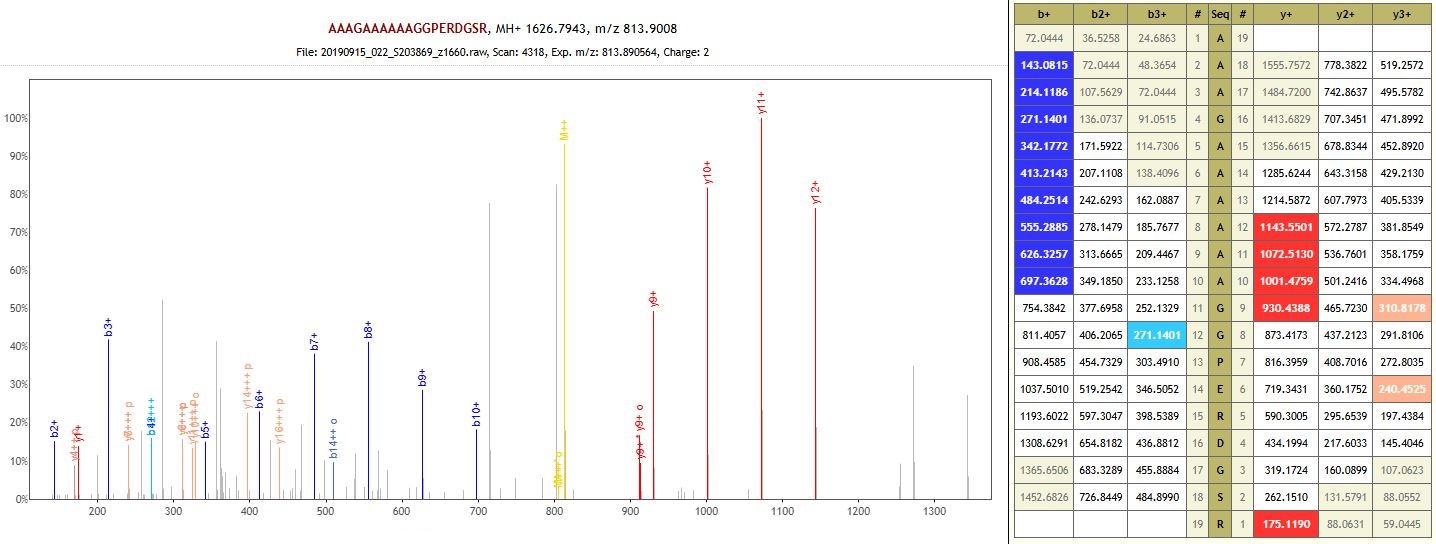

Protein: Pre-mRNA-splicing factor Syf1/CRNKL1-like C-terminal HAT-repeats domain-containing protein (non-specific enzyme)

**MGD018 – Megiddo – Middle Bronze Age**

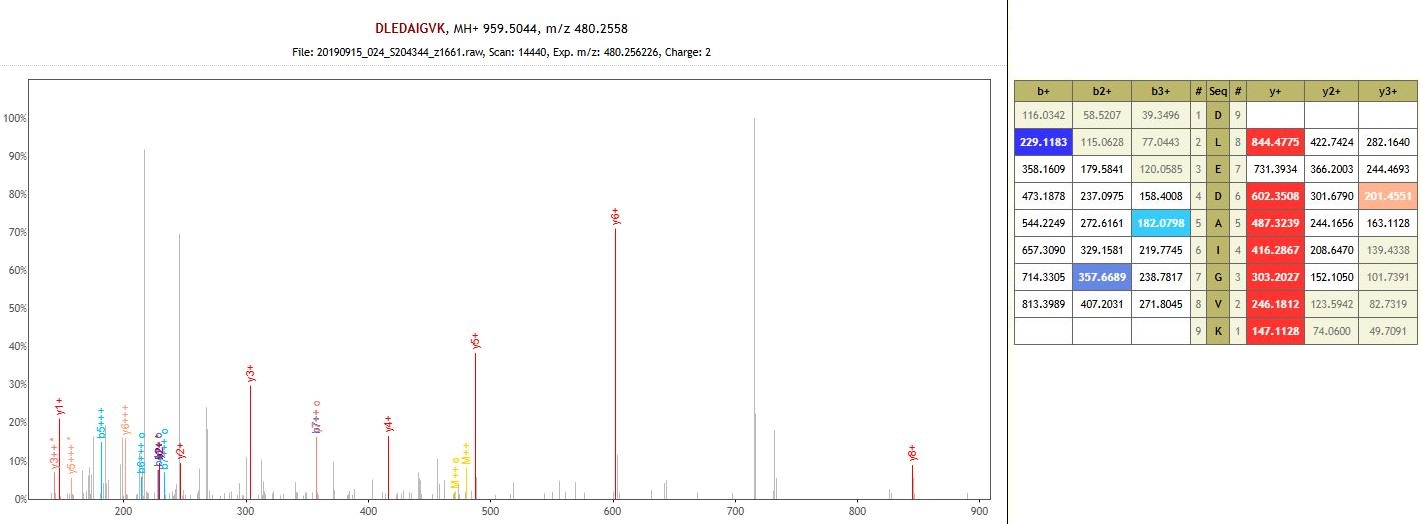

Protein: Disease resistance protein RPM1-like (non-specific enzyme)

**DA094 – Krasikovskyi 1 – Early Bronze Age**

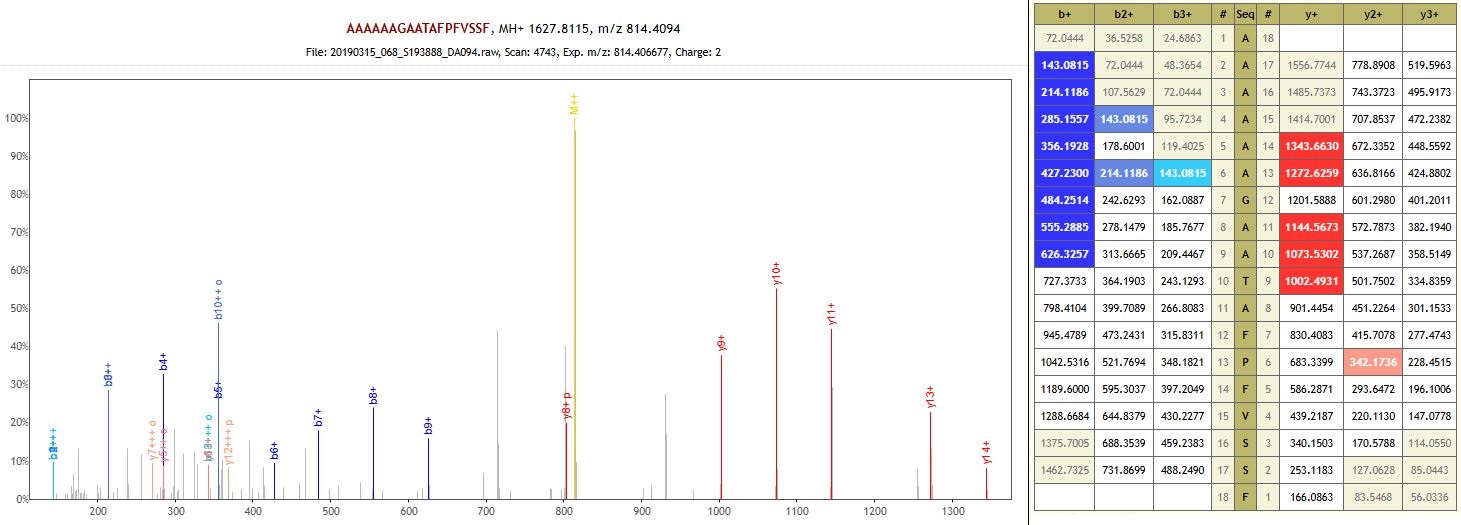

Protein: Peptidoglycan binding-like domain-containing protein (non-specific enzyme)

**DA095 – Krasikovskyi 1 – Middle Bronze Age**

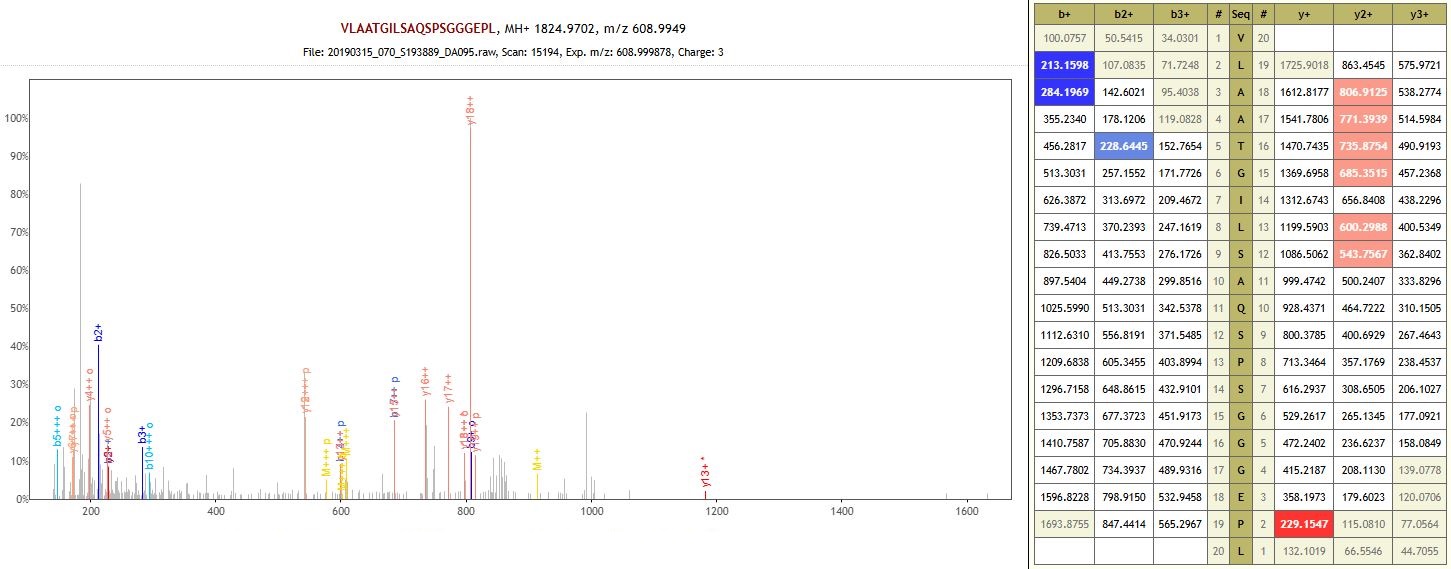

Protein: O-methyltransferase ZRP4-like (auto-enzyme)

**DA096 – Lopatino 1 – Early Bronze Age**
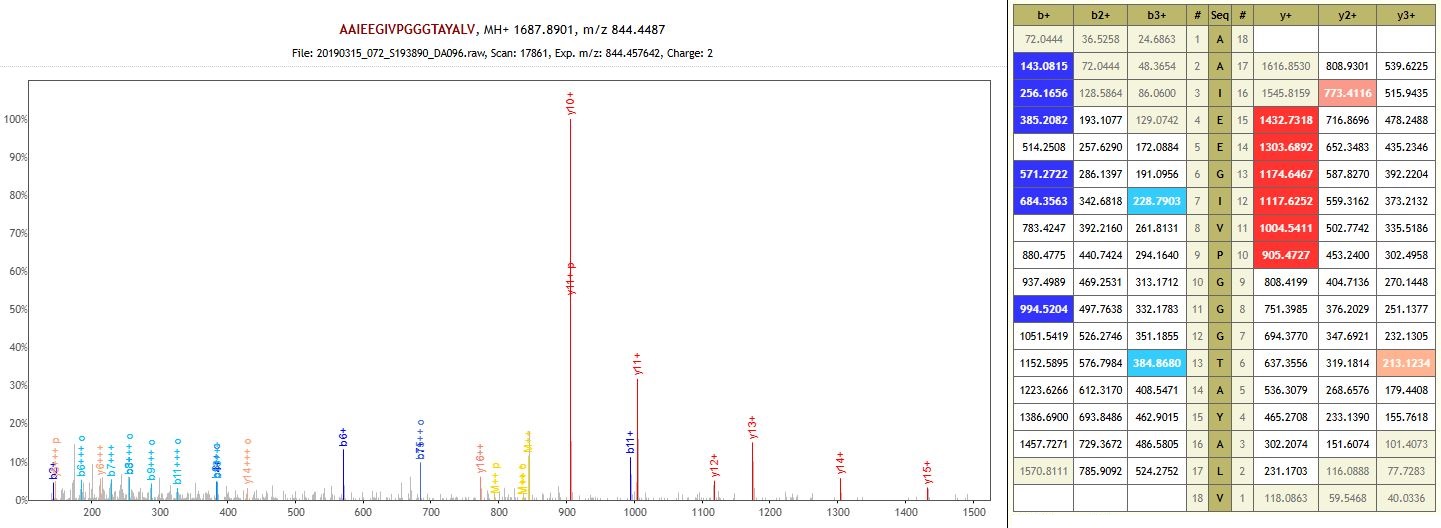

Protein: RuBisCO large subunit-binding protein subunit alpha, chloroplastic (auto-enzyme)

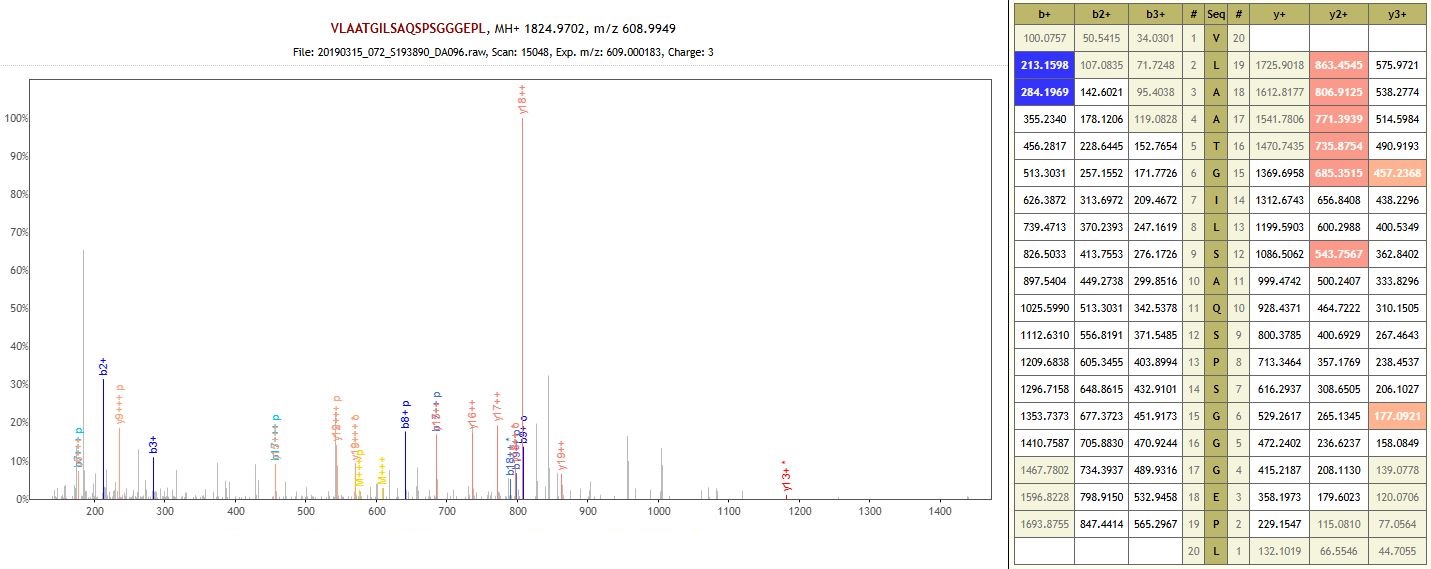

Protein: O-methyltransferase ZRP4-like (auto-enzyme)

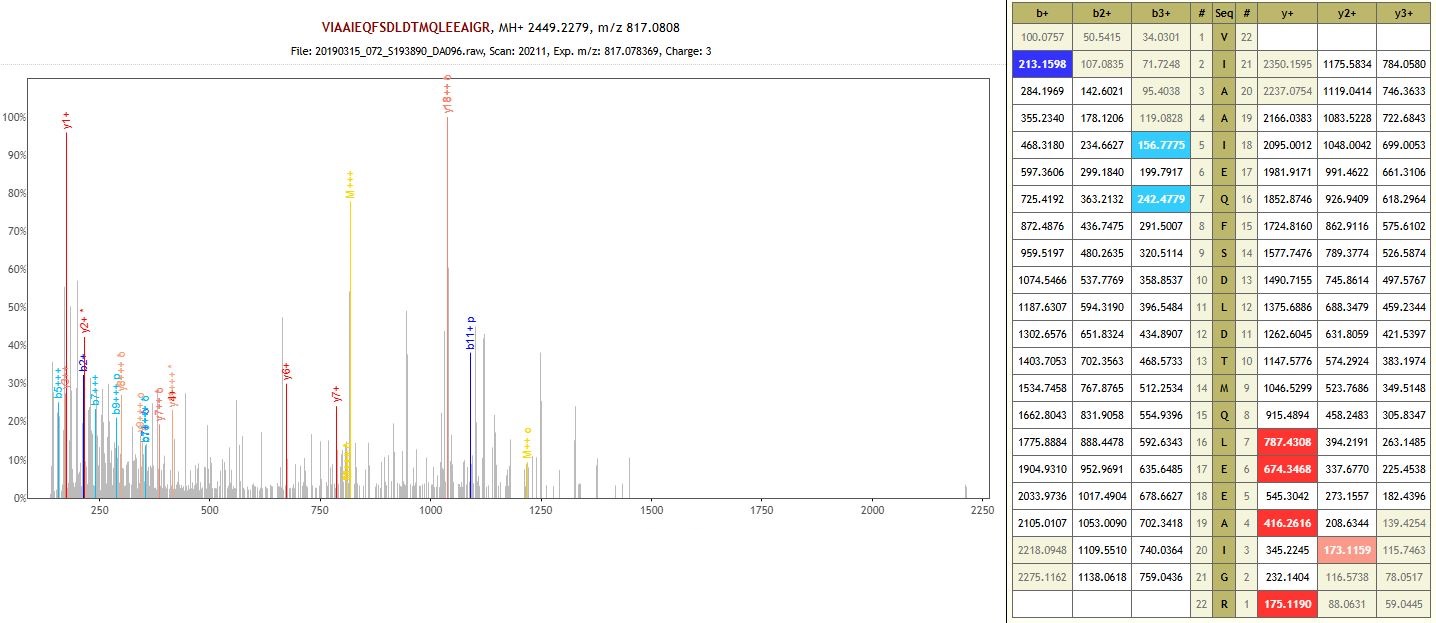

Protein: Retrotransposon protein, putative, Ty1-copia subclass (non-specific enzyme)

**DA097 – Khavalynsk 2 – Eneolithic**

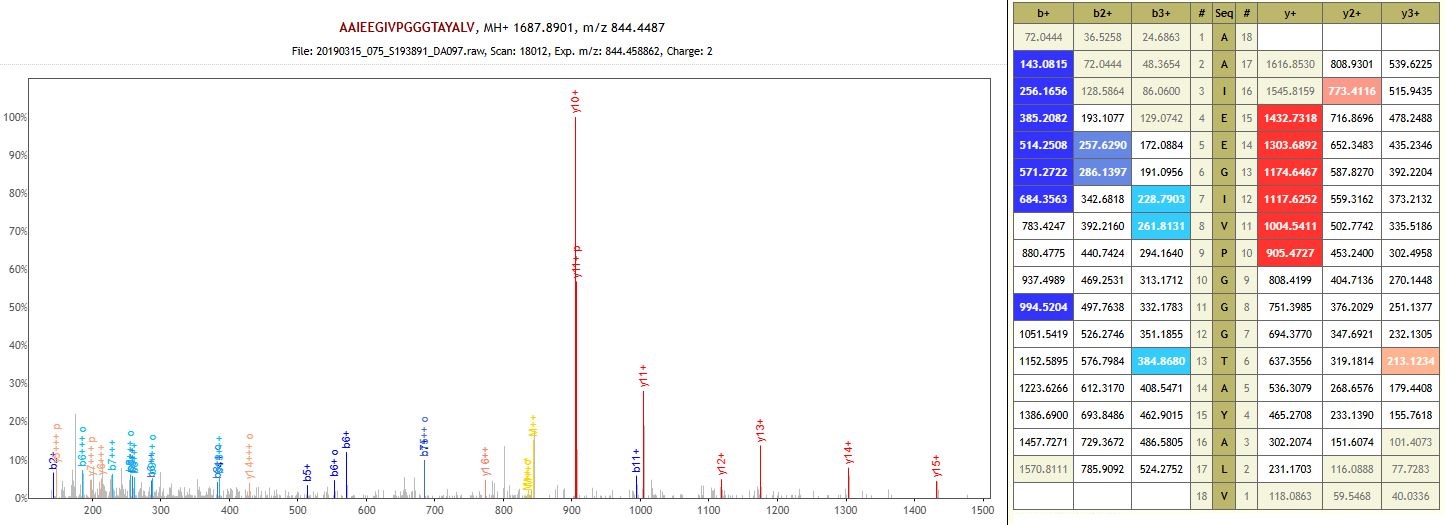

Protein: RuBisCO large subunit-binding protein subunit alpha, chloroplastic (auto-enzyme)

**DA101 – Utevka 6 – Middle Bronze Age**

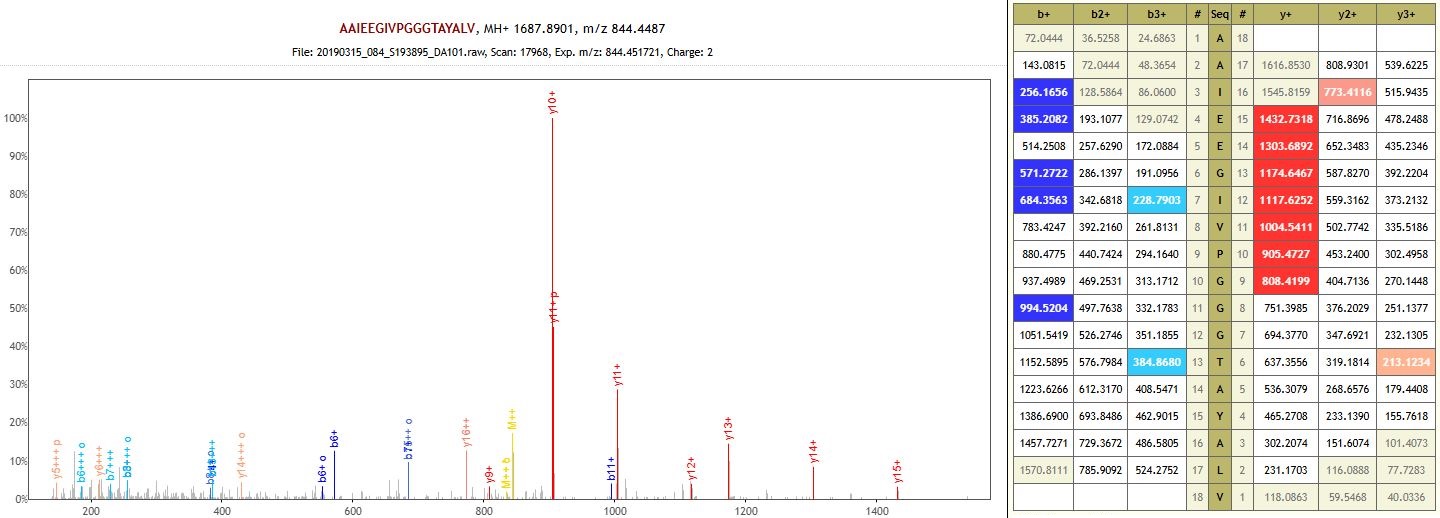

Protein: RuBisCO large subunit-binding protein subunit alpha, chloroplastic (auto-enzyme)

**DA104 – Krivyanskiy 9 – Late Bronze Age**

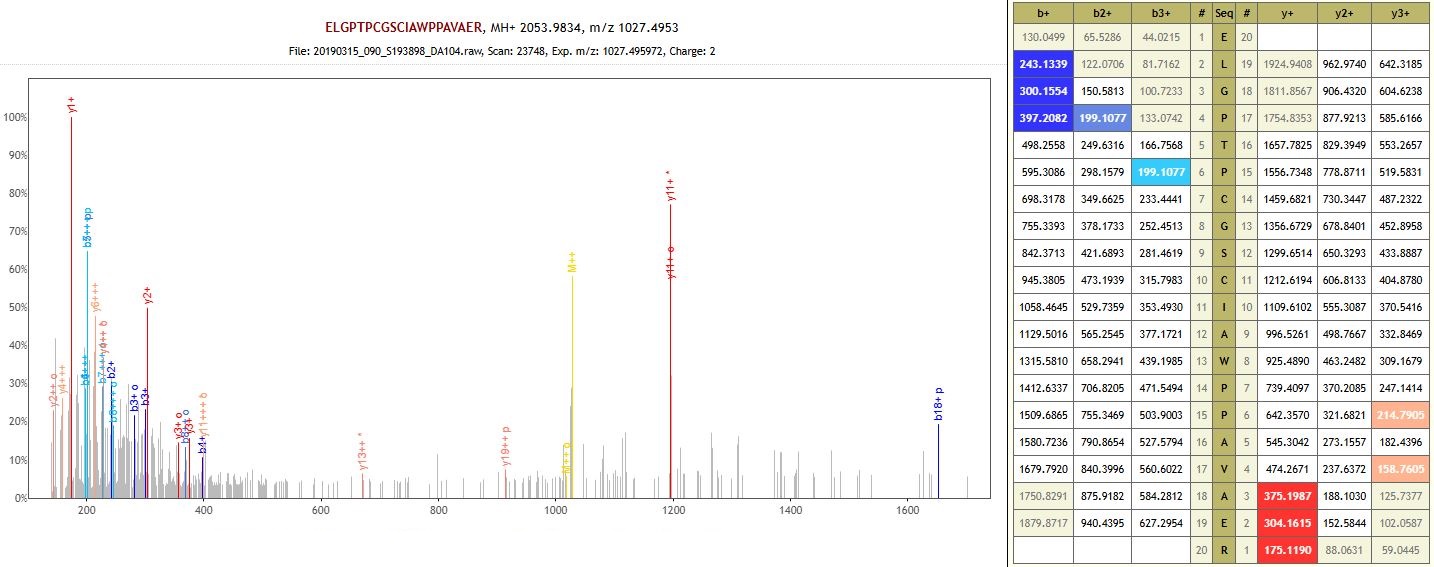

Protein: [RNA-polymerase]-subunit kinase (auto-enzyme)

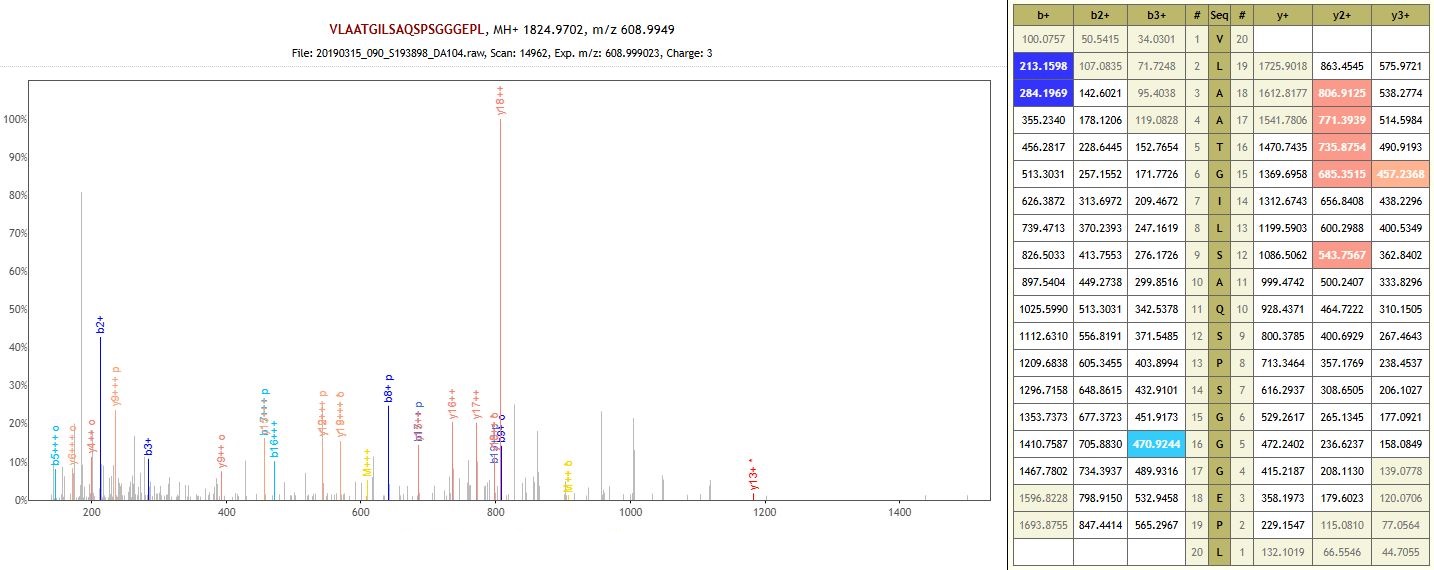
Protein: O-methyltransferase ZRP4-like (auto-enzyme)

**DA109 – Kalinovski 1 – Middle Bronze Age**

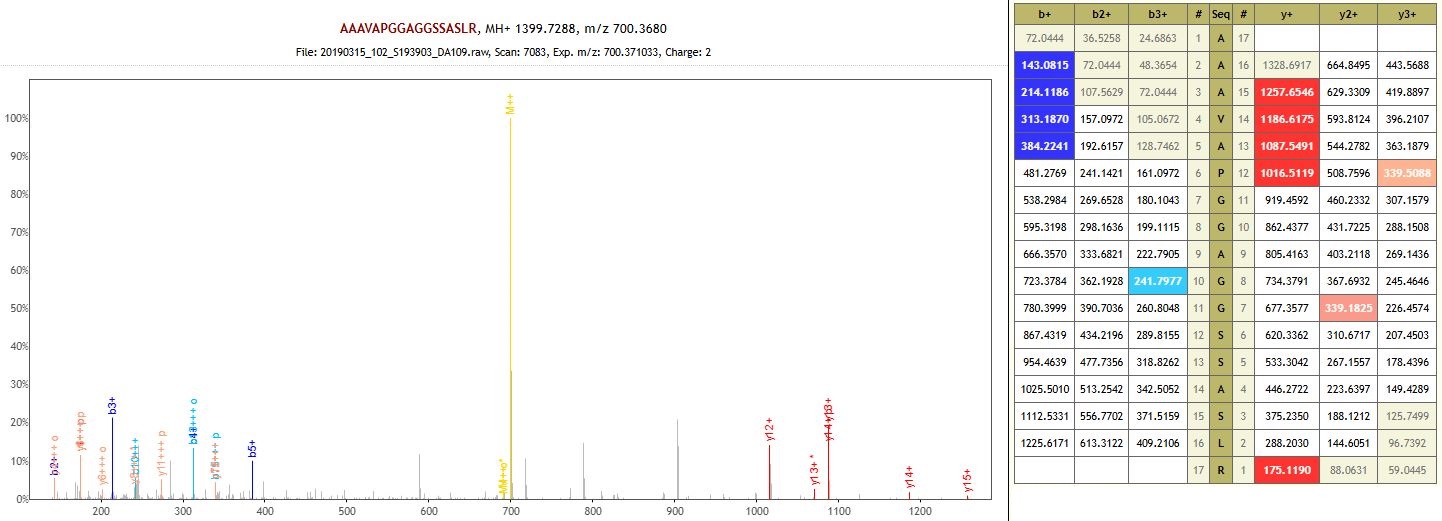

Protein: BHLH domain-containing protein (auto-enzyme)

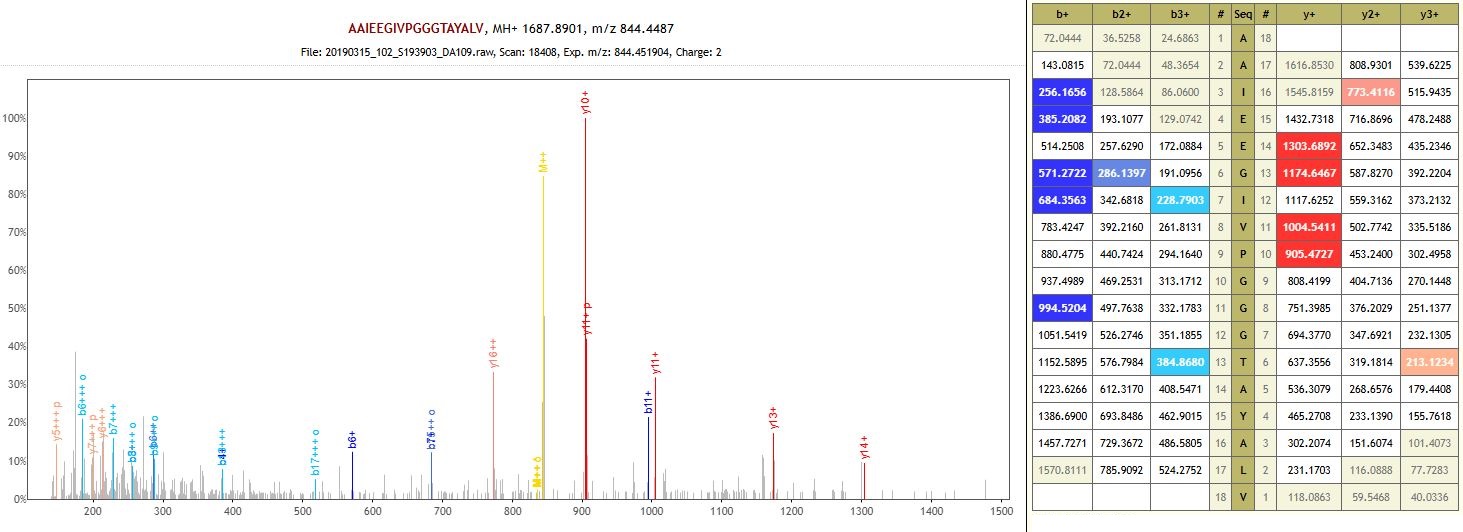

Protein: RuBisCO large subunit-binding protein subunit alpha, chloroplastic (non-specific enzyme)

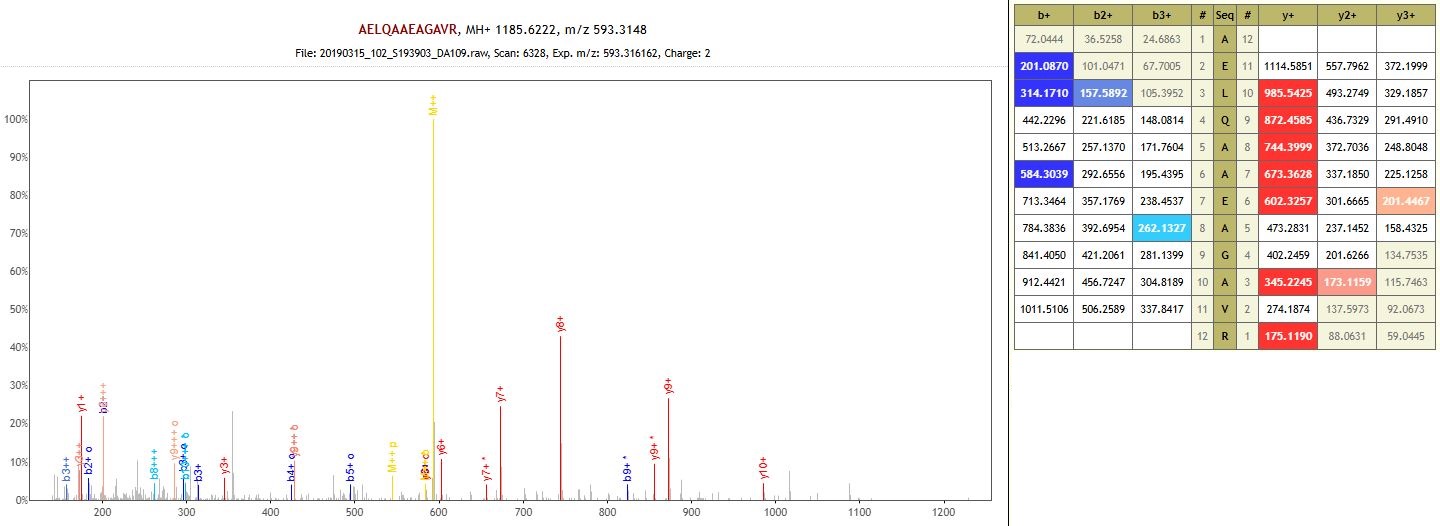

Protein: Pectinesterase (non-specific enzyme)

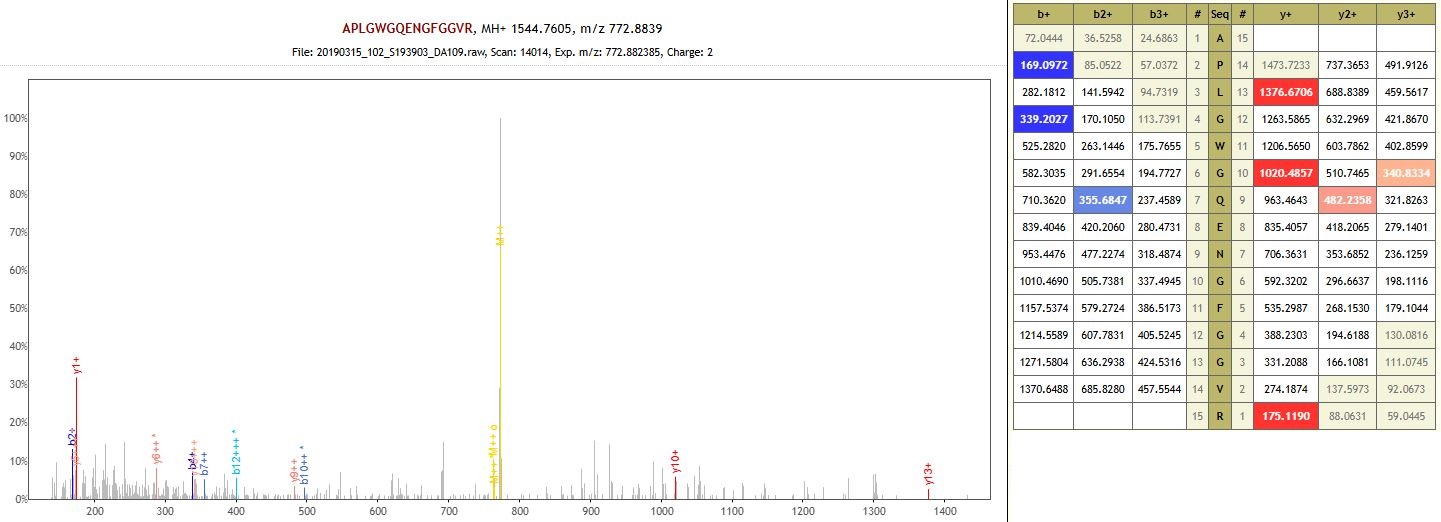

Protein: GATA-type domain-containing protein (non-specific enzyme)

**DA418 – Trudovoy 2 – Early Bronze Age**

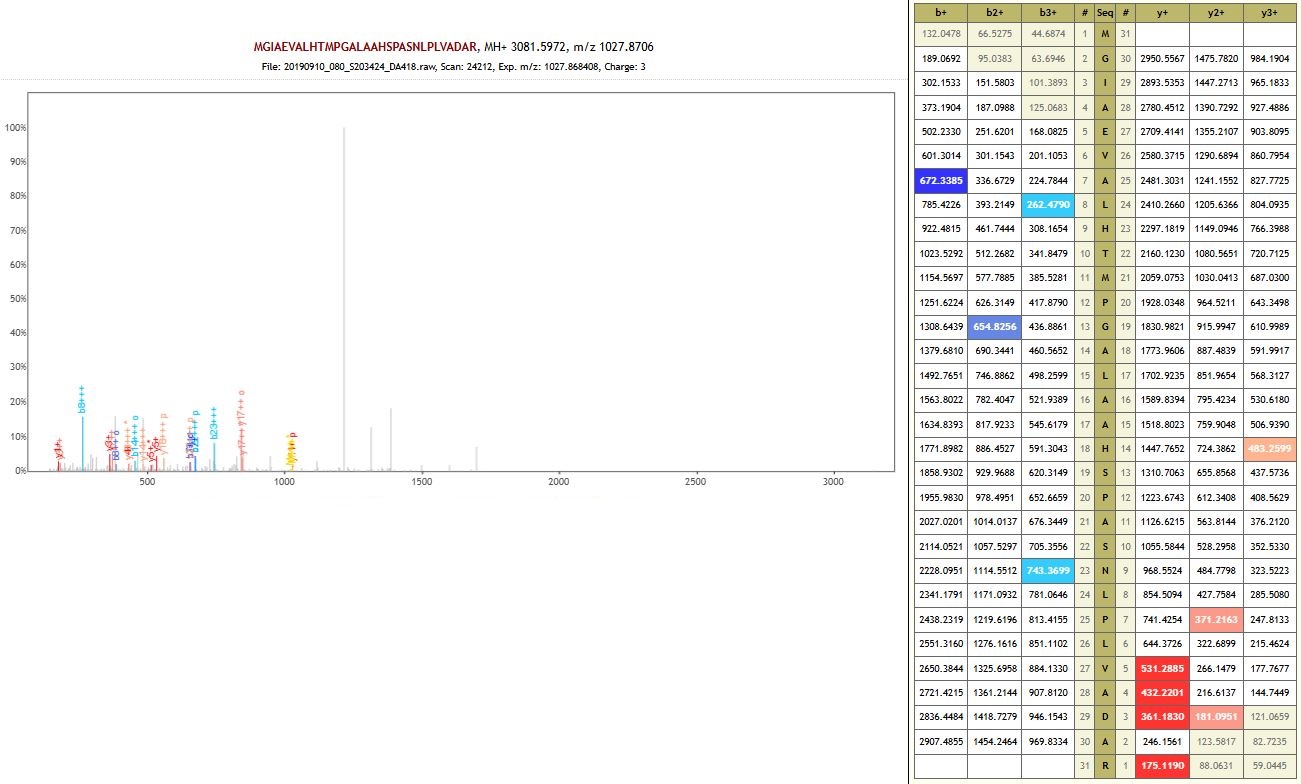

Protein: Alkaline/neutral invertase (auto-enzyme)

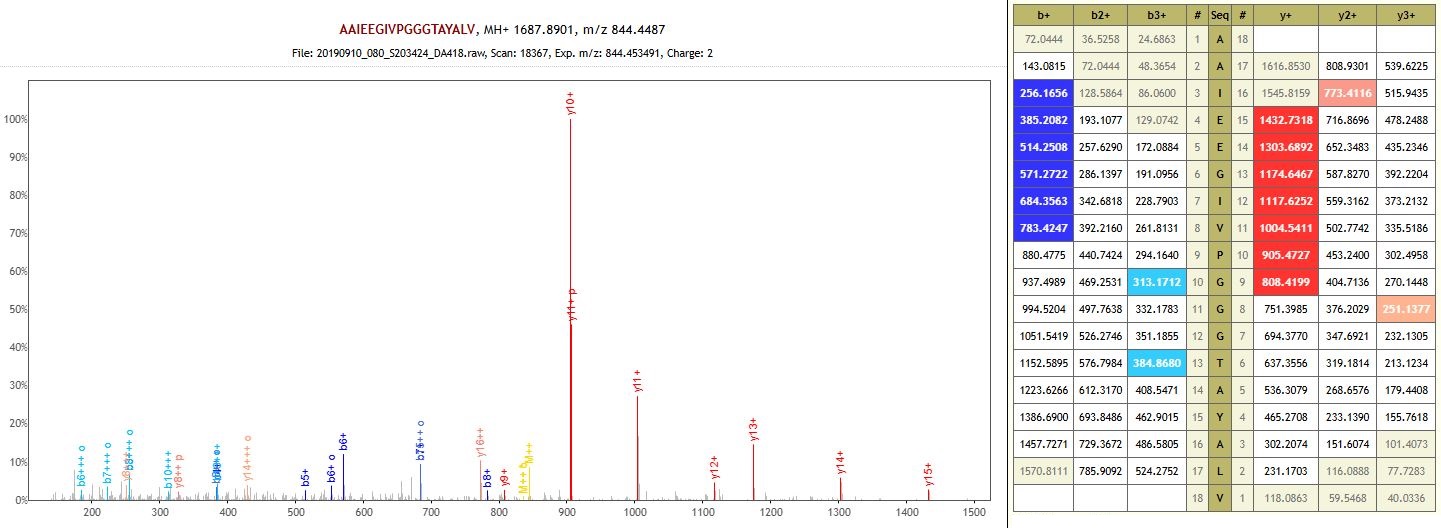

Protein: RuBisCO large subunit-binding protein subunit alpha, chloroplastic (non-specific enzyme)

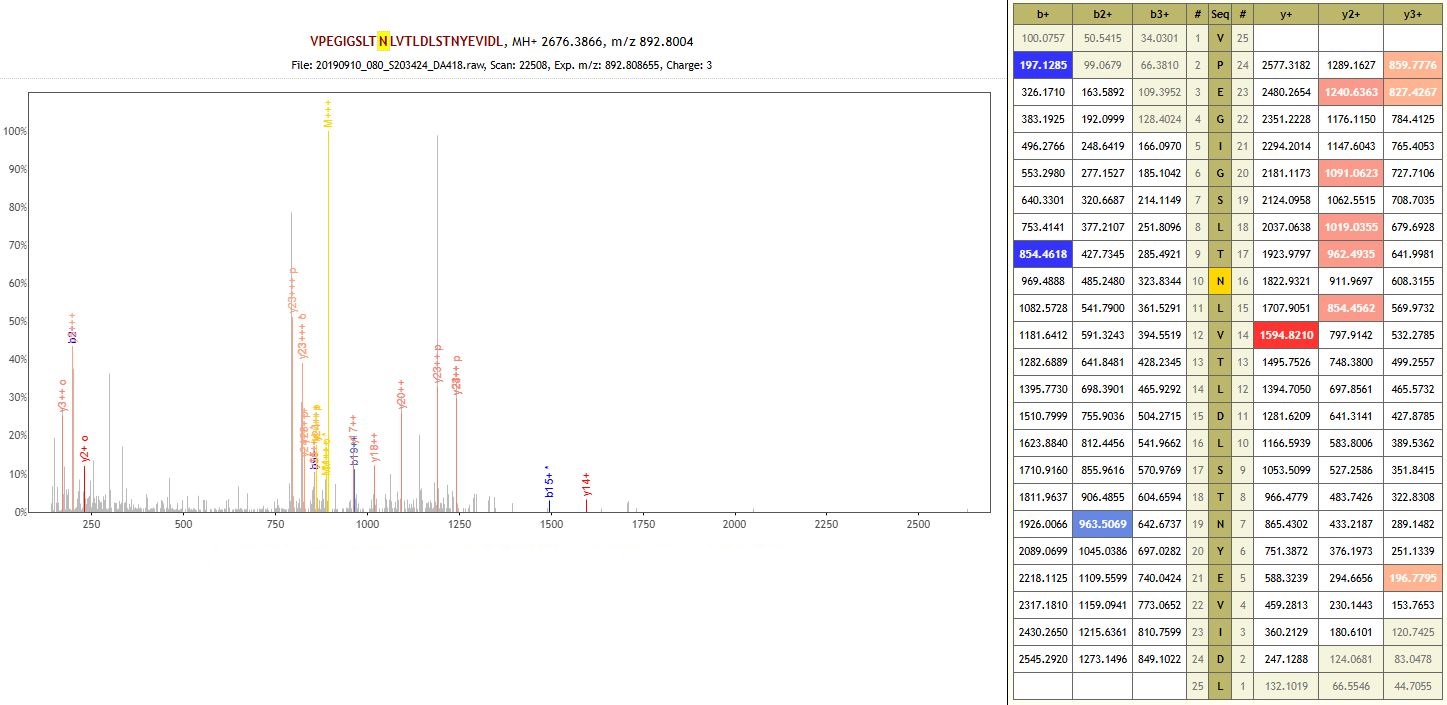

Protein: Leucine-rich repeat-containing N-terminal plant-type domain-containing protein (non-specific enzyme)

**DA420 – Krivyanskiy 9 – Early Bronze Age**

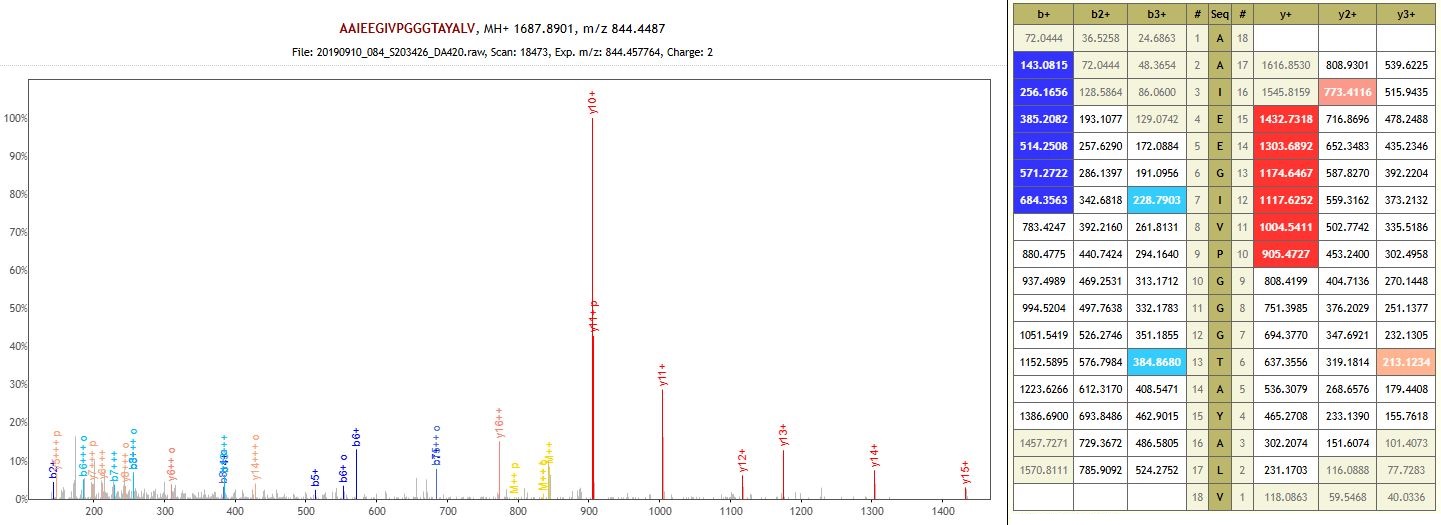

Protein: RuBisCO large subunit-binding protein subunit alpha, chloroplastic (auto-enzyme)

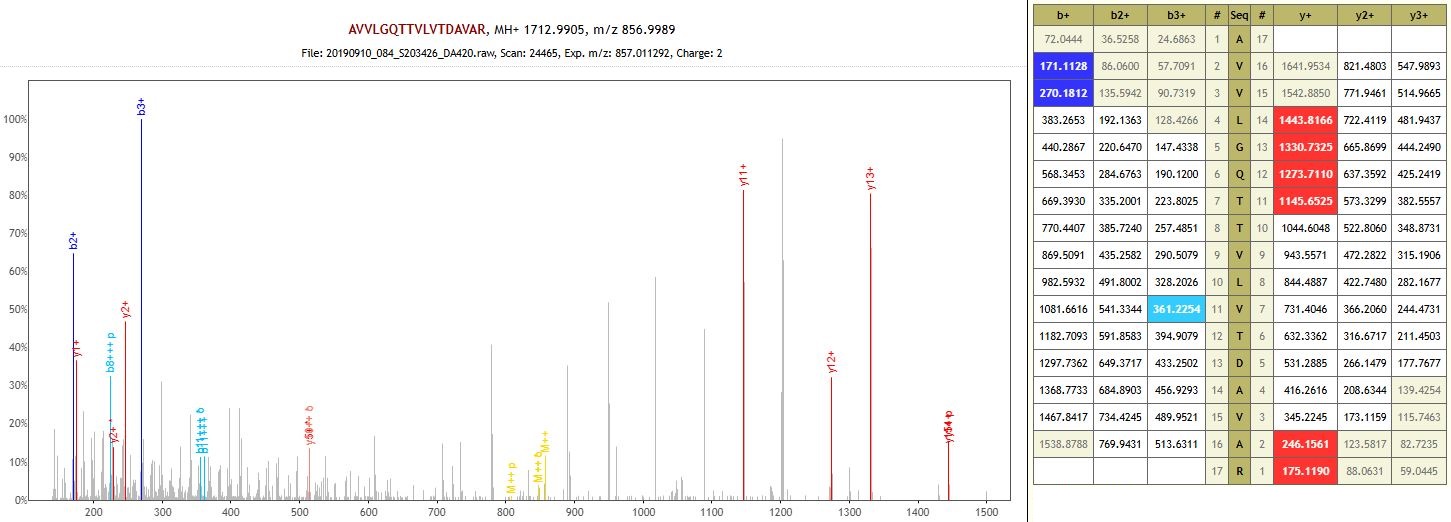

Protein: 4-coumarate--CoA ligase (auto-enzyme)

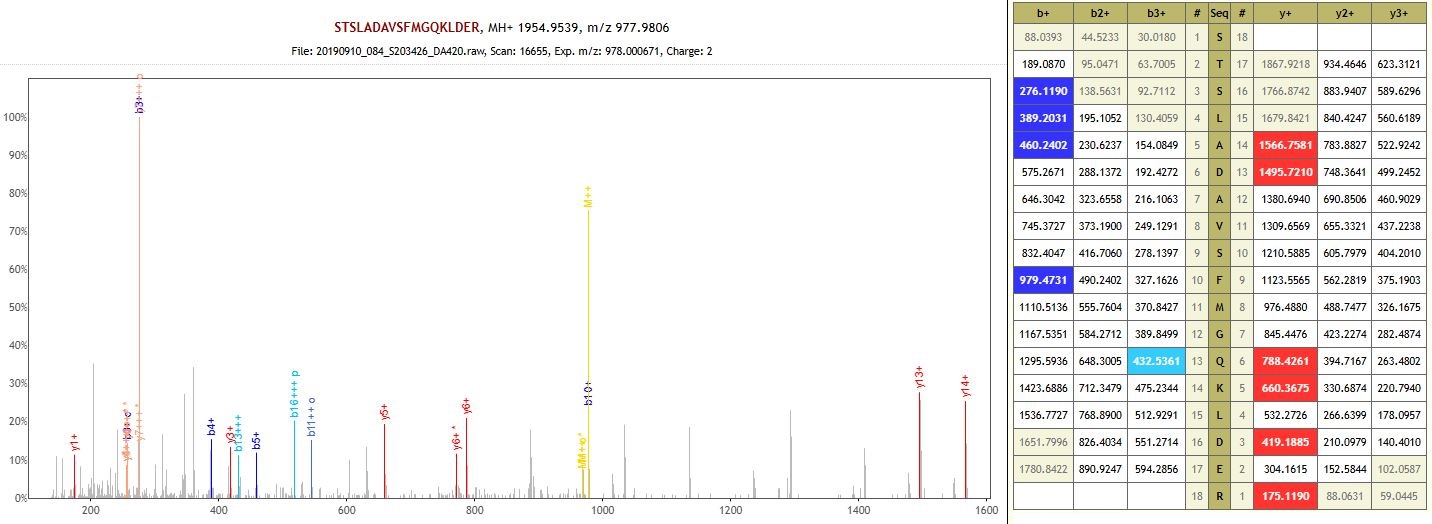

Protein: Ankyrin repeat protein SKIP35-like isoform X1 (auto-enzyme)

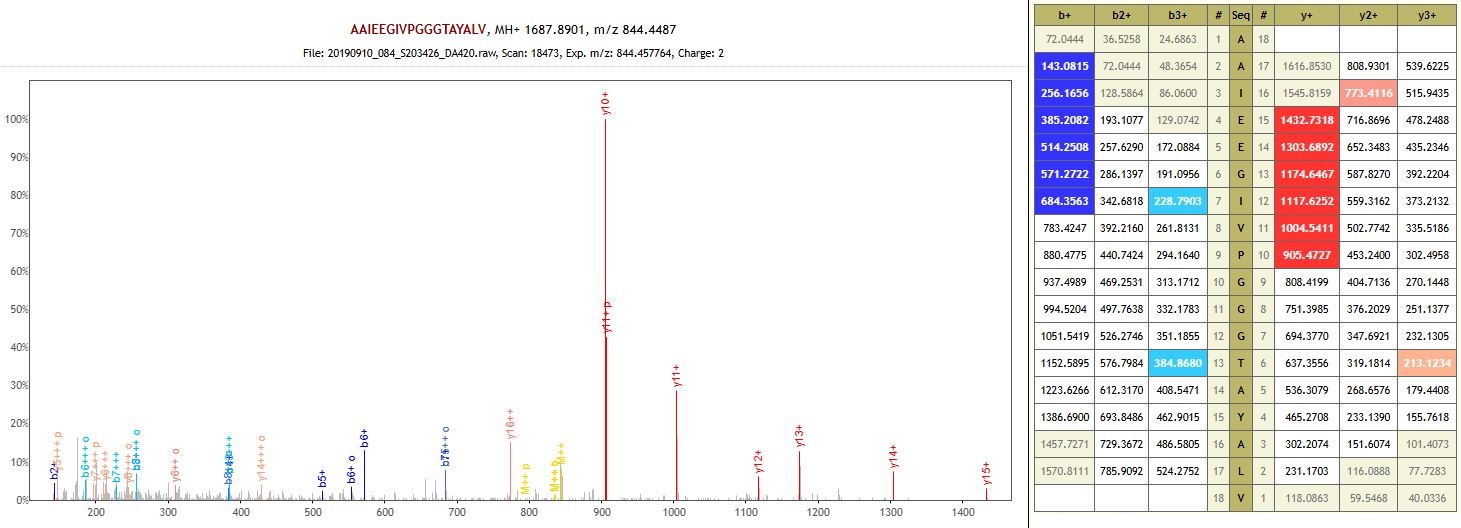

Protein: RuBisCO large subunit-binding protein subunit alpha, chloroplastic (non-specific enzyme)

**DA421 – Utevka 6 – Middle Bronze Age**

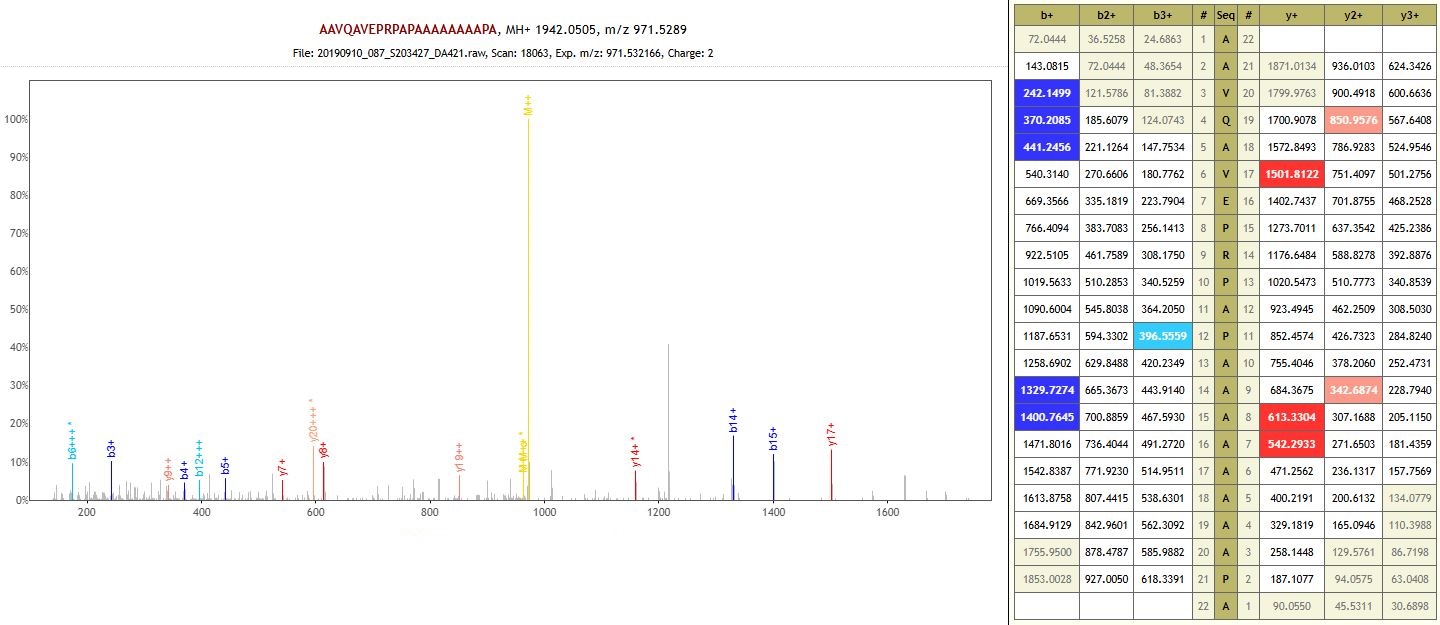

Protein: Calcium-binding mitochondrial carrier protein SCaMC-3-like (auto-enzyme)

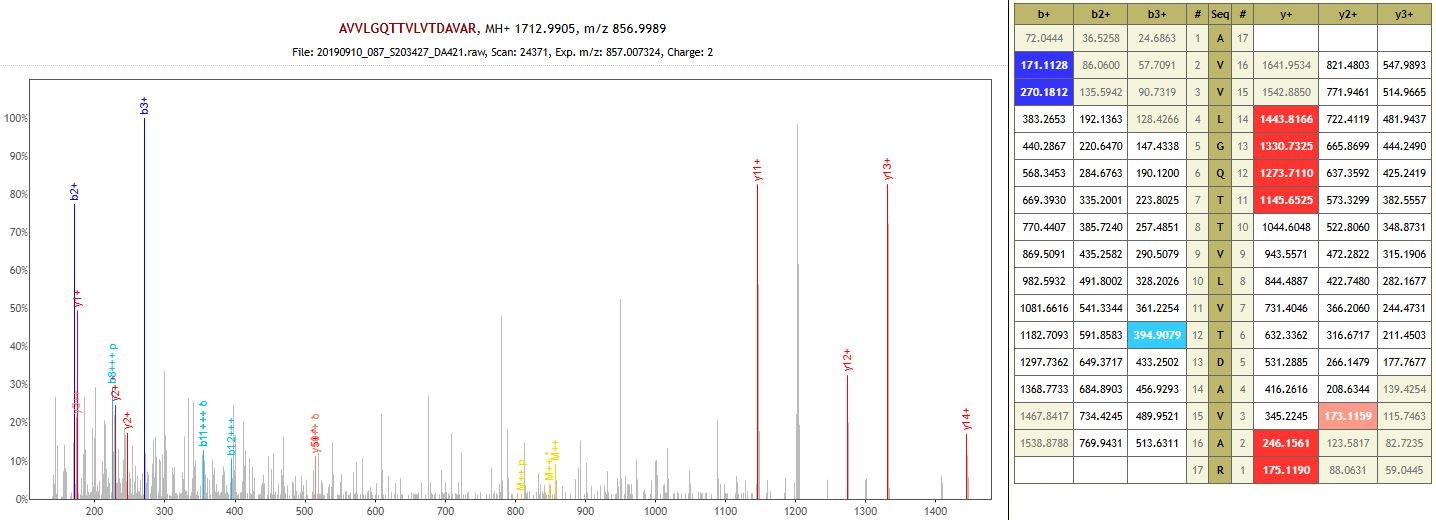

Protein: 4-coumarate--CoA ligase (auto-enzyme)

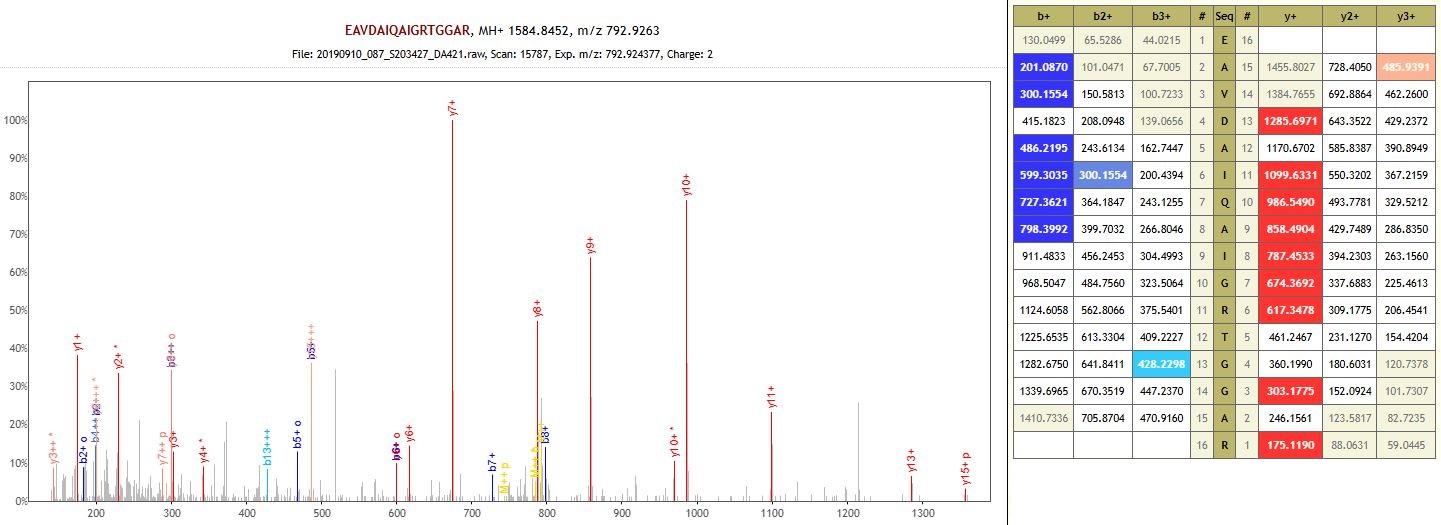

Protein: entatricopeptide repeat-containing protein (auto-enzyme)

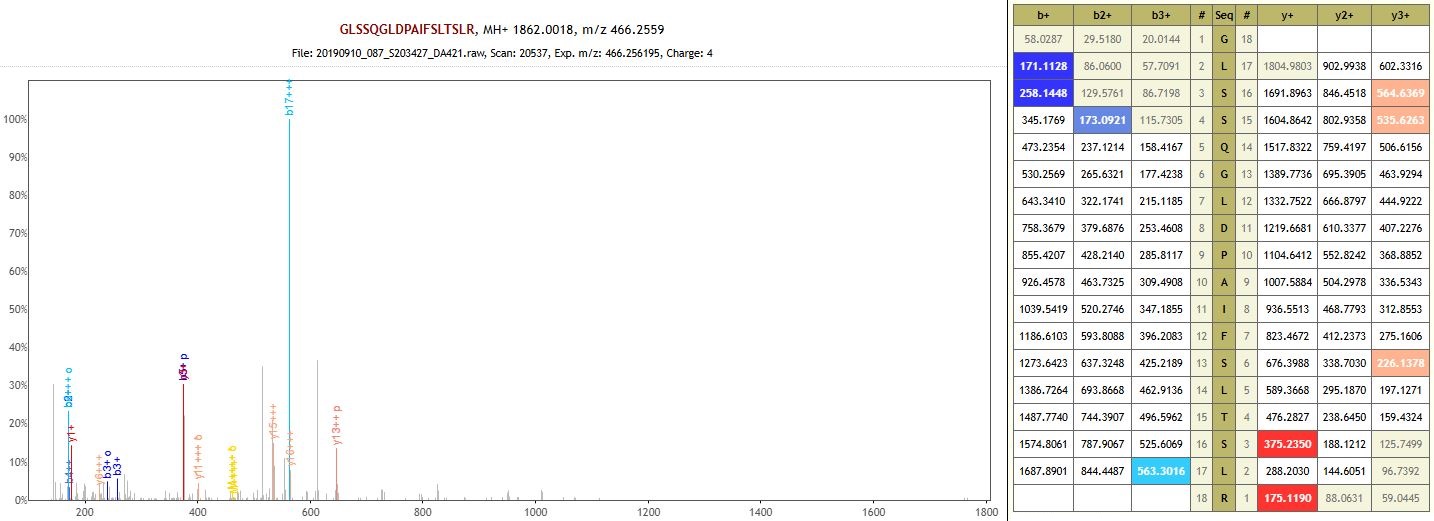

Protein: Receptor-like protein 12 isoform X2 (auto-enzyme)

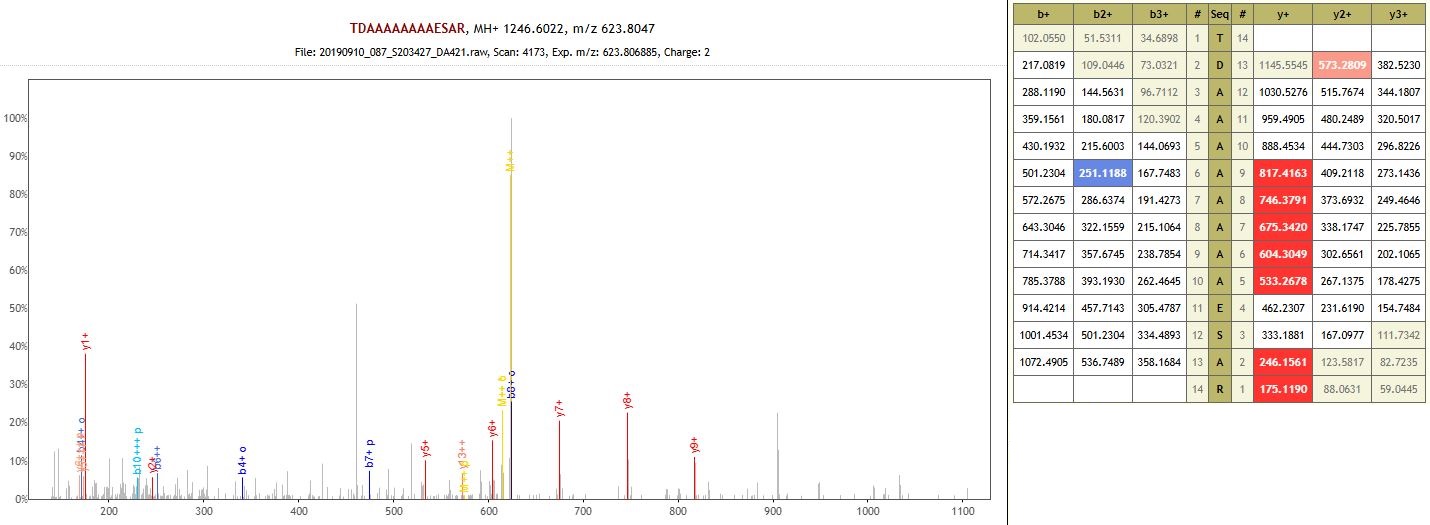

Protein: Embryonic protein DC-8-like (non-specific enzyme)

**DA424 – Khavalynsk 1 – Eneolithic**

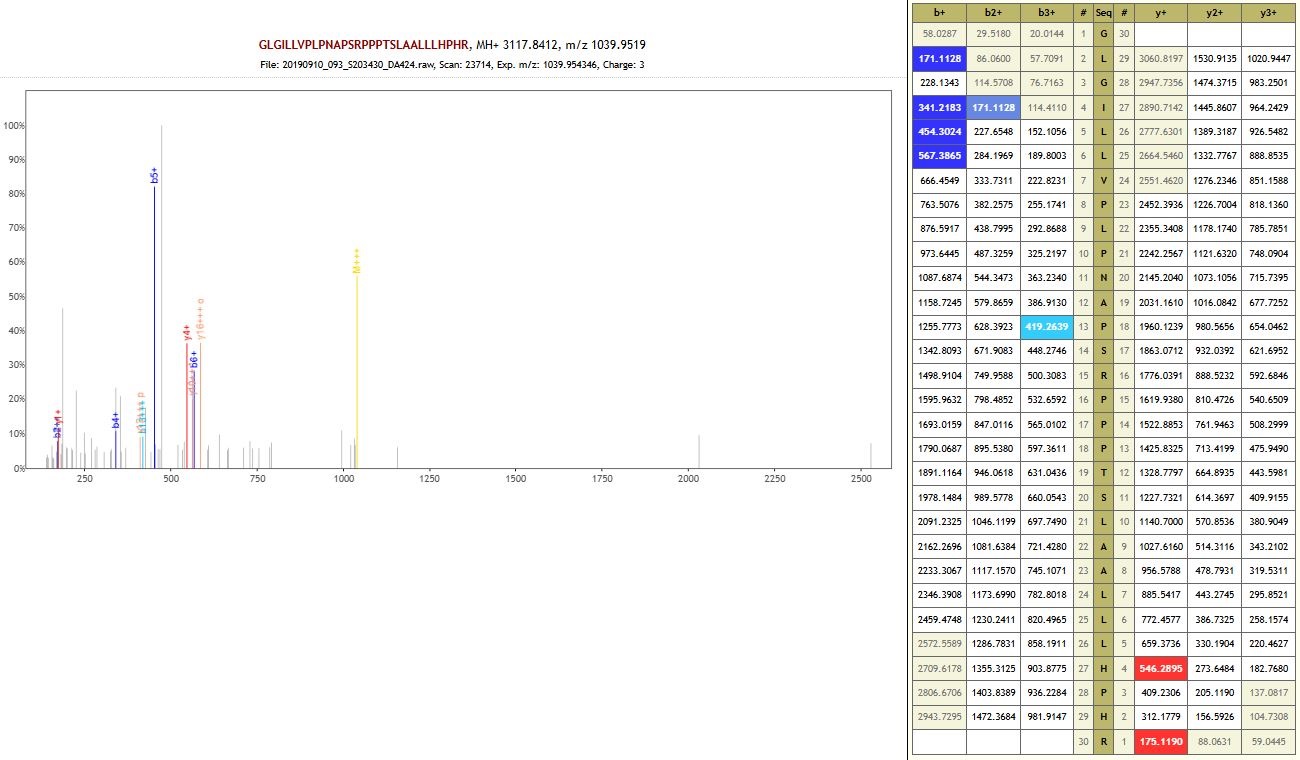

Protein: RNA helicase (auto-enzyme)

**DA425 – Panitskoe 6B – Early Bronze Age**

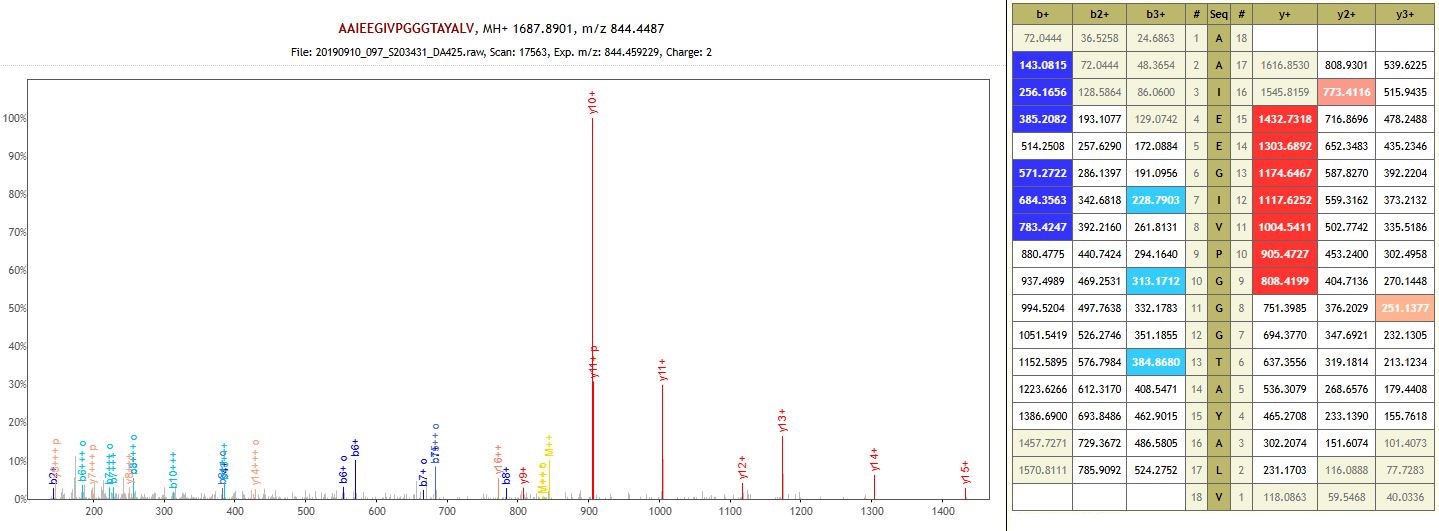

Protein: RuBisCO large subunit-binding protein subunit alpha, chloroplastic (auto-enzyme)

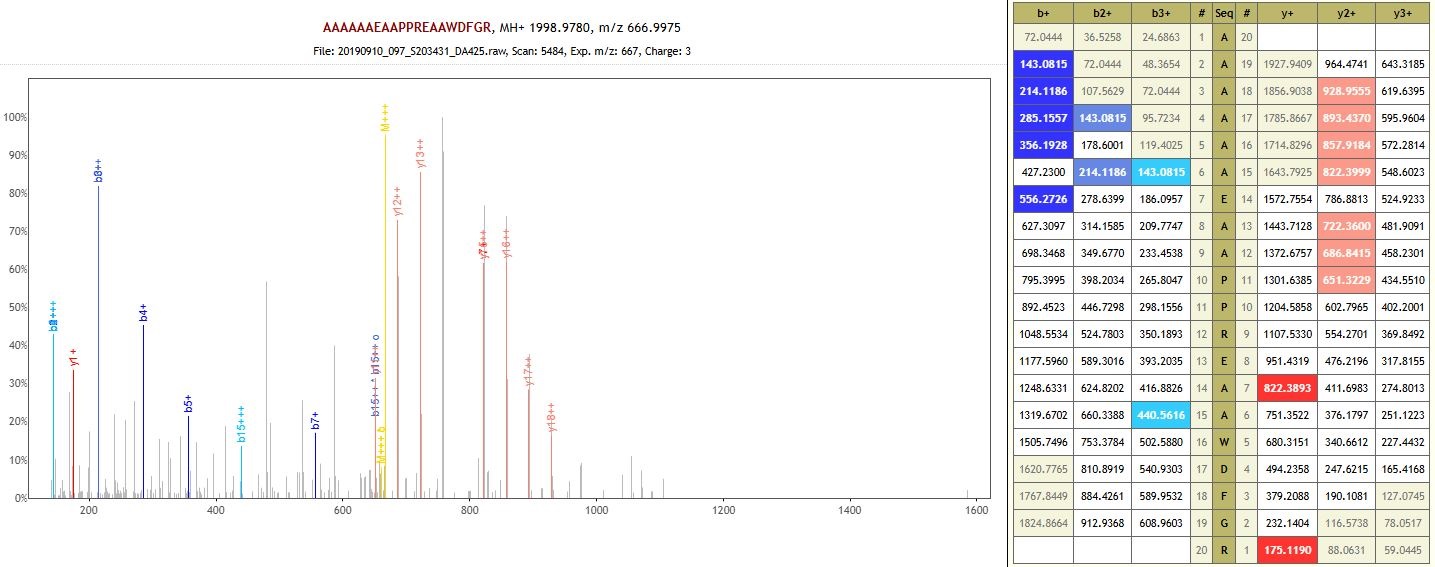

Protein: Uncharacterized protein (non-specific enzyme)

**DA426 – Murziha 2 – Eneolithic**

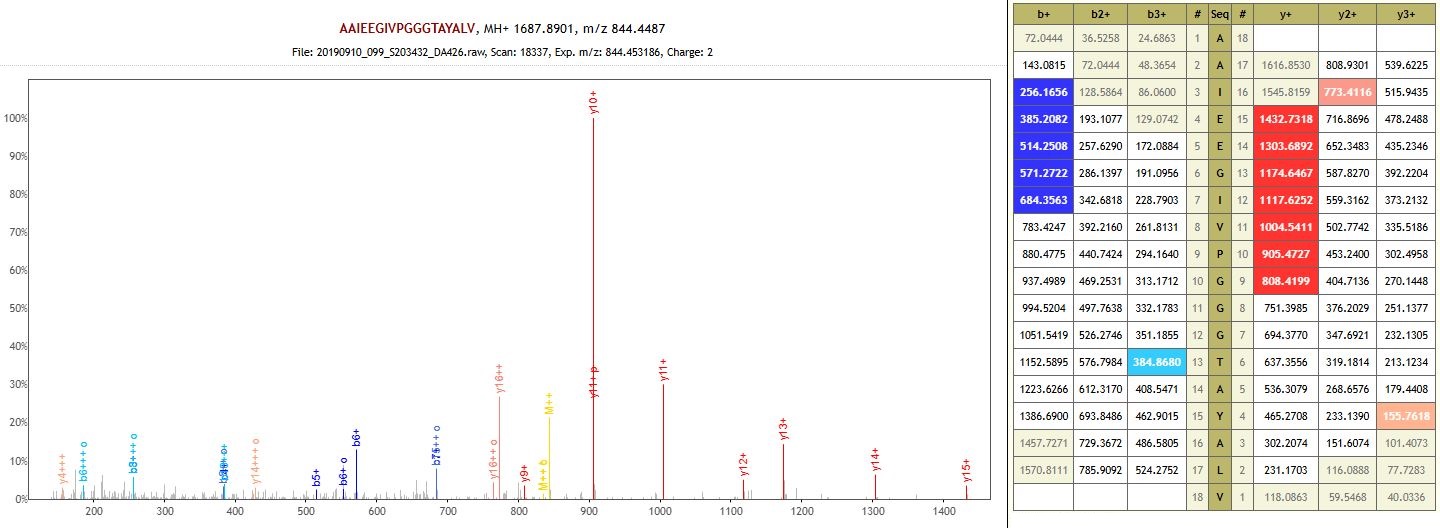

Protein: RuBisCO large subunit-binding protein subunit alpha, chloroplastic (auto-enzyme)

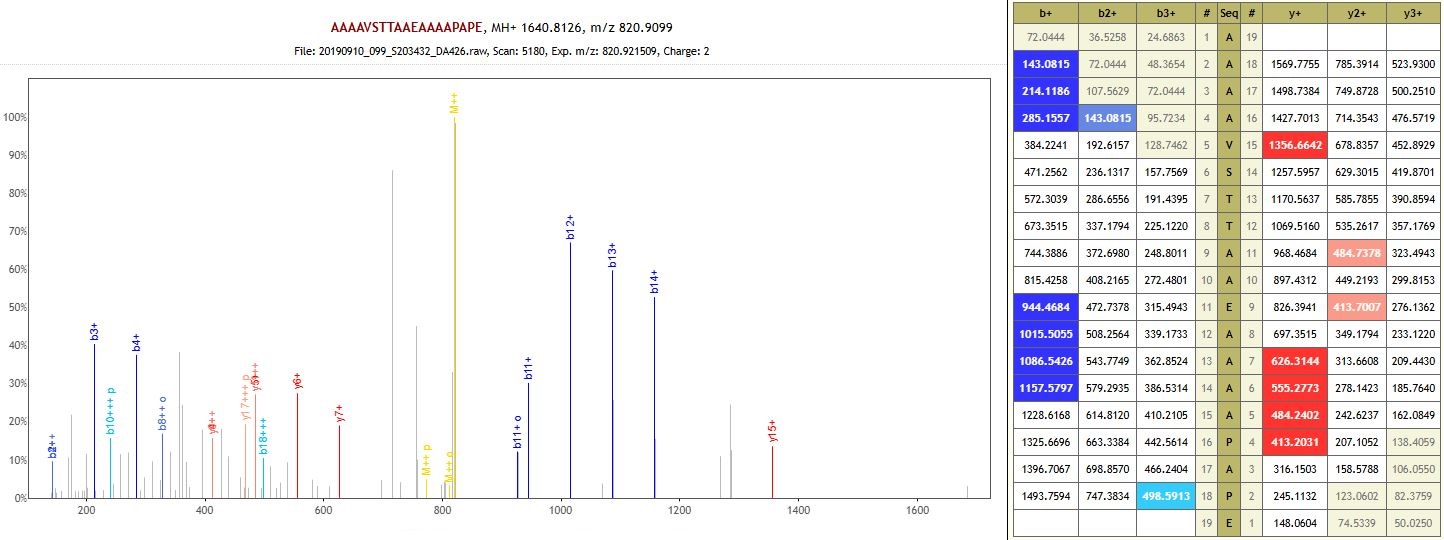

Protein: Enolase-phosphatase E1-like (non-specific enzyme)

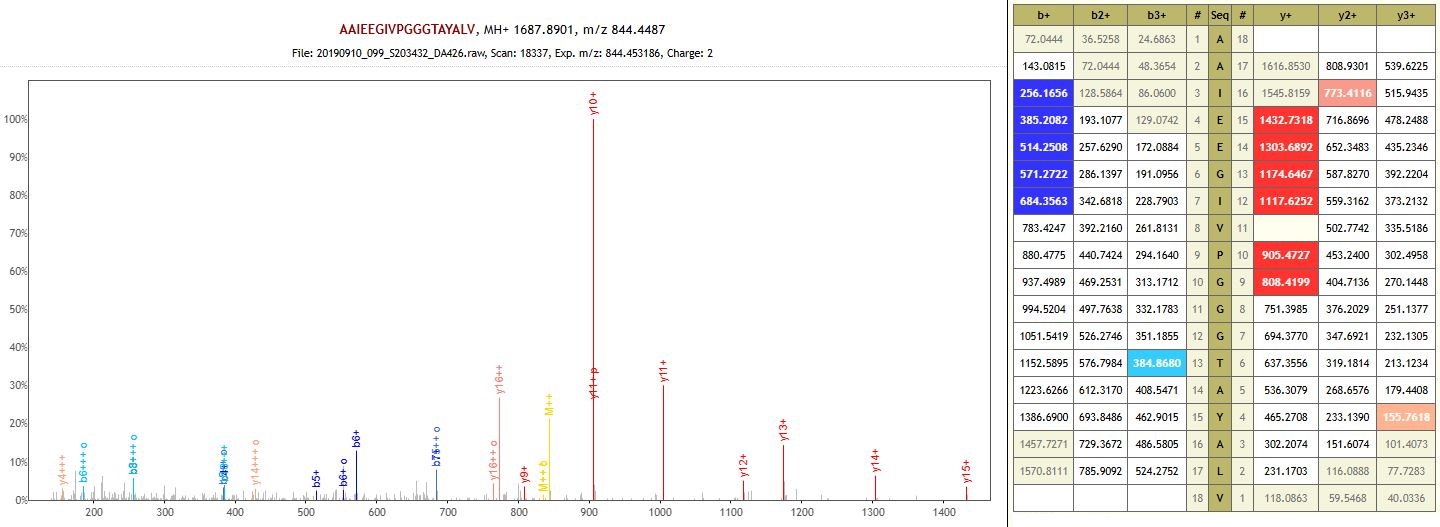

Protein: RuBisCO large subunit-binding protein subunit alpha, chloroplastic (non-specific enzyme)

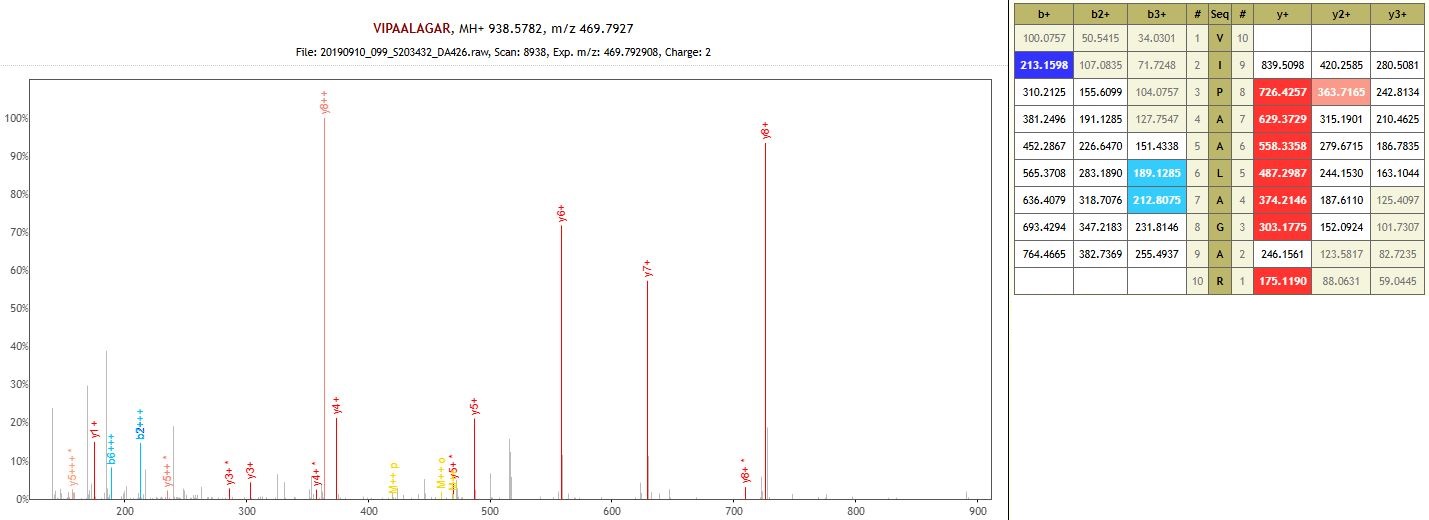

Protein: non-specific serine/threonine protein kinase (non-specific enzyme)

**DA427 – Pyatiletka – Early Bronze Age**

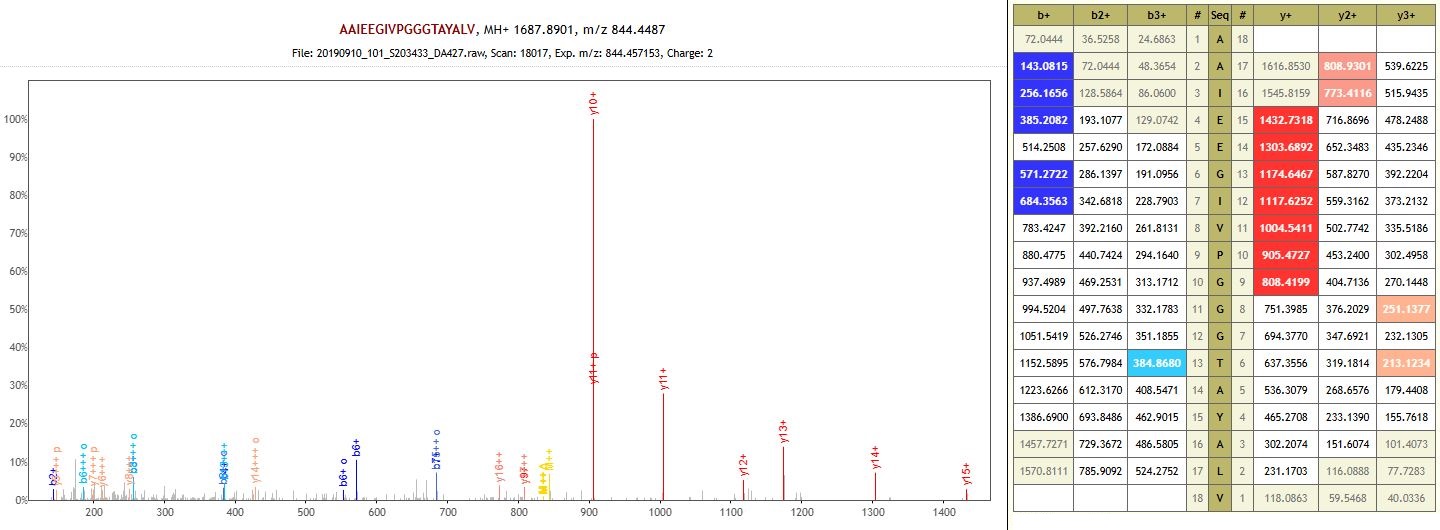

Protein: RuBisCO large subunit-binding protein subunit alpha, chloroplastic (auto-enzyme)

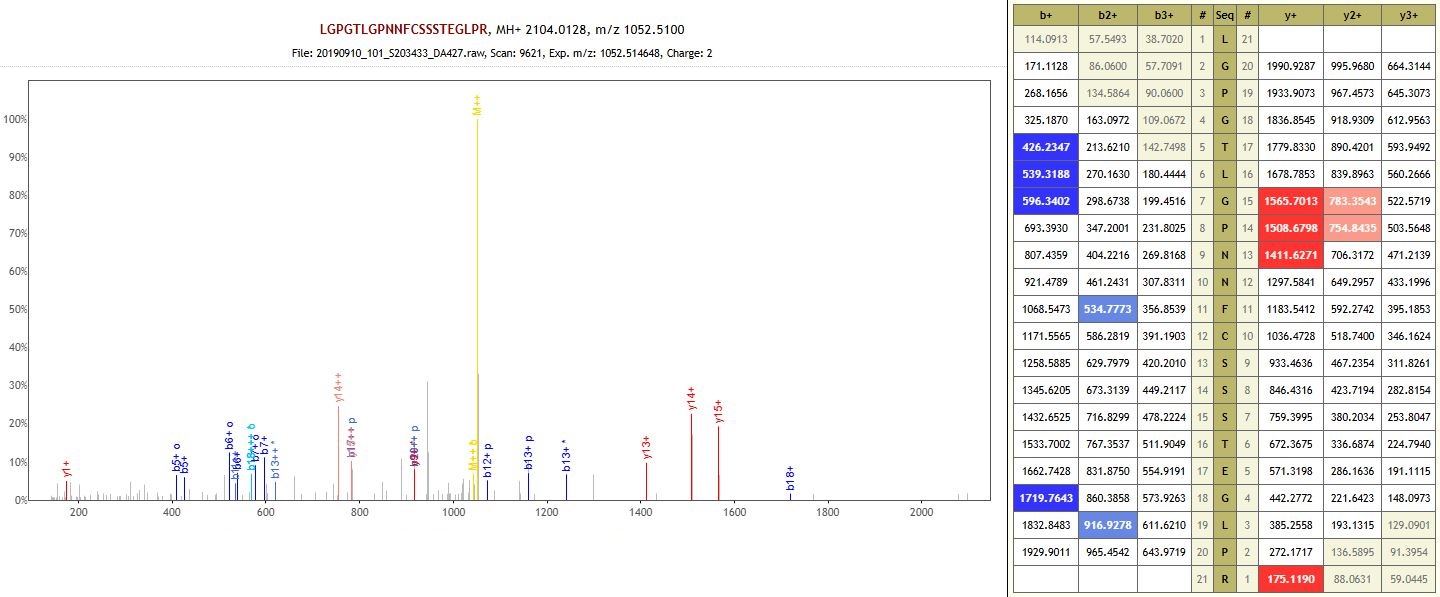

Protein: Uncharacterized protein (non-specific enzyme)

**DA429 – Leshchevskoe 1 – Early Bronze Age**

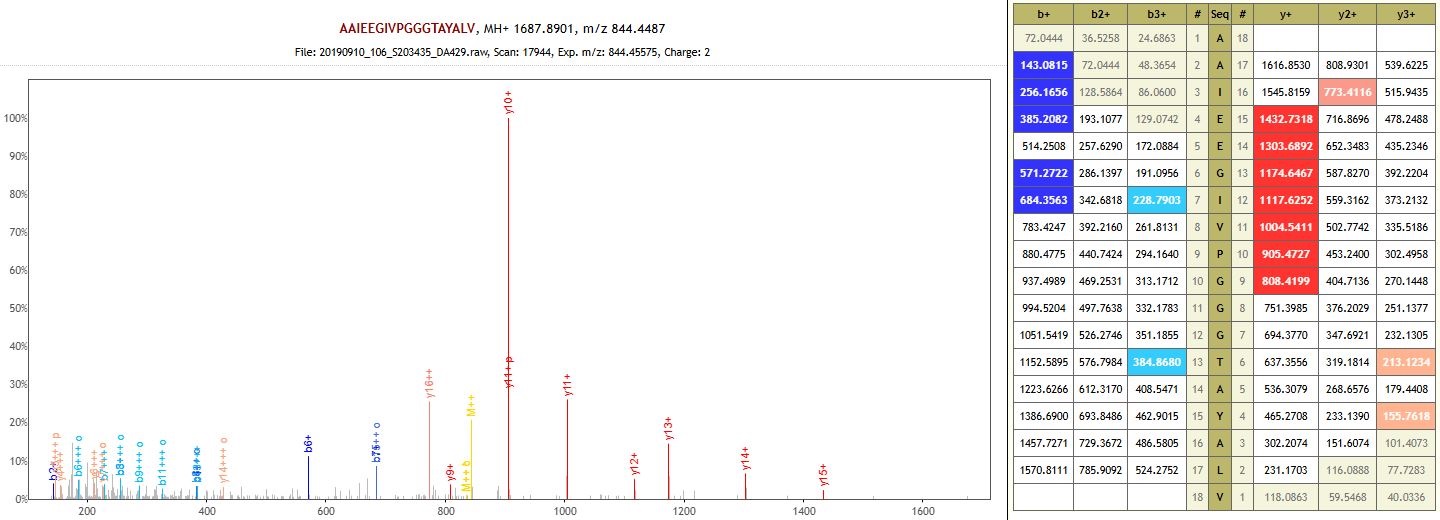

Protein: RuBisCO large subunit-binding protein subunit alpha, chloroplastic (non-specific enzyme)

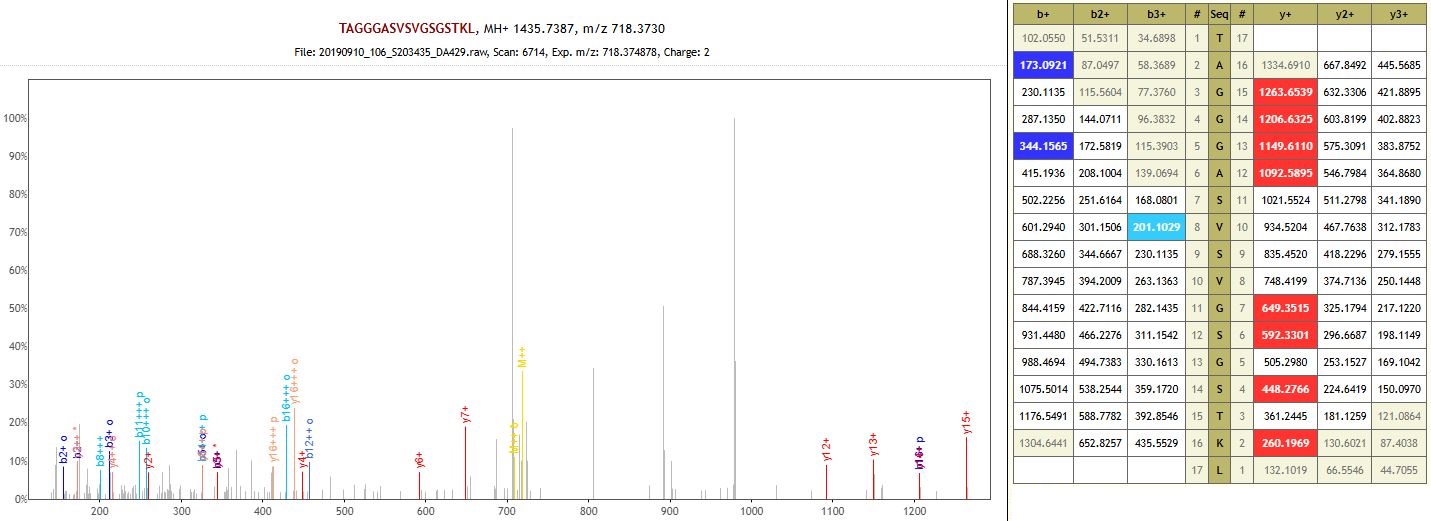

Protein: Protein IRX15-LIKE-like (non-specific enzyme)

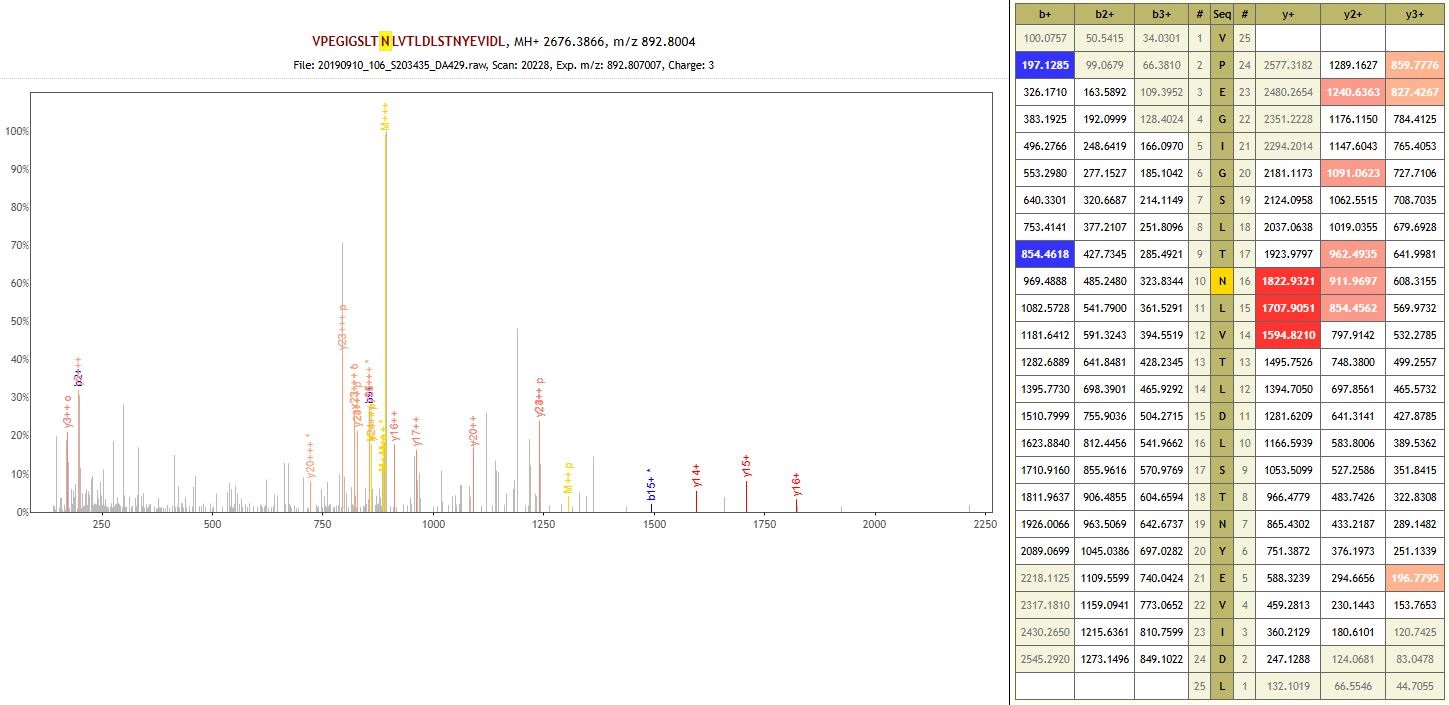

Protein: Leucine-rich repeat-containing N-terminal plant-type domain-containing protein (non-specific enzyme)

**DA430 – Khavalynsk 2 – Eneolithic**

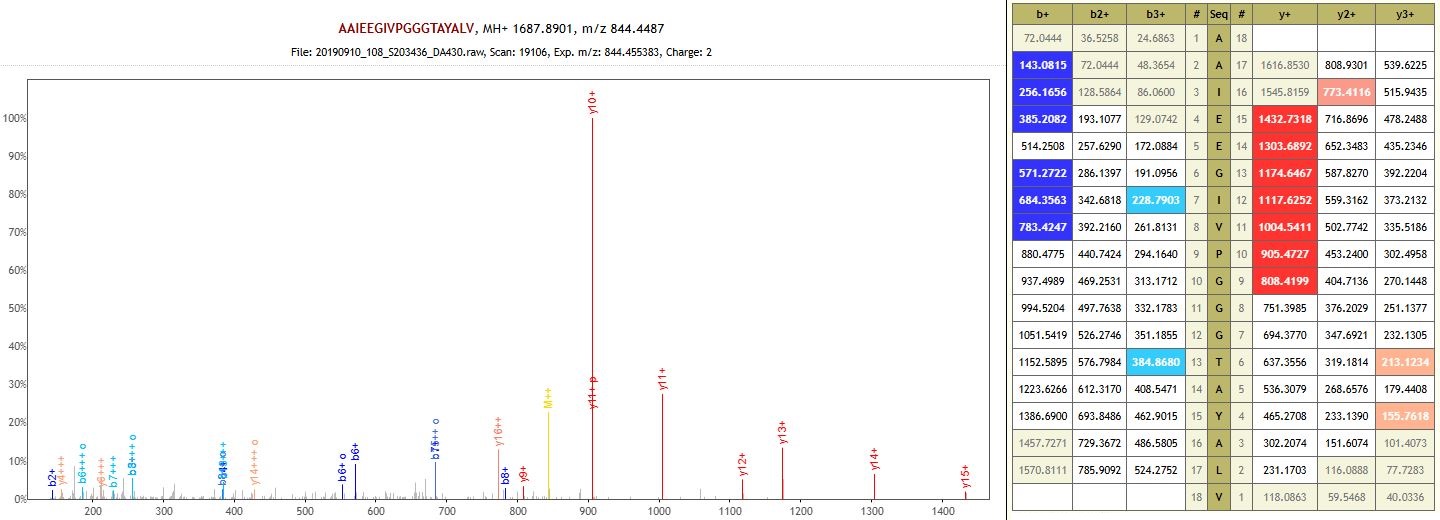

Protein: RuBisCO large subunit-binding protein subunit alpha, chloroplastic (auto-enzyme)

**DA432 – Lopatino 2 – Final Bronze Age**

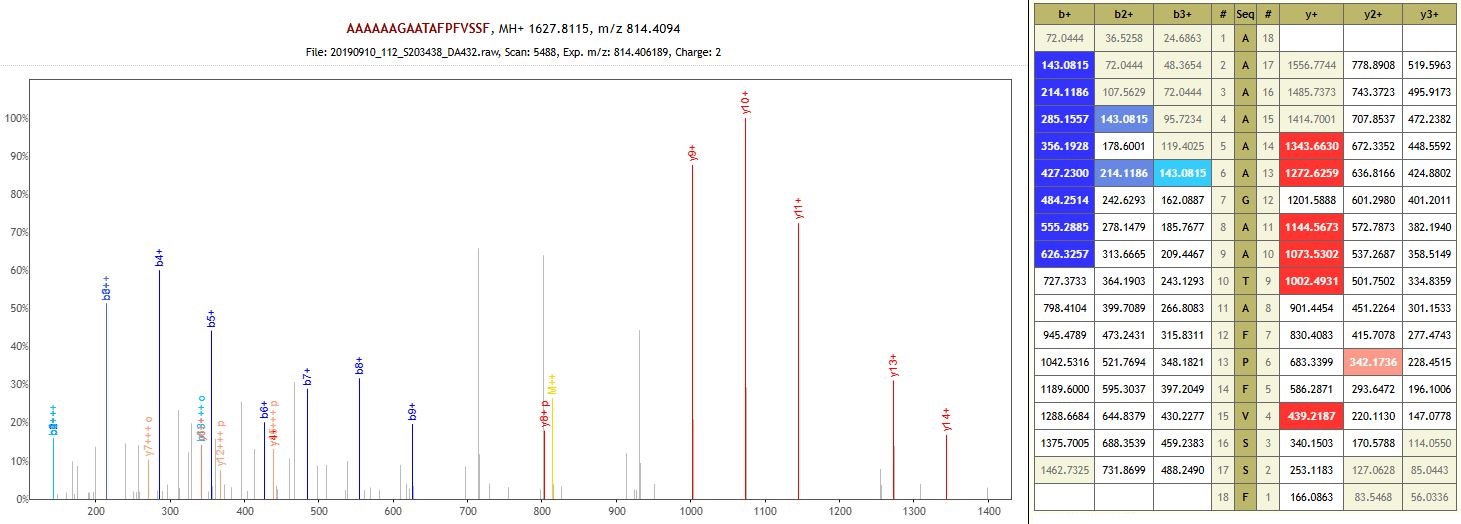

Protein: Peptidoglycan binding-like domain-containing protein (non-specific enzyme)

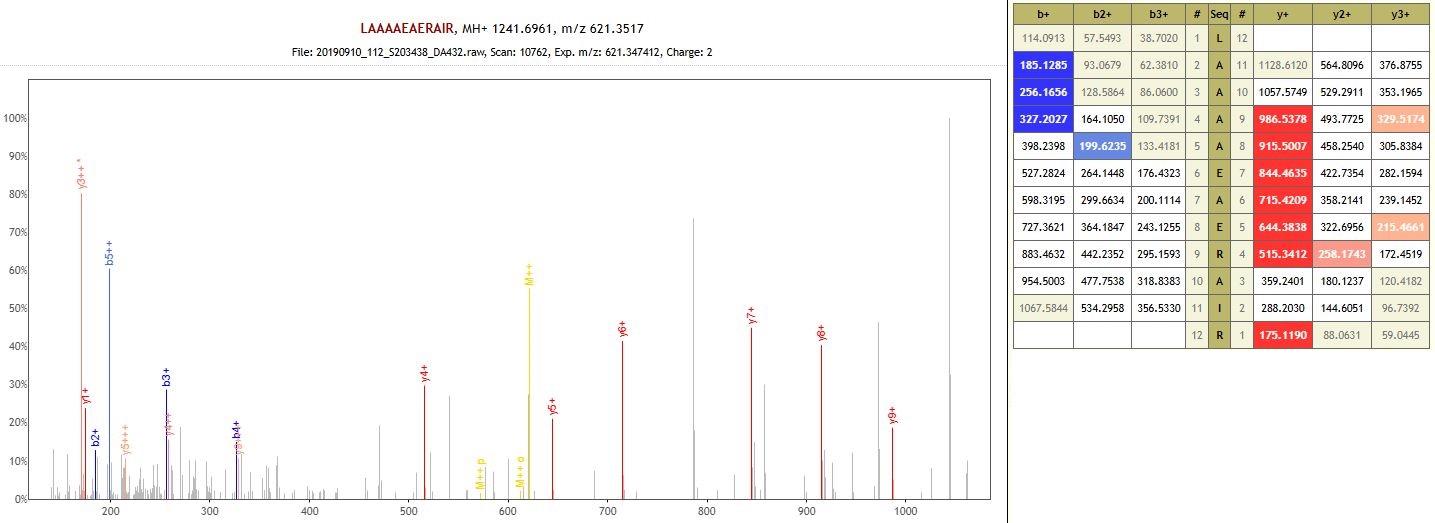

Protein: Biotin synthase (non-specific enzyme)

**DA433 – Mustayevo 5 – Early Bronze Age**

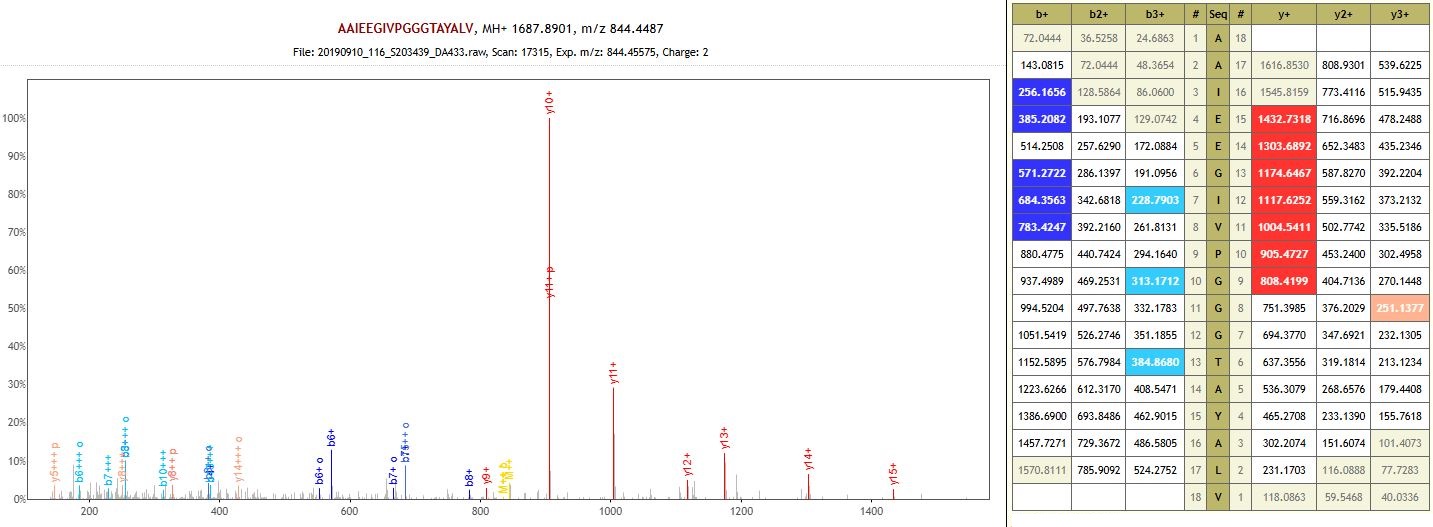

Protein: RuBisCO large subunit-binding protein subunit alpha, chloroplastic (non-specific enzyme)

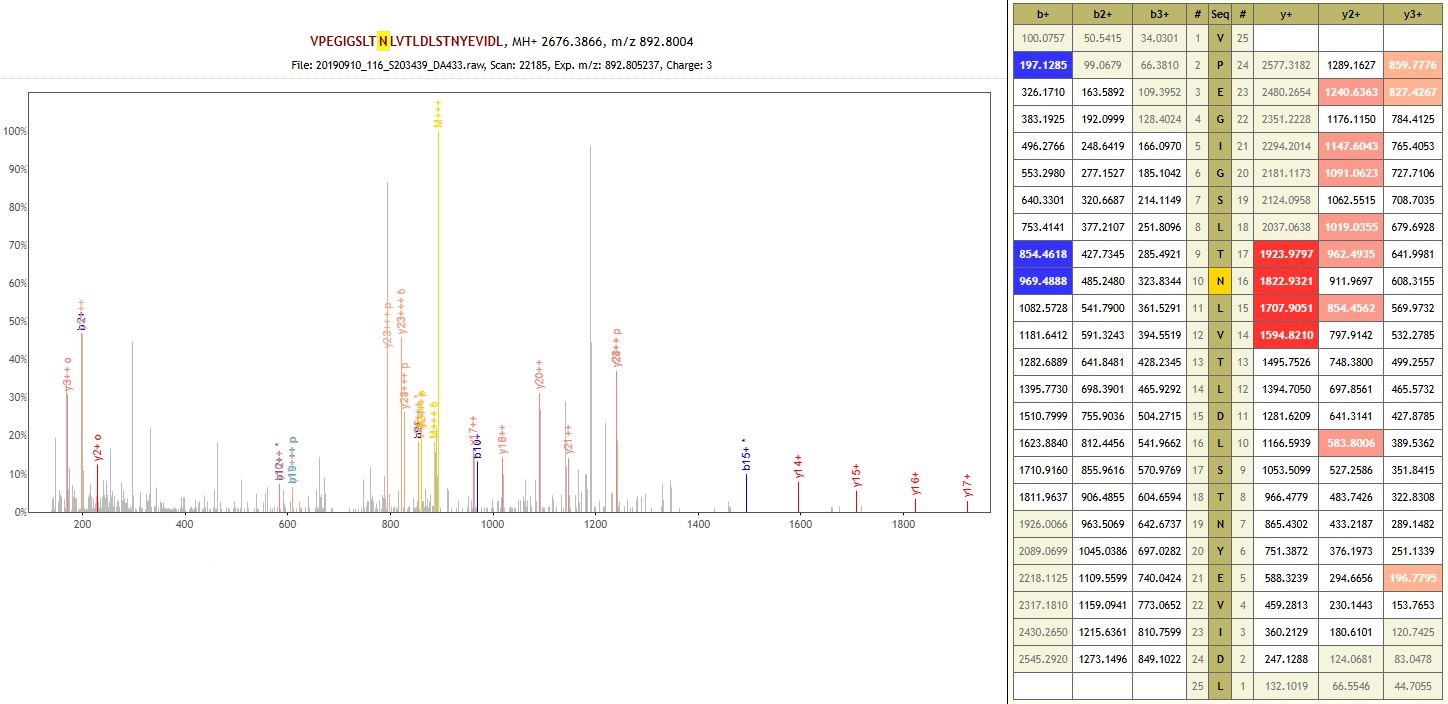

Protein: Leucine-rich repeat-containing N-terminal plant-type domain-containing protein (non-specific enzyme)

**DA434 – Shumayevo 2 – Middle Bronze Age**

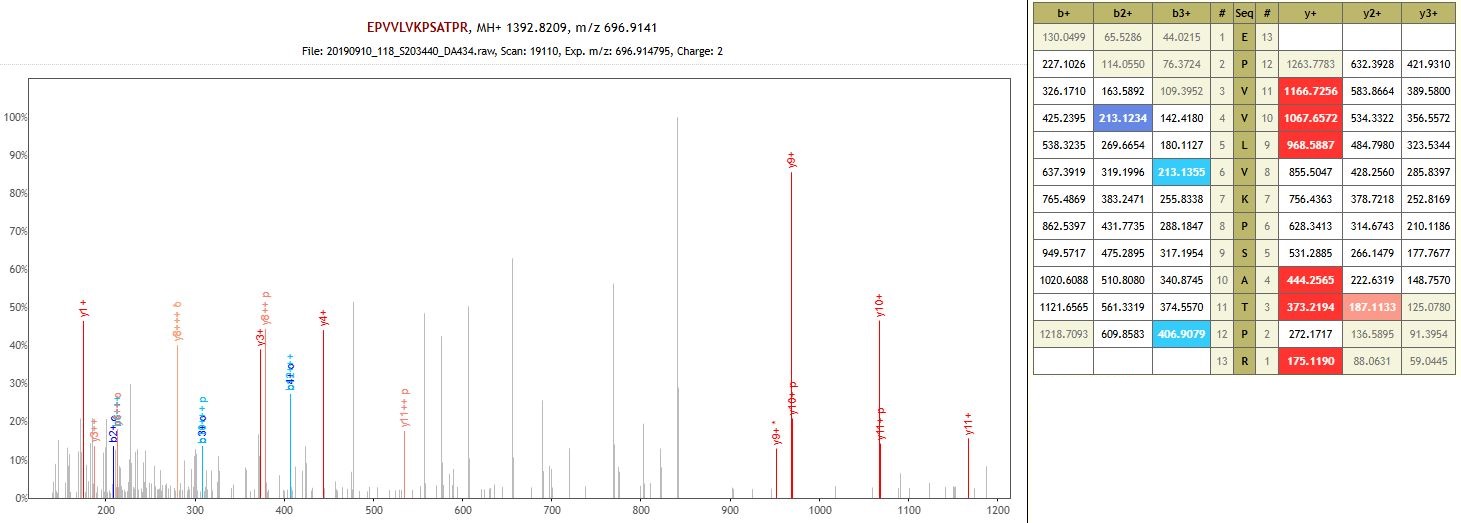

Protein: Patatin (auto-enzyme)

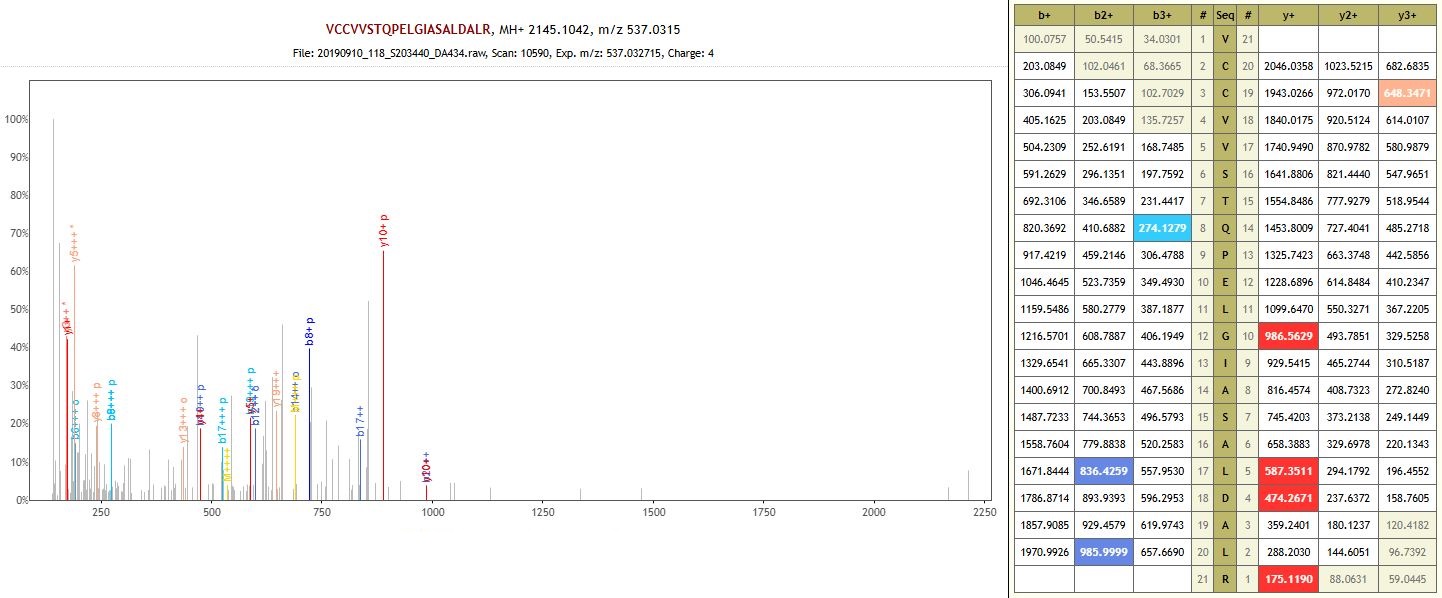

Protein: Uncharacterized protein (auto-enzyme)

**DA435 – Podlesny – Early Bronze Age**

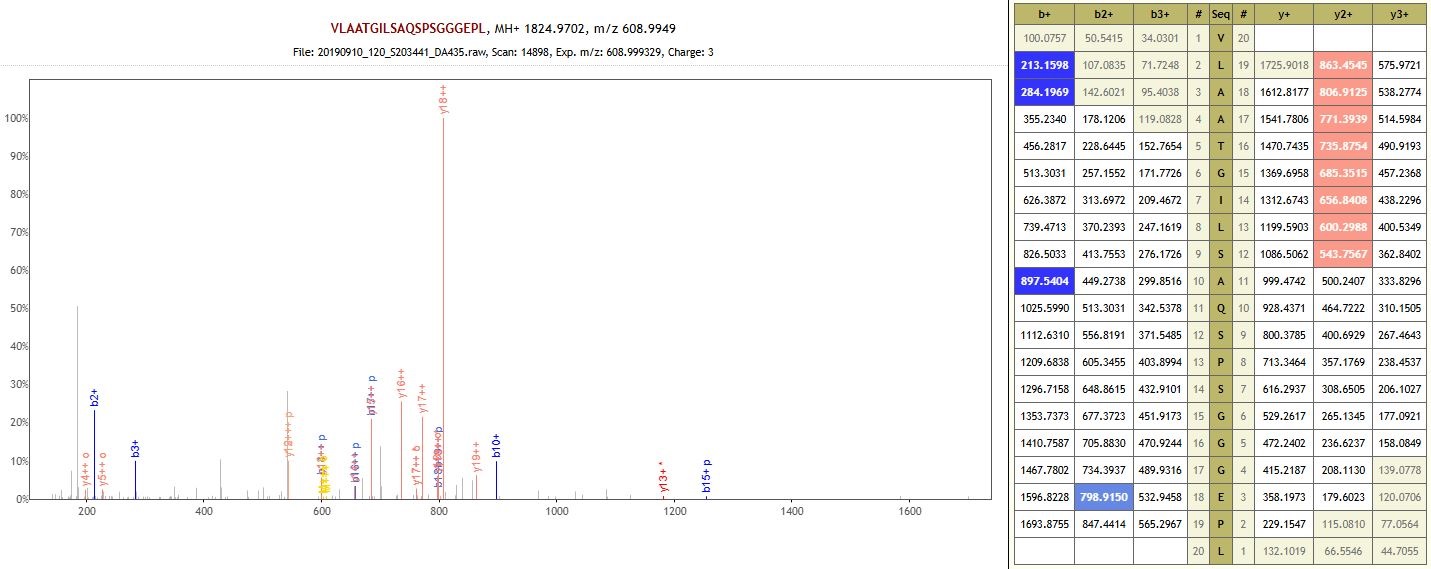

Protein: O-methyltransferase ZRP4-like (auto-enzyme)

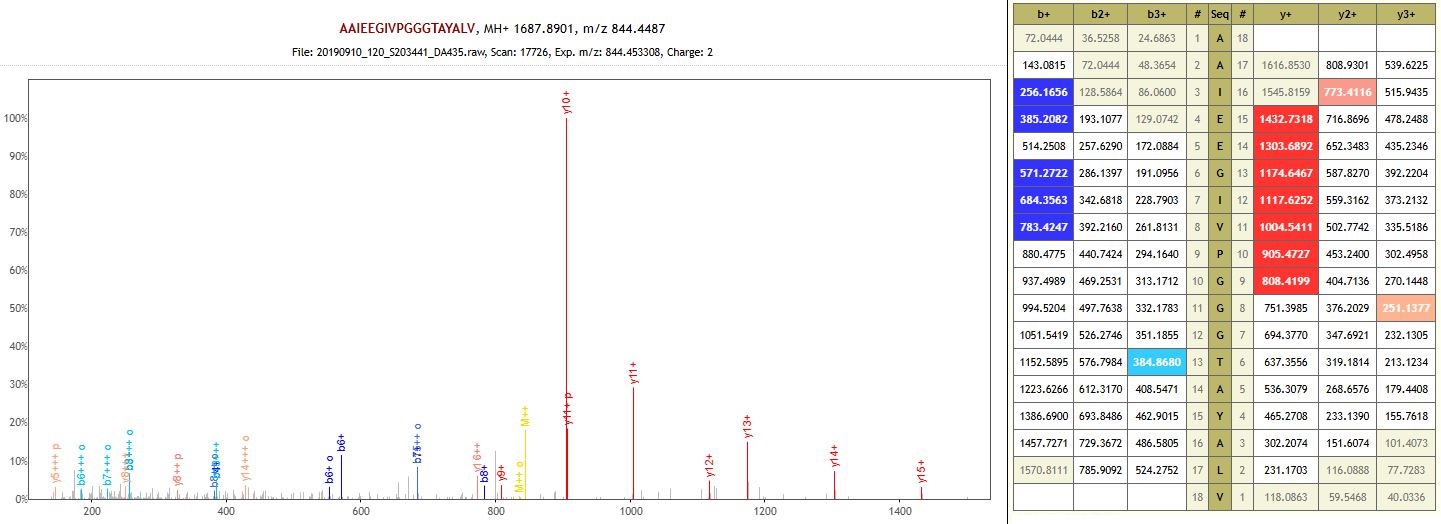

Protein: RuBisCO large subunit-binding protein subunit alpha, chloroplastic (non-specific enzyme)

**DA436 – Khavalynsk 1 – Eneolithic**

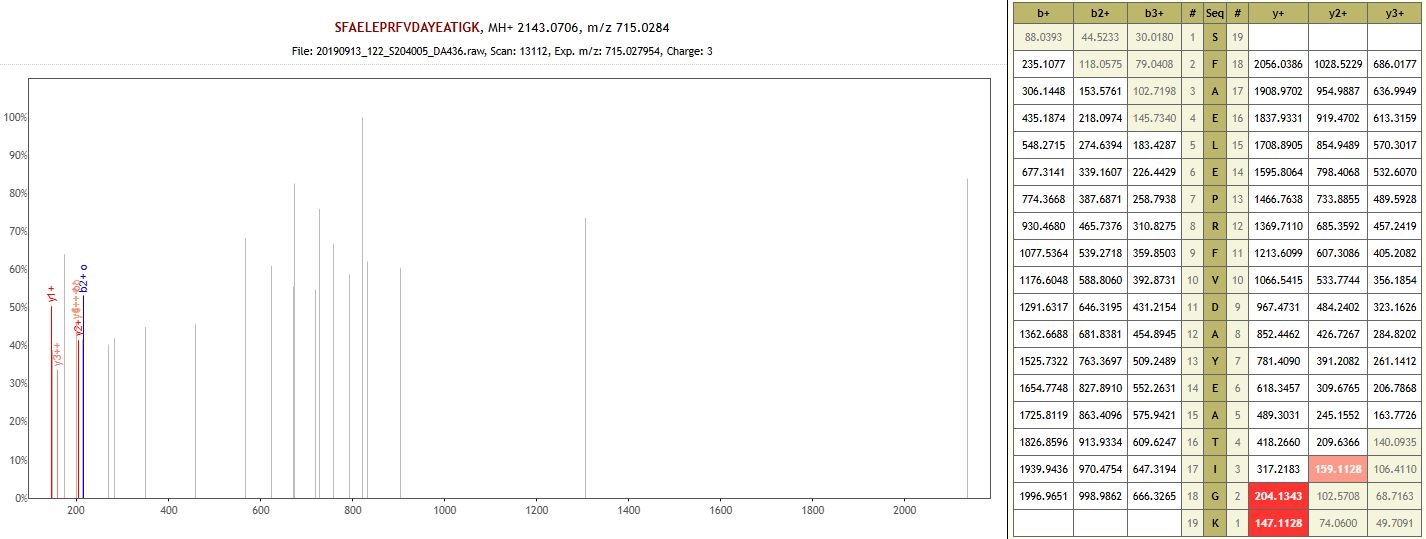

Protein: UDP-glycosyltransferase 73C6-like (non-specific enzyme)

**DA437 – Murziha 2 – Eneolithic**

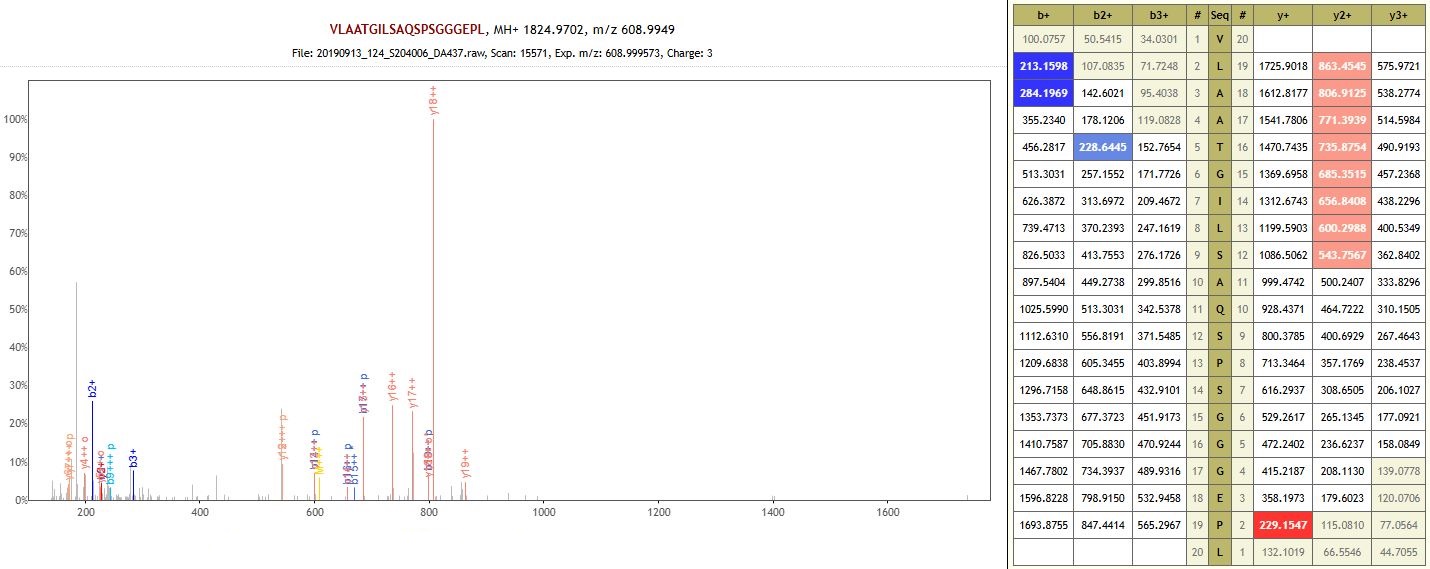

Protein: O-methyltransferase ZRP4-like (auto-enzyme)

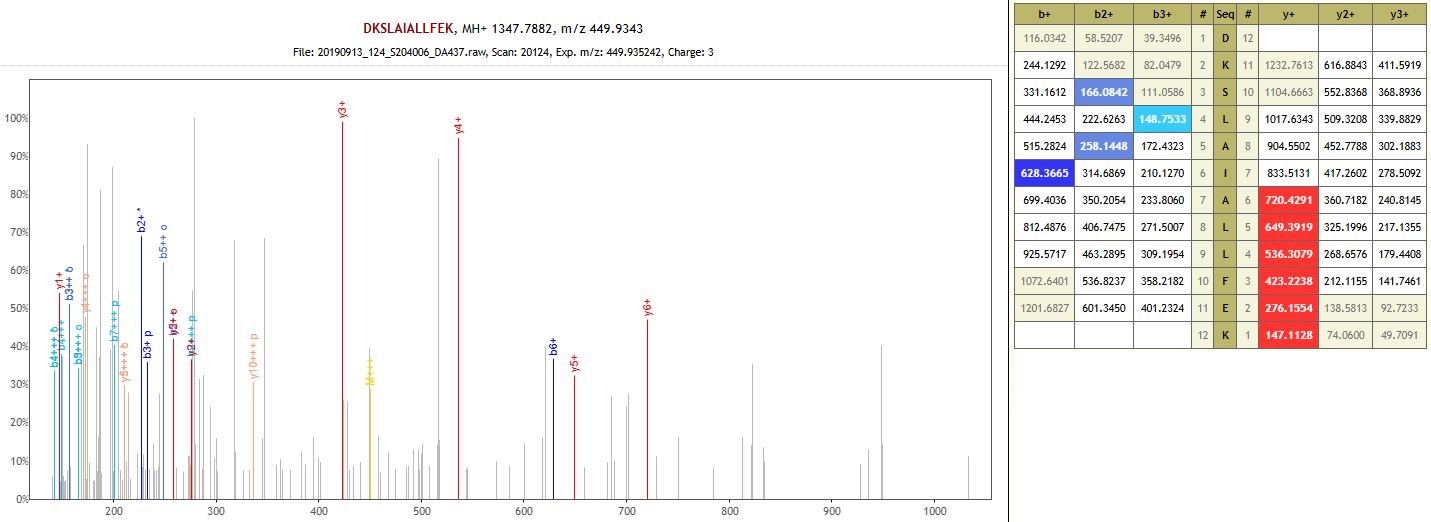

Protein: Monoglyceride lipase (non-specific enzyme)

**DA438 – Potapovka 1 – Middle Bronze Age**

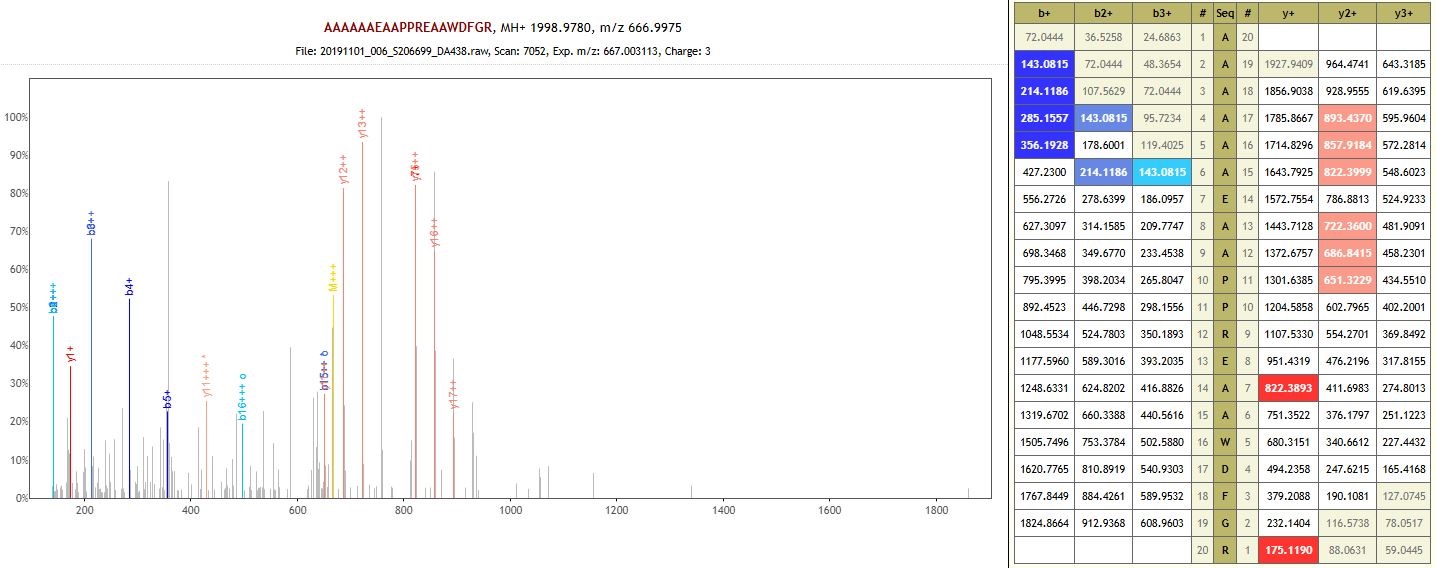

Protein: Uncharacterized protein (non-specific enzyme)

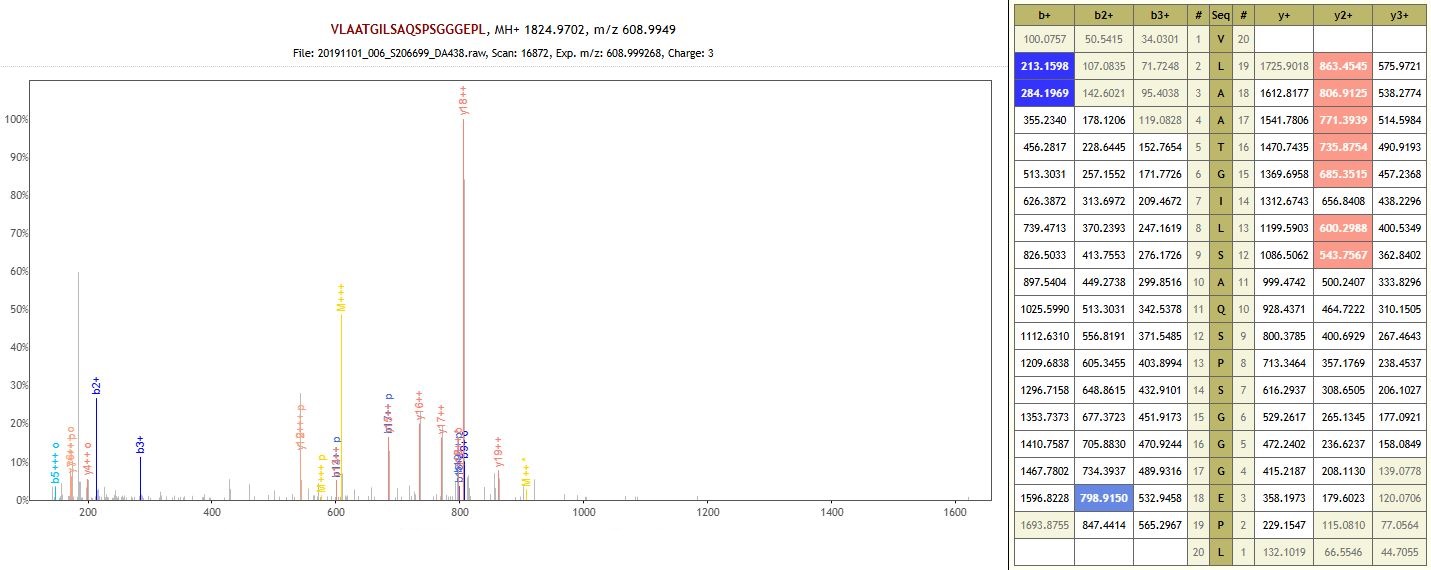

Protein: O-methyltransferase ZRP4-like (non-specific enzyme)

**DA439 – Krasikovskyi 1 – Early Bronze Age**

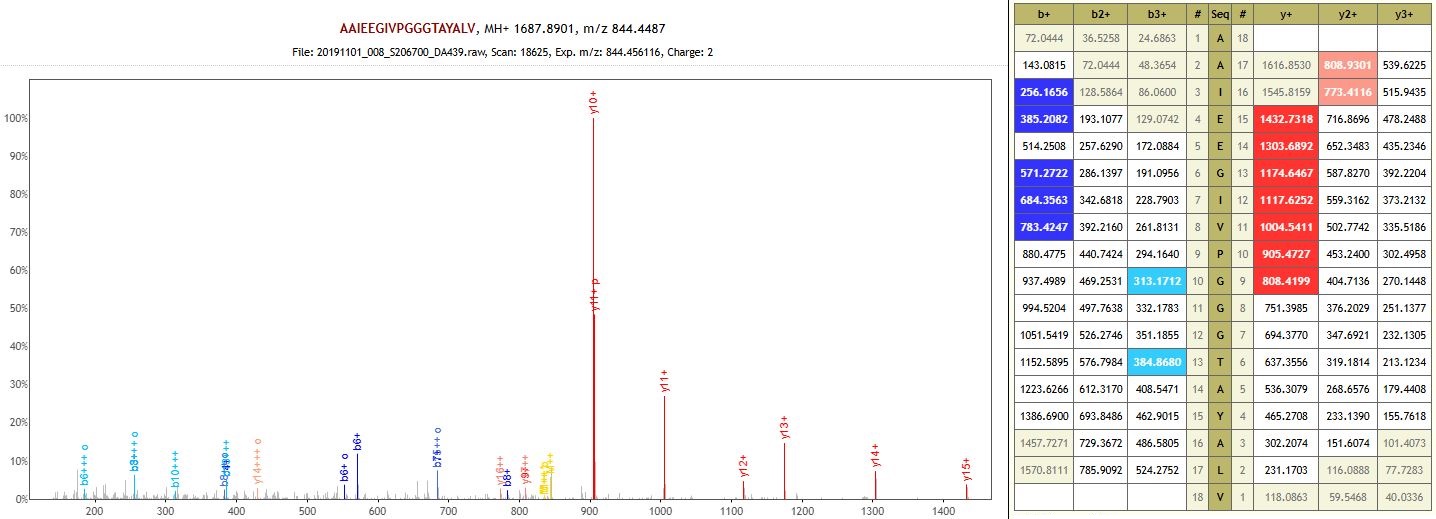

Protein: RuBisCO large subunit-binding protein subunit alpha, chloroplastic (auto-enzyme)

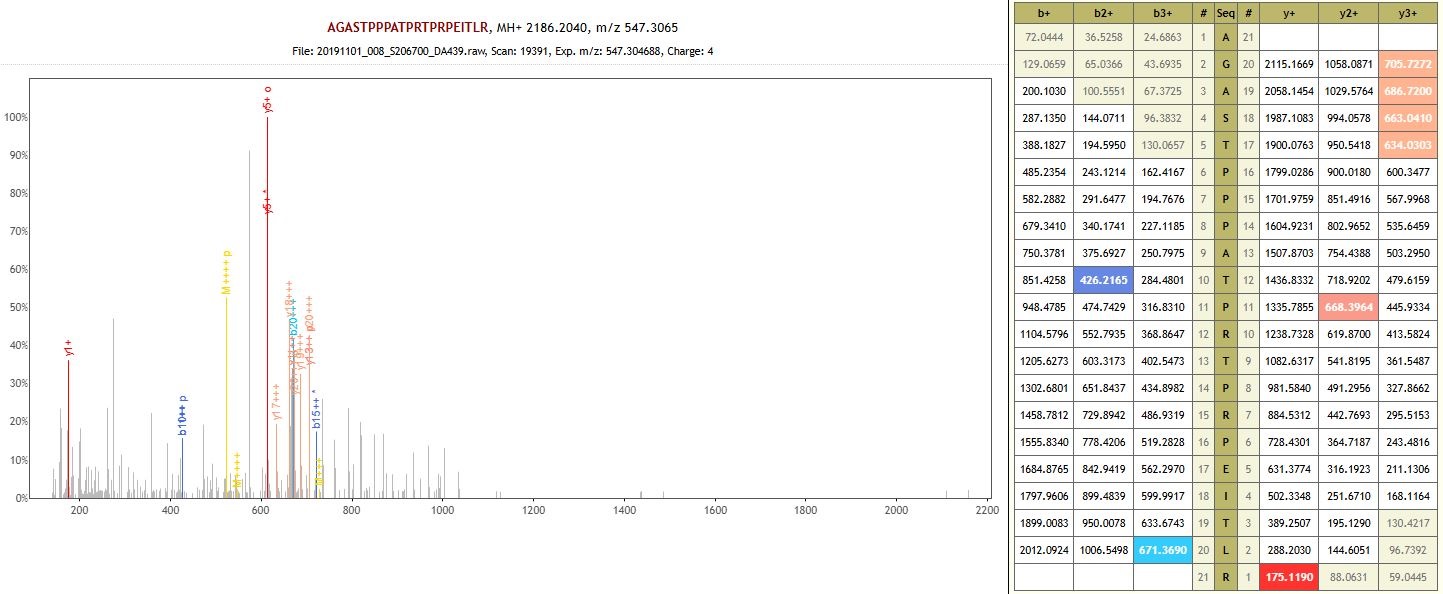

Protein: Protein MIZU-KUSSEI 1-like (auto-enzyme)

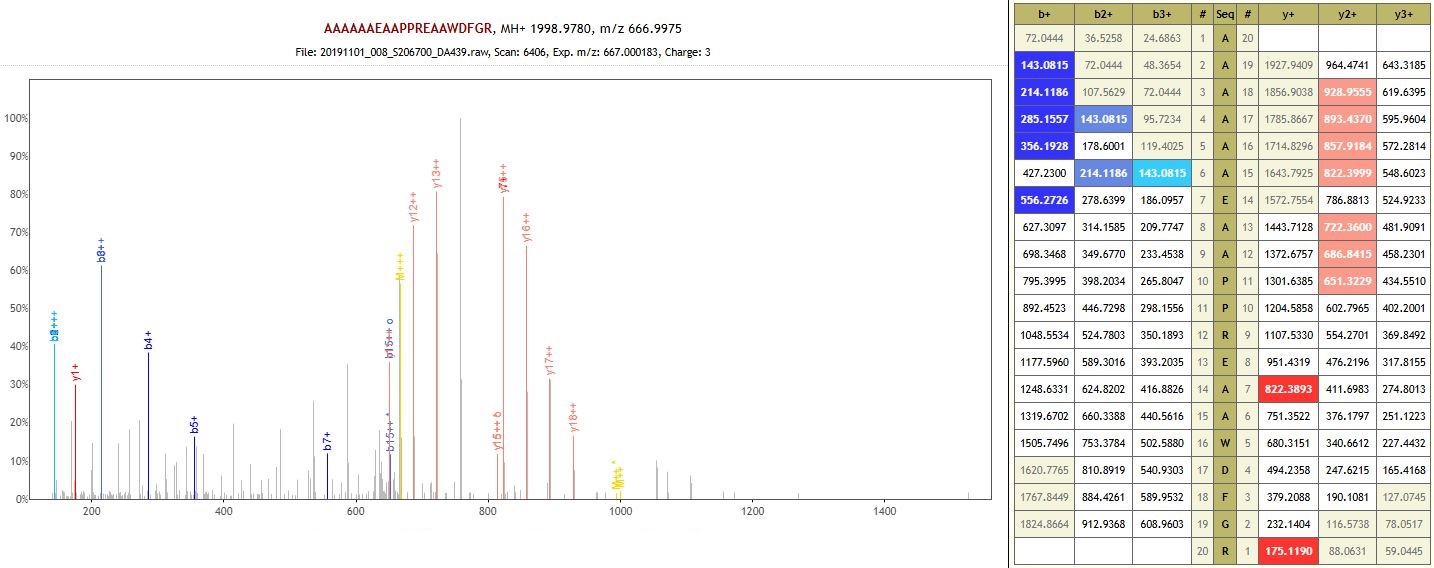

Protein: Uncharacterized protein (non-specific enzyme)

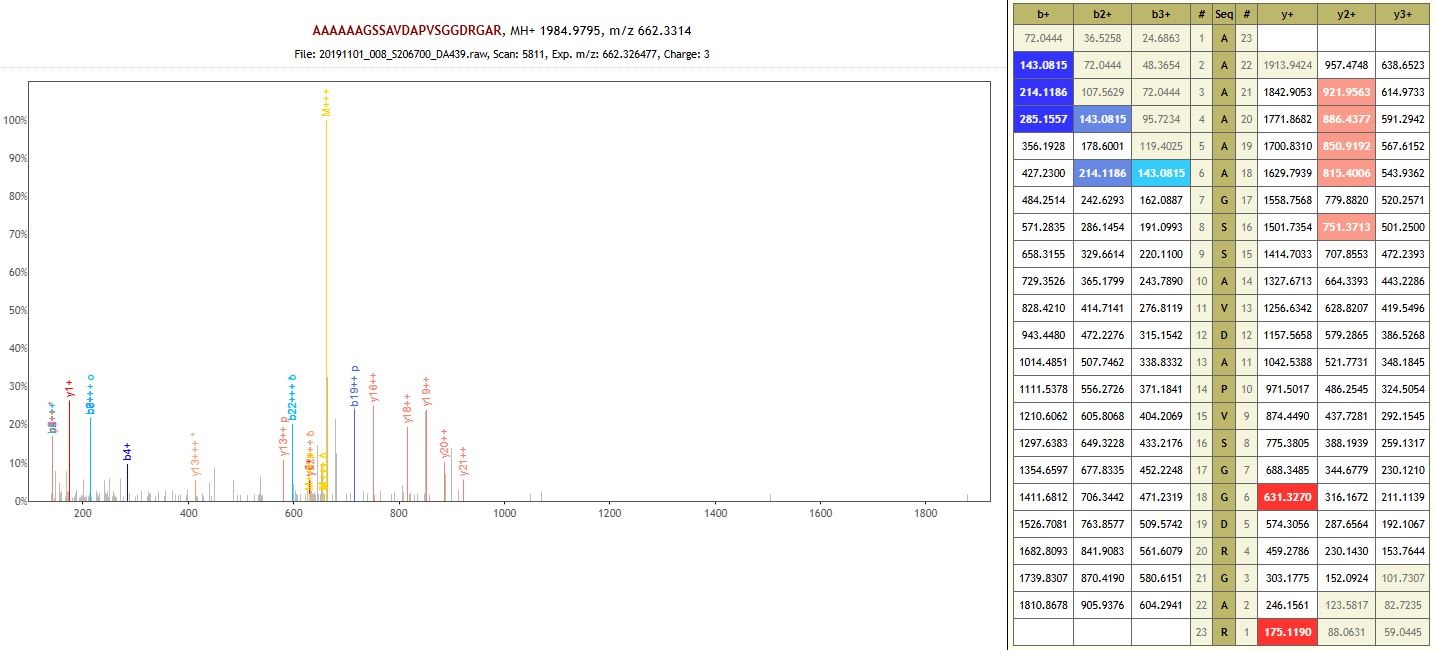

Protein: 3-hydroxyisobutyrate dehydrogenase-like 1, mitochondrial (non-specific enzyme)

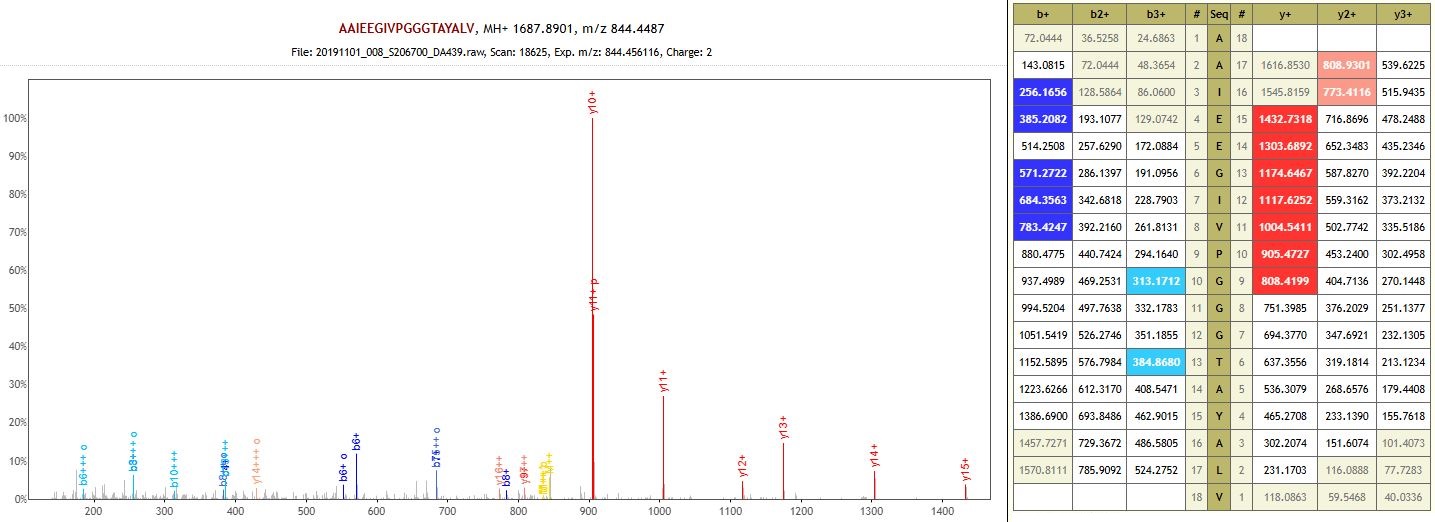

Protein: RuBisCO large subunit-binding protein subunit alpha, chloroplastic (non-specific enzyme)

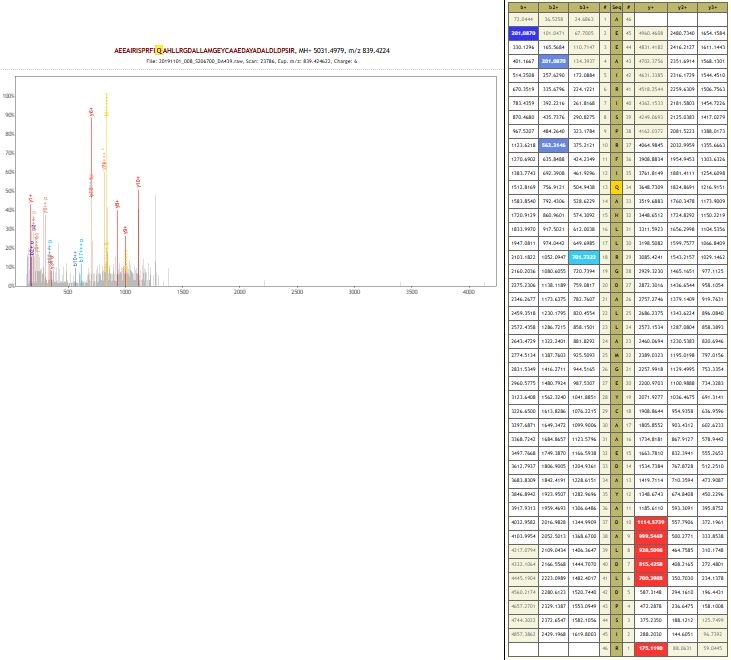

Protein: Small glutamine-rich tetratricopeptide repeat-containing protein 2 (non-specific enzyme)

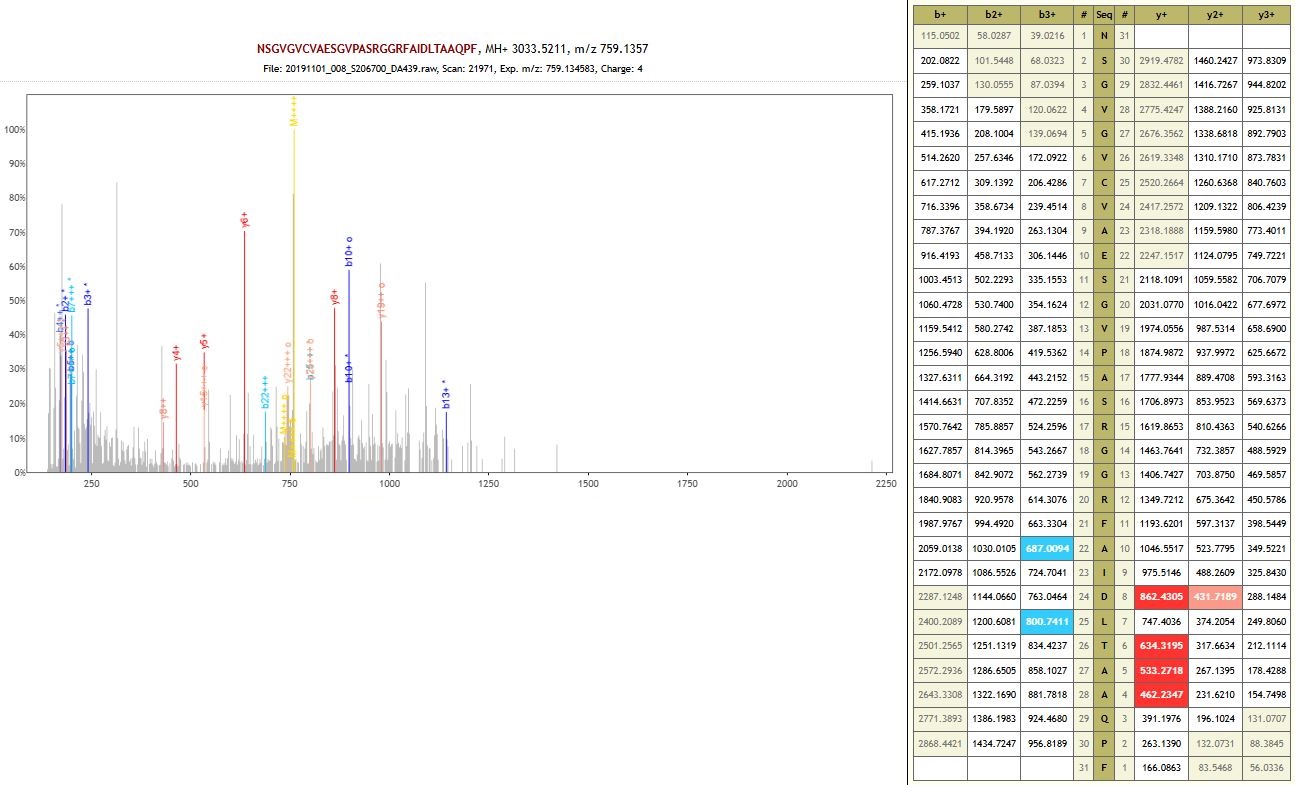

Protein: Formin-like protein 18 (non-specific enzyme)

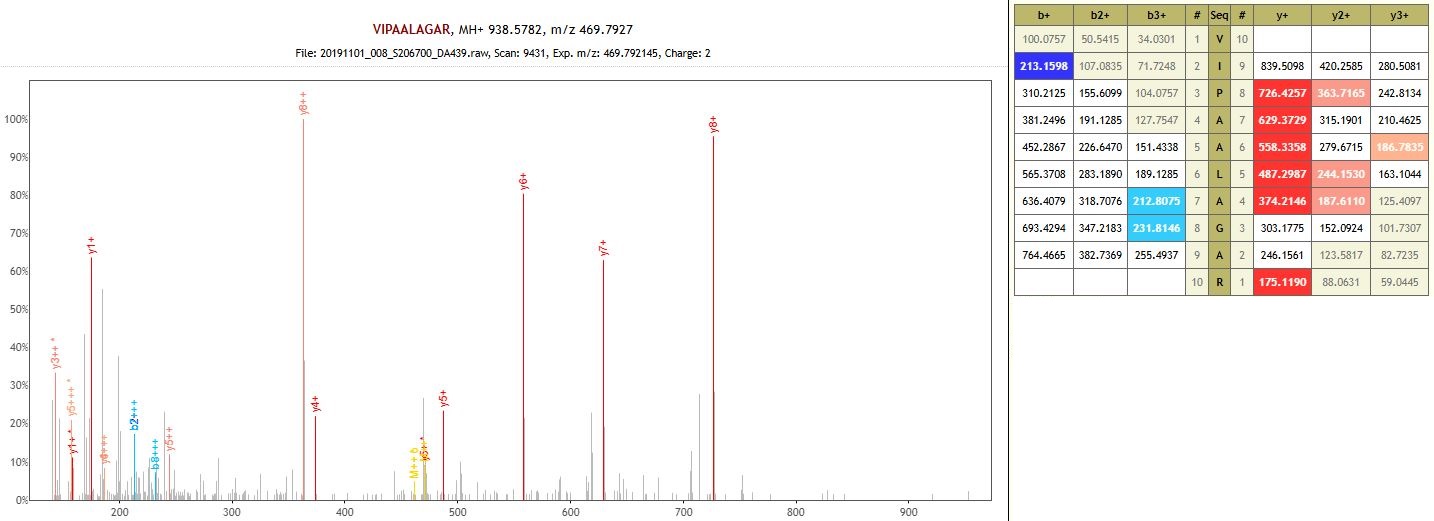

Protein: non-specific serine/threonine protein kinase (non-specific enzyme)

**DA442 – Khavalynsk 2 – Eneolithic**

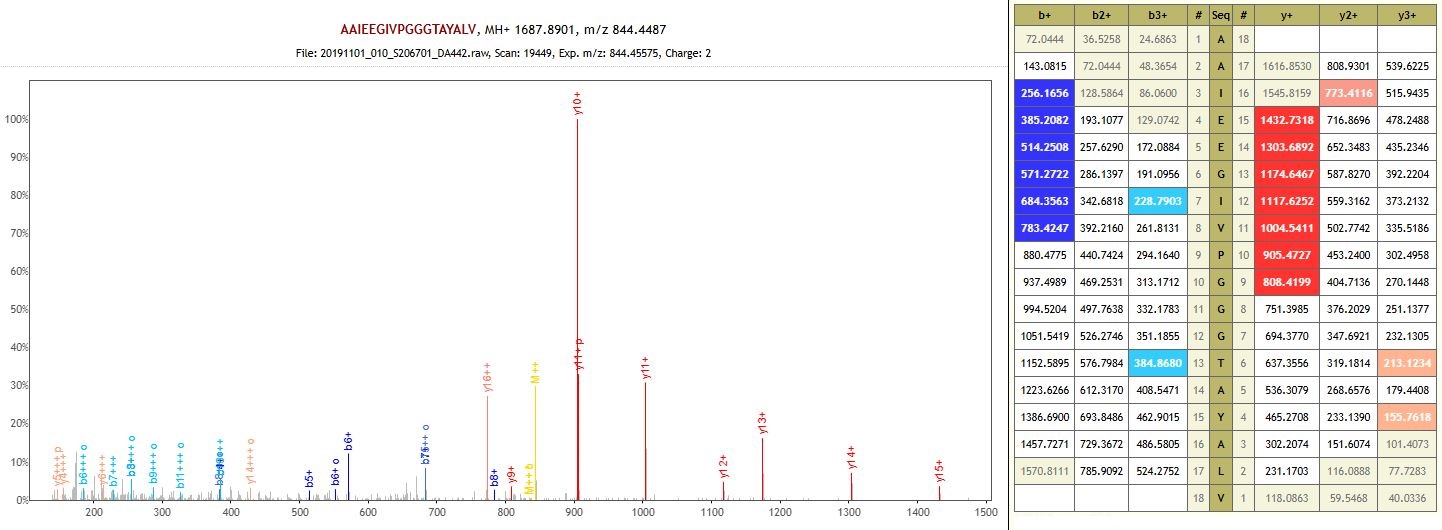

Protein: RuBisCO large subunit-binding protein subunit alpha, chloroplastic (auto-enzyme)

**DA511 – Krasnokholm 3 – Early Bronze Age**

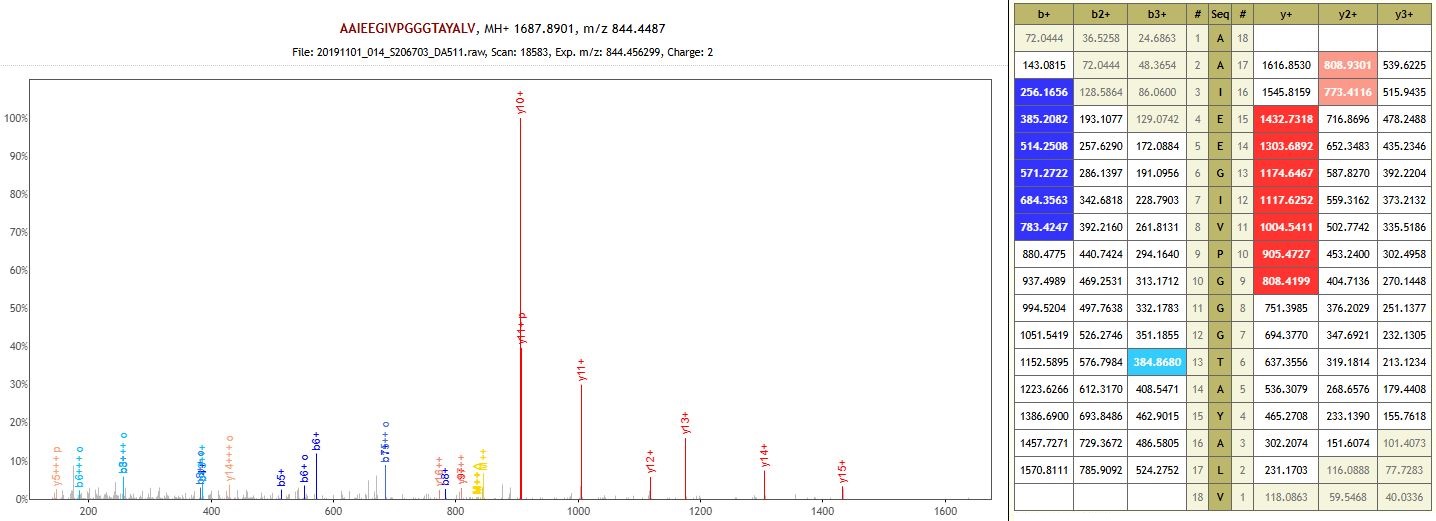

Protein: RuBisCO large subunit-binding protein subunit alpha, chloroplastic (auto-enzyme)

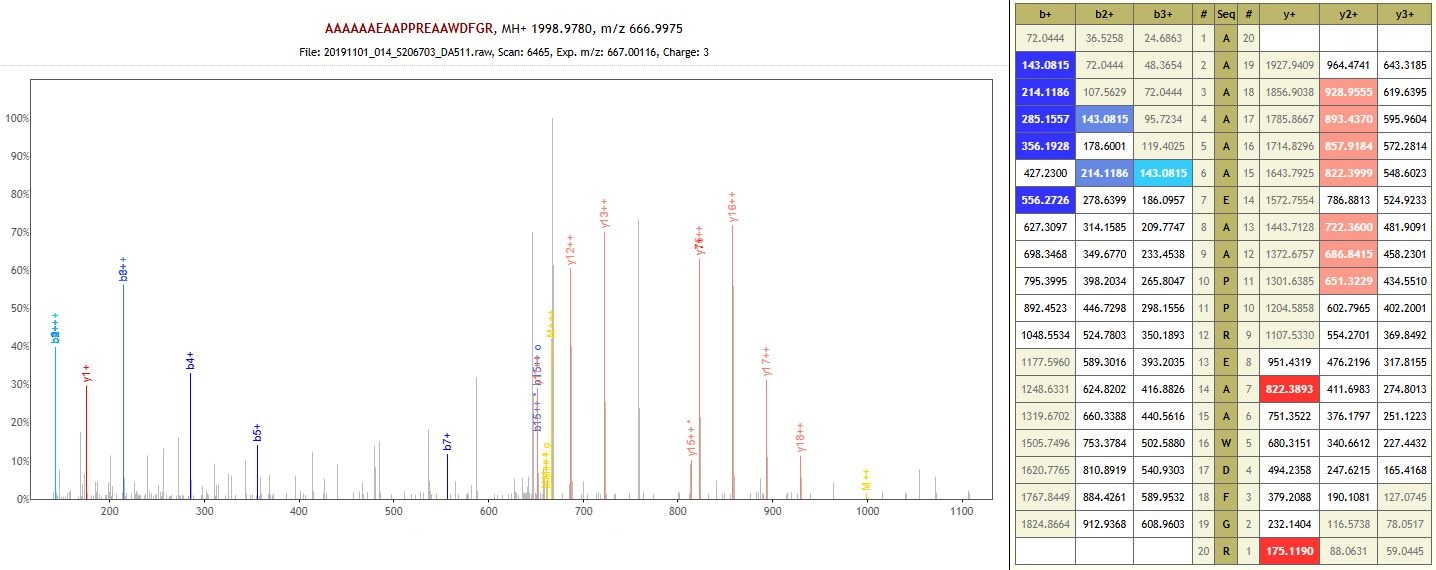

Protein: Uncharacterized protein (non-specific enzyme)

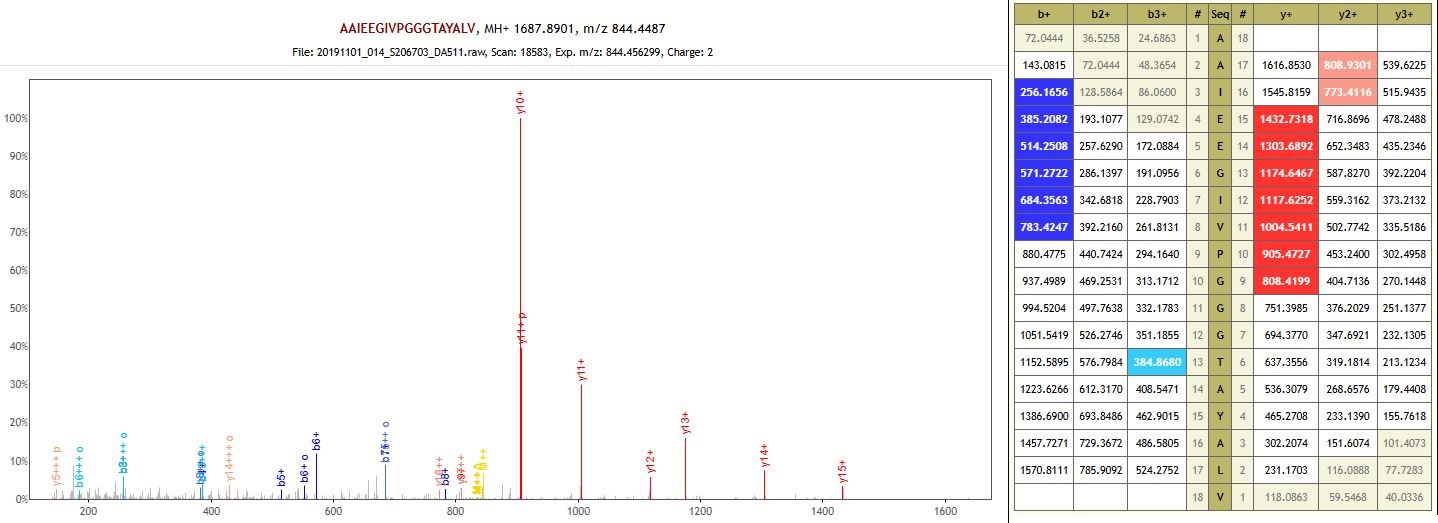

Protein: RuBisCO large subunit-binding protein subunit alpha, chloroplastic (non-specific enzyme)

**DA512 – Nizhnaya Pavlovka 5 – Early Bronze Age**

Protein: Uncharacterized protein (auto-enzyme)

**Z438 – Krivyanskiy 9 – Early Bronze Age**

Protein: Plasma membrane ATPase (non-specific enzyme)

**Z443 – Murziha 2 – Eneolithic**

Protein: Glyceraldehyde-3-phosphate dehydrogenase (non-specific enzyme)

Protein: Glyceraldehyde-3-phosphate dehydrogenase (non-specific enzyme)

**Z447 – Utevka 6 – Middle Bronze Age**

Protein: Glyceraldehyde-3-phosphate dehydrogenase (non-specific enzyme)

Protein: Glyceraldehyde-3-phosphate dehydrogenase (non-specific enzyme)

Protein: Glyceraldehyde-3-phosphate dehydrogenase (non-specific enzyme)

Protein: E3 ubiquitin-protein ligase MARCH2-like (non-specific enzyme)
